# The D4Z4caster DNA methylation signature identifies individuals at epigenetic risk for developing facioscapulohumeral muscular dystrophy (FSHD)

**DOI:** 10.64898/2026.05.26.727947

**Authors:** Takako I. Jones, Brooke Z. Eriksen, Maryam Naveed Farooqi, Taylor Gould, Peter L. Jones, Oliver D. King

## Abstract

**Background:** Facioscapulohumeral muscular dystrophy (FSHD) is caused by epigenetic dysregulation at the chromosome 4q35 D4Z4 repeat array under specific permissive genetic conditions. Due to the complexity, expense, and general inaccessibility of FSHD genetic testing, many individuals displaying characteristic muscle weakness are never genetically confirmed and at-risk relatives cannot get screened. We previously developed a targeted bisulfite sequencing (BSS) protocol using the Sanger method to determine DNA methylation levels at specific D4Z4 loci relevant to distinguishing forms of FSHD from non-FSHD that can be used with DNA isolated from saliva, thereby reducing cost and increasing accessibility compared to traditional D4Z4 deletion testing that uses DNA isolated from blood.

**Methods:** Here, we adapt the D4Z4 BSS protocol to next-generation sequencing (NGS) to increase sequencing depth and further reduce cost, validate both sequencing technologies against several cohorts of genetically defined samples, and introduce the D4Z4caster software for computing DNA methylation signatures with diagnostic utility from raw sequencing data.

**Results:** Both Sanger and NGS BSS methods using D4Z4caster were validated as providing high sensitivity and specificity, with geometric mean of sensitivity and specificity (G-mean) >95% and area-under-the ROC curve (AUC) of 0.99. The NGS method allows for higher throughput and increased read depth, while the Sanger method allows faster processing of individual samples. Importantly, the NGS method could identify FSHD1 cases that are likely mosaic and would otherwise be missed.

**Conclusions:** D4Z4caster methylation signatures can accurately detect contracted FSHD1-permissive chromosome 4q35 alleles, hypomethylation of D4Z4 arrays indicative of FSHD2, and SNPs that are important for diagnostic use. This workflow is amenable to transitioning to clinical settings for an accurate, low-cost FSHD molecular diagnostic test that could be accessible worldwide.

**What is already known on this topic:** Currently accepted genetic diagnostics for FSHD1 are complex and expensive and can mischaracterize certain complex genetic cases. These diagnostics all require high molecular weight genomic DNA typically freshly isolated from blood, highly specialized equipment, and additional testing for FSHD2, making FSHD diagnostics the most expensive among neuromuscular diseases and inaccessible to much of the world. However, the epigenetic status of the 4q35 and 10q26 D4Z4 repeat arrays, as determined by DNA methylation status using our bisulfite sequencing-based protocol, distinguishes genetically FSHD1, FSHD2, and non-FSHD samples. Additionally, since our protocol is PCR-based, it can utilize DNA isolated from multiple sources, including saliva and buccal swabs.

**What this study adds:** This study validates the relevant DNA methylation signatures against several large cohorts of genetically-confirmed FSHD and non-FSHD samples and optimizes the DNA methylation data analysis for the greater accuracy required for diagnostic utility, including the exclusion of nonpathogenic chromosome 10q or 4A166 contractions. In addition, we introduce the D4Z4caster analysis software, which runs in a portable and scalable Docker container, and provides increased quantitative accuracy important for: 1) confirming likely clinical cases of FSHD that do not meet the currently accepted genetic definition of FSHD1 or FSHD2, 2) identifying FSHD1 somatic mosaicism, and 3) potential prognostic applications.

**How this study might affect research, practice or policy:** FSHD1 is genetically defined by a D4Z4 array at the 4q35 locus that is contracted to 1-10 repeat units. However, disease penetrance is influenced by repeat number, epigenetic modifications, and genetic background, causing a misalignment of current genetic diagnosis with clinical diagnosis. This study will improve the accuracy of epigenetic analysis for determining cases of genetic FSHD, help broaden the definition of genetic FSHD to more accurately correspond to clinical FSHD, and allow identification of those at risk for developing clinical FSHD in affected families and in large population studies now being performed and proposed. In addition, it will better inform how an individual’s epigenetic status is interpreted for potential prognostic value. Overall, this methodology is: 1) significantly less expensive than current clinically-approved FSHD diagnostic technologies, 2) more accessible due to compatibility with DNA isolated from multiple sources including saliva, and 3) compatible with the current sequencing equipment and workflow for DNA isolation used in commercial clinical laboratories. Together, these advantages will help move the technology toward becoming an approved molecular diagnostic test for FSHD in the USA, Europe, and countries currently lacking clear access to testing.

## Introduction

Facioscapulohumeral muscular dystrophy (FSHD) is currently considered the third most common muscular dystrophy, with a reported prevalence of 5-12/100,000 (1, 2); however, due to limitations regarding genetic testing, its overall prevalence is likely much higher. The genetic criteria for FSHD are complex and not assessed in typical neuromuscular disease screens using candidate gene panels or whole exome sequencing, and, even when available, FSHD-specific genetic testing is not always conclusive (3–5). Additionally, the high variability in age of onset, progression, presentation, and severity of FSHD symptoms (6, 7) can complicate and delay an accurate clinical diagnosis (4). Many individuals with genetic FSHD are mildly affected or asymptomatic most of their lives and never seek out evaluation by a neurologist or undergo genetic testing (8–14). Since FSHD1 (∼95% of cases) has autosomal dominant inheritance, albeit with incomplete penetrance, and FSHD2 (∼5% of cases) is digenic with dominant inheritance (15, 16), the late onset of clinical FSHD or the presence of asymptomatic FSHD in a family often results in pedigrees with many genetically FSHD family members being differentially affected across several generations (17, 18). Overall, many individuals at risk for developing clinical FSHD later in life or passing it on to the next generation do not have the necessary information for making lifestyle, long term care, or family planning decisions. Thus, there is a critical need for more accurate and accessible molecular diagnostics for confirming clinical FSHD and identifying those at risk for developing the disease or passing it on to their children.

All forms of FSHD are associated with genetic mutations that affect the epigenetic status of the chromosome 4q35 D4Z4 macrosatellite repeat array, leading to the pathogenic increased expression of the *DUX4* (Double homeobox 4) gene from the distal-most repeat unit (RU) within the array (5, 19–22). Genetic FSHD1 (OMIM #158900) is currently defined by deletions that reduce the number of D4Z4 RUs to 1-10, resulting in an epigenetic dysregulation of the contracted array (23–26). FSHD2 is primarily caused by mutations in the *SMCHD1* (Structural Maintenance of Chromosomes flexible Hinge Domain Containing 1) gene (∼80-85% of cases), and more rarely in the *DNMT3B* (DNA Methyltransferase 3 Beta) and *LRIF* genes, all of which are genes encoding epigenetic repressors of the 4q35 D4Z4 array, again leading to epigenetic dysregulation of the locus on both alleles as well as on the chromosome 10q26 D4Z4 array (15, 16, 27). OMIM subdivides “contraction-independent” FSHD into FSHD2 (OMIM **#** 619477) for *SMCHD1* mutations, FSHD3 (OMIM **#**619478) for *LRIF1* mutations, and FSHD4 (OMIM **#**619478) for *DNMT3B* mutations. This nomenclature is also used in ClinVar but is not fully penetrant in the FSHD community, and our use of the term FSHD2 will typically encompass all of them, as in recent guidelines for best practices on genetic diagnosis (28).

There is an additional genetic requirement for all forms of FSHD, located in a subtelomeric sequence distal to the 4q D4Z4 array: a somatic polyadenylation signal (PAS) for *DUX4* that is required for stable expression of *DUX4* mRNA in FSHD (21, 29–31). This PAS in the third exon of *DUX4* is present on roughly half of the chromosome 4s (termed 4qA or 4A) in the human population and enables production of stable full-length *DUX4* mRNA when there is epigenetic dysregulation of the *cis* D4Z4 array; thus, these alleles are considered FSHD permissive. The remaining ∼50% of chromosome 4s (termed 4qB or 4B) lack this somatic PAS and cannot make full-length *DUX4* mRNA; thus, they are considered FSHD nonpermissive, as contraction or epigenetic dysregulation of these nonpermissive 4B D4Z4 arrays does not result in FSHD (32).

Importantly for sequence-based analyses, the region proximal to the 4A exon also exists as an alternate longer allele, termed 4AL (24). In addition, there are major and minor sub-haplotypes for each of 4A, 4AL and 4B. These sub-haplotypes are often referred to by lengths of a simple sequence length polymorphism (SSLP) ∼3kb proximal to the D4Z4 array (e.g., 4A161, 4A166, 4A161L, 4B163), but there can be variation in the distal repeat even among alleles with the same SSLP, arising in part from intra- and inter-chromosomal rearrangements (33). Some 4A166 sub-haplotypes are not typically associated with FSHD even when contracted, despite containing the somatic PAS that characterizes permissive vs non-permissive alleles (29, 33). The reason for this is not yet fully elucidated, but these 4A166 sub-haplotypes can be distinguished based on certain single-nucleotide polymorphisms (SNPs) in the distal repeat (33), which could potentially affect processing or stability of *DUX4* mRNA. Accurate genetic testing for FSHD needs to account for all these complex criteria.

The current genetic testing technologies that are clinically accepted for FSHD1 diagnostics involve measuring the sizes of the relevant intact 4q35 D4Z4 arrays combined with distal haplotyping for FSHD-permissive alleles (3, 34–39). Not surprisingly, these complex approaches require high quality high molecular weight (HMW) genomic DNA (gDNA) freshly isolated from peripheral blood mononuclear cells (PBMCs) and run on specialized equipment. These tests are expensive and are generally not accessible to much of the world. FSHD2 testing is, on the surface, simpler and more amenable to candidate gene sequencing and/or DNA methylation analysis (4, 15, 40), but these typically do not account for the additional 4q35 D4Z4 array requirements for FSHD2: having an FSHD-permissive 4A haplotype that – despite the continued use of the historical descriptor “contraction-independent” to distinguish FSHD2 from FSHD1 – is usually on a “semi-short” array of 11-20 RU. Alternatively, since both FSHD1 and FSHD2 mutations lead to increased expression of *DUX4* from the distal-most RU of epigenetically dysregulated 4q-type D4Z4 arrays only in the context of an FSHD-permissive distal PAS, we proposed that a DNA sequence-based epigenetic analysis specific for the distal-most D4Z4 RU could address these issues and potentially revolutionize FSHD genetic testing. Therefore, we developed an epigenetic approach using bisulfite genomic sequencing (BSS) to identify FSHD1, FSHD2, and non-FSHD DNA methylation profiles for the FSHD-associated 4q35 D4Z4 array and related chromosome 10q26 D4Z4 arrays (Figure 1). We demonstrated that this approach produced data diagnostic for each indication over a limited number of samples (13, 41, 42). Importantly, this approach analyzes the epigenetic status of FSHD-permissive alleles while also assessing key sequence variations in the region. Here, we further validate this technique against several large genetically-defined cohorts and improve the analysis workflow by transitioning to a next-generation sequencing (NGS) approach that allows multiplexed processing of samples and increased sequencing depth, coupled with new bioinformatic software, called D4Z4caster, that computes DNA-methylation-based scores that are diagnostic for genetic FSHD.

**Figure 1:**
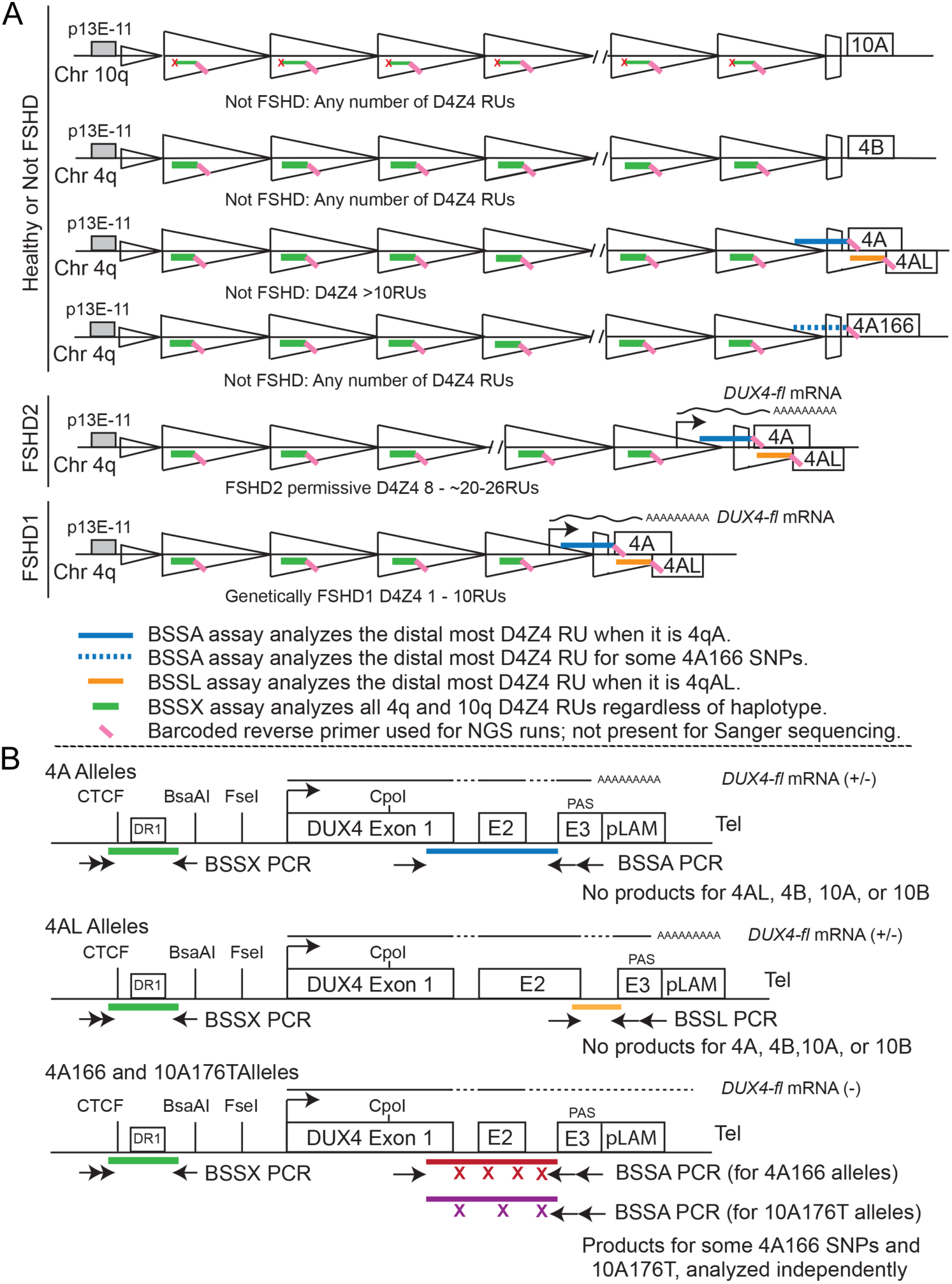
Regions analyzed by targeted BS-PCRs for FSHD1, FSHD2, and non-FSHD chromosomes. A: The BSSA assay is specific to the distal repeat unit (RU) of 4qA alleles, the BSSL assay is specific to the distal RU of 4qAL alleles, and the BSSX assay picks up all 4q and many 10q D4Z4 RUs regardless of haplotype. B: The locations of the BS-PCR primers are shown on the distal RU. The *DUX4* gene structure, including the exon 3 PAS, the FseI, BsaAI, and CpoI sites used for methylation-sensitive restriction enzyme analysis, the DR1 region used for FSHD2 BSSX methylation analysis, and the putative CTCF binding site are shown for reference, as detailed in our earlier study from which these schematics are adapted (Ref 41).

## Methods

### Human Subjects

This project was approved by the University of Nevada, Reno Institutional Review Board (Study ID #1316095, approved on 31 July 2019). All identifiable participants provided signed informed consent. For the UNR IRB saliva study, saliva collection kits (Oragene Discover ORG-500, DNA Genotek) or cheek swab collection kits (Oragene Discover ORG-575, DNA Genotek) were sent to control non-FSHD and FSHD participants around the world by mail or courier. Sample self-collection was performed at home and kits were returned by mail or courier. Most participants had confirmed genetic testing for FSHD by other entities, documented in Table S1, although the reports were not always provided to us. Participants with self-reported status and report data not provided (DNP) are distinguished from those for whom the genetic status was not determined by testing (ND). De-identified samples provided by other institutions and studies were declared exempt from UNR IRB approval (Exemption Category #4). De-identified myoblast and fibroblast cell culture samples were obtained from the University of Rochester Medical Center and the Senator Paul D. Wellstone Muscular Dystrophy Specialized Research Center (MDSRC) at the University of Iowa. Additional FSHD BSS data collected by the Jones lab and included in this extended analysis was from previously published cohorts (13, 40–42).

The FSHD genetic status of subjects and cell lines was confirmed by prior genetic testing with D4Z4 array lengths and/or *SMCHD1* mutations provided when available (Table S1). FSHD1 was defined as having at least one FSHD-permissive chromosome 4q D4Z4 array with 1-10 D4Z4 RUs if using optical genome mapping (OGM) or <38kb in length if using *Eco*RI/*Bln*I digestion and Southern blotting. FSHD2 was defined as having at least one FSHD-permissive chromosome 4qA D4Z4 array (limited to 8-20 RU if this information was provided) and a mutation in the *SMCHD1* gene reported to be pathogenic in the ClinVar (43) (if mutation information was provided). None of the subjects in this study were reported to have pathogenic mutations in *LRIF1* or *DNMT3B,* which is not itself surprising as these are much rarer causes of FSHD2, though because they are also more recently-discovered causes they may not have been sequenced or examined in all cases. It is possible that some individuals listed as genetically FSHD2 for whom reports were not provided may have variants of uncertain significance (VUS) in *SMCHD1* that were interpreted by clinicians as being likely pathogenic when, in fact, they are benign, and that some may not have semi-short 8-20 RU 4qA or 4qAL D4Z4 arrays, which, although found in most clinically-manifesting cases of FSHD2, may not be a strict requirement (28); one recent study reported shortest permissive arrays of up to 35 RU (44). Individuals with pathogenic *SMCHD1* mutations and 4qA or 4qAL D4Z4 arrays of 1-10 RU are regarded as clinically FSHD1 + FSHD2.

### Genomic DNA (gDNA) isolation, haplotyping, and simple sequence length polymorphism (SSLP) analysis

gDNAs were isolated from cultured cells, peripheral blood mononuclear cells (PBMCs), or saliva samples using the Wizard Genomic DNA Purification Kit (Promega). Distal A/B haplotyping and proximal SSLP analysis were performed on all gDNAs as described previously (13, 21, 41). Purified gDNAs (1-1.5µg per sample) were then bisulfite converted as per manufacturer’s instructions (EpiTect Bisulfite Kit, Qiagen) as described previously (13, 41) and used for BS-PCR.

### BS-PCR for Sanger sequencing protocol

DNA methylation analysis was performed using the targeted BSS assay for 4qA (BSSA), 4qL (BSSL), and *DUX4* 5’ region (BSSX) as described previously (13, 41, 45). The BSS assay for 4qB alleles (BSSB) was performed as described previously (46). PCR products were TA-cloned using the pGEM-T Easy kit (Promega) and Sanger sequencing (Eton Biosciences) was performed from bacterial colonies isolated directly from agar plates.

### BS-PCR for NGS protocol

DNA methylation analysis was performed using the described BSS assay with the addition of barcode (BC) fused primers unique to each subject but common to all three BSS products at the nested PCR step (detailed below) to allow multiplexed sequencing. The EpiTect Bisulfite Kit (Qiagen) was used for bisulfite conversion of gDNA (1.5 ug) following the manufacturer’s instructions. The initial BS-PCR to amplify 4qA, 4qAL, and the *DUX4* 5’ regions were performed as described previously (13, 41, 45). A nested PCR using 10% of the initial BS-PCR product was performed with fusion primers 1438F-P1 and 3626R-BC-A for BSSA, 4ALF-P1 and 3626R-BC-A for BSSL, or 475F-P1 and 1036R-BC-A for *DUX4* 5’ regions (Table S2). Each forward primer includes a P1 adapter. Ion Express barcode sequences are available through Ion Torrent Server (ThermoFisher.com). All nested PCRs used GoTaq Hot Start Polymerase (Promega) using the following PCR conditions: 4qA- 94°C for 2 min, 5 cycles of 94°C for 15 sec, 58°C for 15 sec, 72°C for 45 sec, 20 cycles of 94°C for 15 sec, 65°C for 15 sec, 72°C for 45 sec, followed by a final extension of 72°C for 10 min; 4qAL- 94°C for 2 min, 5 cycles of 94°C for 15 sec, 58°C for 15 sec, 72°C for 35 sec, 20 cycles of 94°C for 15 sec, 65°C for 15 sec, 72°C for 35 sec, followed by a final extension of 72°C for 10 min; *DUX4* 5’- 94°C for 2 min, 5 cycles of 94°C for 15 sec, 60°C for 15 sec, 72°C for 45 sec, 20 cycles of 94°C for 15 sec, 67°C for 15 sec, 72°C for 45 sec, followed by a final extension of 72°C for 10 min. All BS-PCR products were run on agarose gel, confirmed for expected product size, and extracted using NucleoSpin Gel & PCR Clean-Up kit (Takara). The concentration of purified BS-PCR products was checked using 1x Qubit dsDNA HS kit (Thermo Fisher Scientific) before pooling 0.2 pmol of 4qA and *DUX4* 5’ fragments and 0.1 pmol of 4qAL fragments per sample in 50 µl. These per-individual samples were further cleaned using 0.9x volume of AMPure XP magnetic beads (Beckman Coulter) and eluted to have 200-1000bp product at 4166 pM concentration per manufacturer’s instructions. Each per-individual sample was then diluted 1:13.9 to be 300pM, and samples for multiple individuals (each with a distinct BC) were pooled evenly per number of fragments in a final volume of 100µL (300pM from each library). The library was then diluted to be 30pM with low TE (10mM Tris, 0.1mM EDTA), and 50 µL was denatured at 65°C for 5 min, then placed on ice for several minutes until loading on the Ion Chef (Thermo Fisher Scientific).

### Next-Generation Sequencing (NGS)

NGS was performed using the GeneStudio S5 System (Thermo Fisher Scientific). Samples were sequenced in a total of 51 runs each with up to 70 multiplexed samples, including many samples that are not from the studies discussed herein (Figure 2). Before running the Ion Chef, a run plan was created in the Torrent Suite Software using the following settings: Research Application- DNA, Target Technique- AmpliSeq DNA, Ion Reporter- None, Flows- 1350. The Ion Chef System was loaded using the Ion 530 Chip (Thermo Fisher Scientific) and the Ion 520 & Ion 530 ExT Kit-Chef (Thermo Fisher Scientific) as directed in the Ion 520 & Ion 530 ExT Kit-Chef User Guide (Pub.No. MAN0015805 E.0). The Ion GeneStudio S5 Semiconductor Sequencer was initialized, and the chip was run as directed in the user guide. The Torrent Suite Software demultiplexes reads from different samples based on barcodes to generate an unaligned BAM (uBAM) file for each sample. Following the sequencing run, these uBAM files were analyzed using the D4Z4caster program.

**Figure 2:**
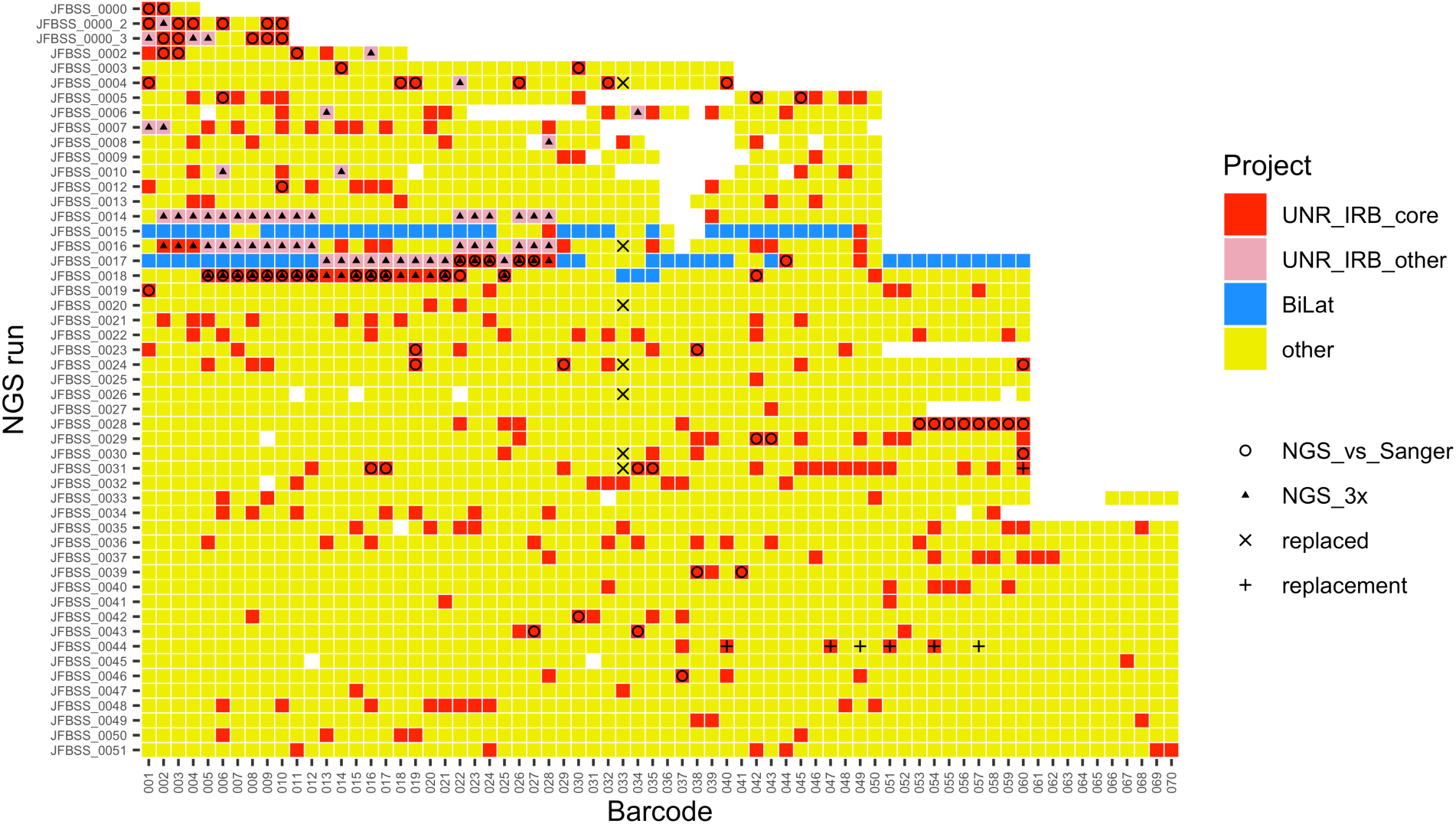
Overview of samples analyzed by NGS. In the grid, rows indicate the NGS run and columns indicate the barcode (001 to 070) used for multiplexing samples within an NGS run. Colors and symbols denote the sub-studies that the samples were used in. UNR IRB “core” samples (red; n=308) are from distinct subjects, to avoid double-counting in evaluating FSHD vs control predictions. Some of these samples were also included in the NGS_vs_Sanger (circles; n = 76), and/or the NGS_3x validation sub-studies; the latter consisted of 3 technical replicates from the same subjects (triangles; n = 27 red + 2*27 pink). Samples from the BiLat study (yellow; n = 71) are from muscle biopsies of subjects with FSHD (n = 32), with paired left and right biopsies for all but a few subjects; in some cases, BSSX assays were done in different runs from BSSA/BSSL assays. Samples not used in any of these studies (“other”) are shown in gray, and barcodes not used in an NGS run are shown in white. Due to a technical problem with barcode 033 in some of the earlier NGS runs, some samples were re-rerun (with x and + symbols denoting the replaced and replacement samples, respectively). The studies discussed in this paper also include 325 samples analyzed by Sanger sequencing. This includes 76 that were also analyzed with NGS for the NGS_vs_Sanger comparison, and for these subjects the Sanger data was excluded for the FSHD vs control comparisons in the UNR IRB study to avoid double-counting subjects.

### D4Z4caster

The D4Z4caster program (Figure 3) provides an end-to-end workflow for analyzing a directory of uBAM files and a sample sheet generated by the NGS workflow described above, and can be customized for different projects (e.g., different conventions for uBAM file names and sample sheet columns). FASTQ or FASTA files can also be used for input, and the latter allows D4Z4caster to be used for analyzing reads from Sanger sequencing. D4Z4caster uses a modified version of Bismark (47) for bisulfite-aware alignment of bisulfite converted sequence reads to a custom set of reference sequences. Here the reference sequences consist of prototype amplicons for the 4A (specifically 4A161, the most common permissive haplotype), 4AL, 4B, and BSSX PCR products along with “decoy” 4A sequences based on 4A166 and 10A176T reference sequences: these haplotypes have >99% homology to 4A161 in the region being amplified, so may in some cases by amplified by the BSSA assay even with mismatches to the PCR primers (particularly in the absence of 4A alleles with no mismatches); including them as references allows Bismark to assign each read to only the best-matching haplotype. The specific 4A166 reference sequence used is from the distal repeat of a 4A166 sub-haplotype that is typically not pathogenic even when contracted; other 4A166 sub-haplotypes may have distal repeats that preferentially map to the 4A161 reference sequence. We also included a 4A161x55 decoy reference that differs from the 4A161 reference at a single base in the 55th CpG, since this was commonly observed in reads even in the absence of the small number of other SNPs that distinguish the 4A166 and 10A176T prototypes from the 4A161 prototype (Table S3). Reads from the BSSA assay for rarer permissive haplotypes such as 4A159 and 4A168 typically mapped to the 4A161 reference sequence although reads for some 4A159 haplotypes can map to 4A161x55 instead.

**Figure 3:**
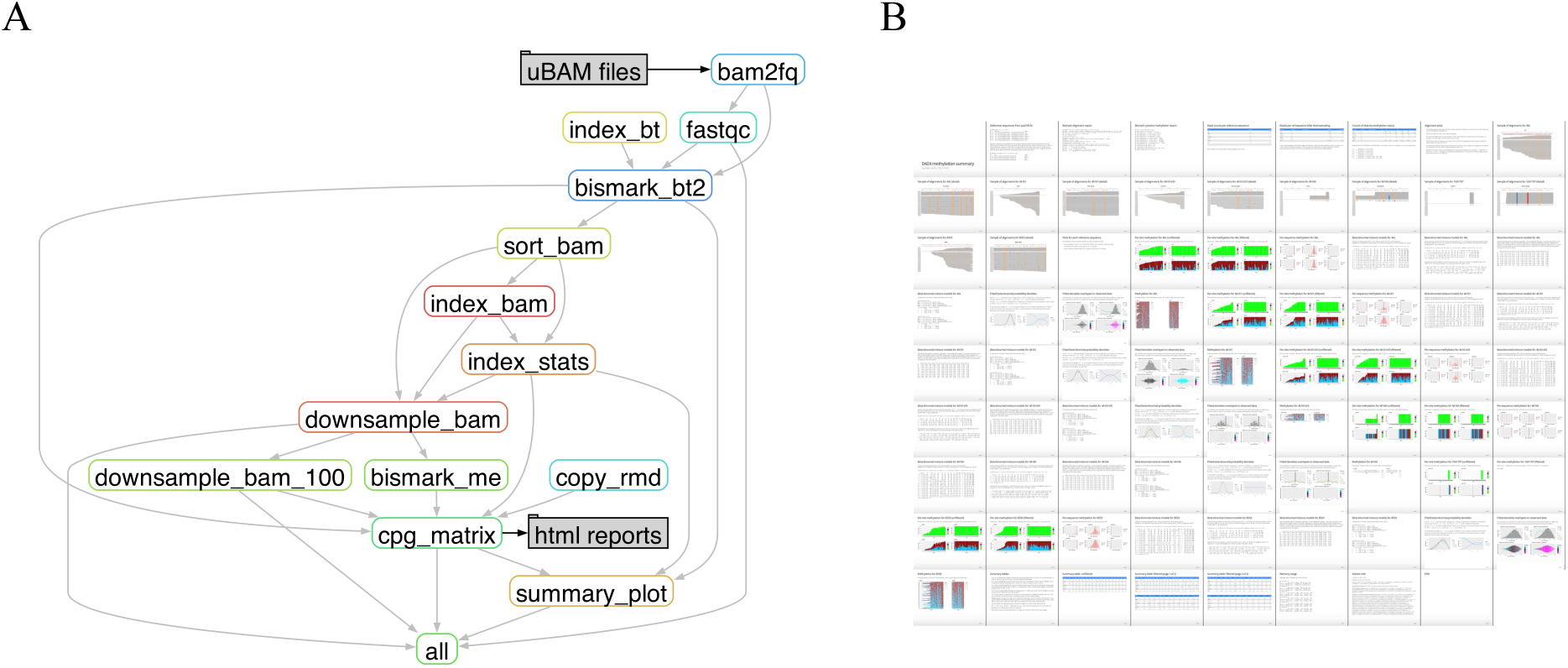
Overview of D4Z4caster software. **A:** D4Z4caster uses a Snakemake workflow run in a Docker container for scalable, portable, and reproducible analyses. Input is a folder of unaligned BAM (uBAM) or FASTQ or FASTA files for one or more samples, which are processed as a batch. The rule-graph of sub-processes is shown. Reads are mapped using a modified version of Bismark with Bowtie2 (bismark_bt2 step). BAMs with aligned reads are down-sampled to retain at most 10000 reads per reference sequence, from which methylation calls are extracted using Bismark (bismark_me step). Per-read counts of methylated CpG calls are modeled with a mixture of beta-binomial distributions using custom R code (cpg_matrix step). BAMs are further down-sampled to retain at most 100 reads per reference sequence only for the read-alignment pileup tracks in the output reports. Output includes a summary plot for all samples in the batch and a detailed report for each sample in the batch. **B:** An example of the output report for a single sample, in HTML slide-show format. This show thumbnails of all slides; re-arranged and slightly modified version of figures from several of the slides are shown in Figure 4.

Bismark methylation extractor is used to export tables of cytosine methylation calls for each read at each CpG site in the reference sequence to which the read mapped. D4Z4caster’s modified version of Bismark sets the context of cystines as “unknown” in reads that do not have the expected guanine in the second position of CpGs in the reference sequences they are mapped to, which can arise from sequencing errors, from genomic variation in targeted haplotypes, or from sequencing of non-targeted 4q haplotypes or homologous 10q sequences. Setting the context of such sites as “unknown” rather than “CpG” makes the mismatch evident in methylation grid plots. The table of methylation calls is processed by custom R code, including R Markdown code that generates reports in html format (with options for pdf or Word docx formats). These reports include methylation plots and tables of summary statistics including means, quartiles (Q1, Q2, Q3) and other percentiles (pct03, pct05, pct10, pct90, and pct95) of mean per-read methylation level for each reference sequence. Notably, this R code also attempts to disentangle the contributions of multiple alleles with the same reference haplotype (e.g., two 4A161 or two 4AL alleles), by fitting the methylation data with beta-binomial mixture models (Figures 4 and 5). This was an innovation in our earlier work (13), and the general analysis framework in D4Z4caster follows that approach but with the following modifications and extensions:

1. The earlier work used a fully Bayesian approach with Markov Chain Monte Carlo (MCMC) used for inference. As there were typically only 10-20 cloned Sanger sequences per assay, priors on parameters played an important role in stabilizing parameter estimates. For our new NGS framework, which typically has thousands of reads for each reference haplotype that occurs in an individual, the prior has much less influence and we instead use Maximum Likelihood Estimation (MLE) to fit the parameters of the mixture models.
2. The earlier work considered only the case of at most two alleles with the same reference haplotype (4A161, 4AL, etc.), and for BSSA and BSSL assays, which target just the terminal D4Z4 repeat, the mixture model assigned each read an equal prior probability of arising from each allele of the same haplotype. (Reads for BSSX assays can also arise from internal D4Z4 repeats, so are expected to occur in proportion to the total number of RUs in the alleles.) Here we extend the mixture models to allow for up to three alleles per reference sequence, as can occur due to translocations between chr4 and chr10 (33) or from having two D4Z4 arrays in *cis* on the same chromosome (48). We also consider both equal and unequal mixing proportions of the alleles. For the latter, these proportions are fit using MLE simultaneously with the beta-binomial parameters (Figure 5). This can be useful for detecting and quantifying mosaicism, as is often seen in *de novo* FSHD cases, in which only a small proportion of 4qA alleles may have a contraction (49, 50). Mixture model names are of the form **e***k* and **u***k*, where **e** and **u** denote **e**qual and **u**nequal (technically just not-necessarily-equal) mixing proportions, and *k* = 1, 2, 3 denotes the number of components in the mixture model (with e1 = u1 denoting a fit to a single beta-binomial distribution). Means *m* = α/(α + β) and concentrations *M* = α + β of the beta distributions for the *k* components of a mixture model are denoted by suffixes m*j* and M*j* (for *j* from 1 to *k*), for example e2.m1 and e2.m2, with components ordered so that e2.m1 ≤ e2.m2.
3. Particularly for Sanger data in which there are relatively few reads there is a risk of overfitting data using MLE, since more complex models (such as models with two vs one mixture components or models with fitted vs fixed mixing proportions) will typically fit the data better than simpler models, giving a higher log-likelihood score. To combat this, we also compute information-theoretic Bayesian Information Criteria (BIC) scores that penalize more complex models, compute weights for models based on these scores (with weights proportional to exp(–BIC/2)) and use these weights for model averaging (51). This strategy can be used to shrink the estimated mean for each component of a two-allele model toward the estimated mean for a one-allele model, by an amount that depends on the difference in BIC scores.

**Figure 4:**
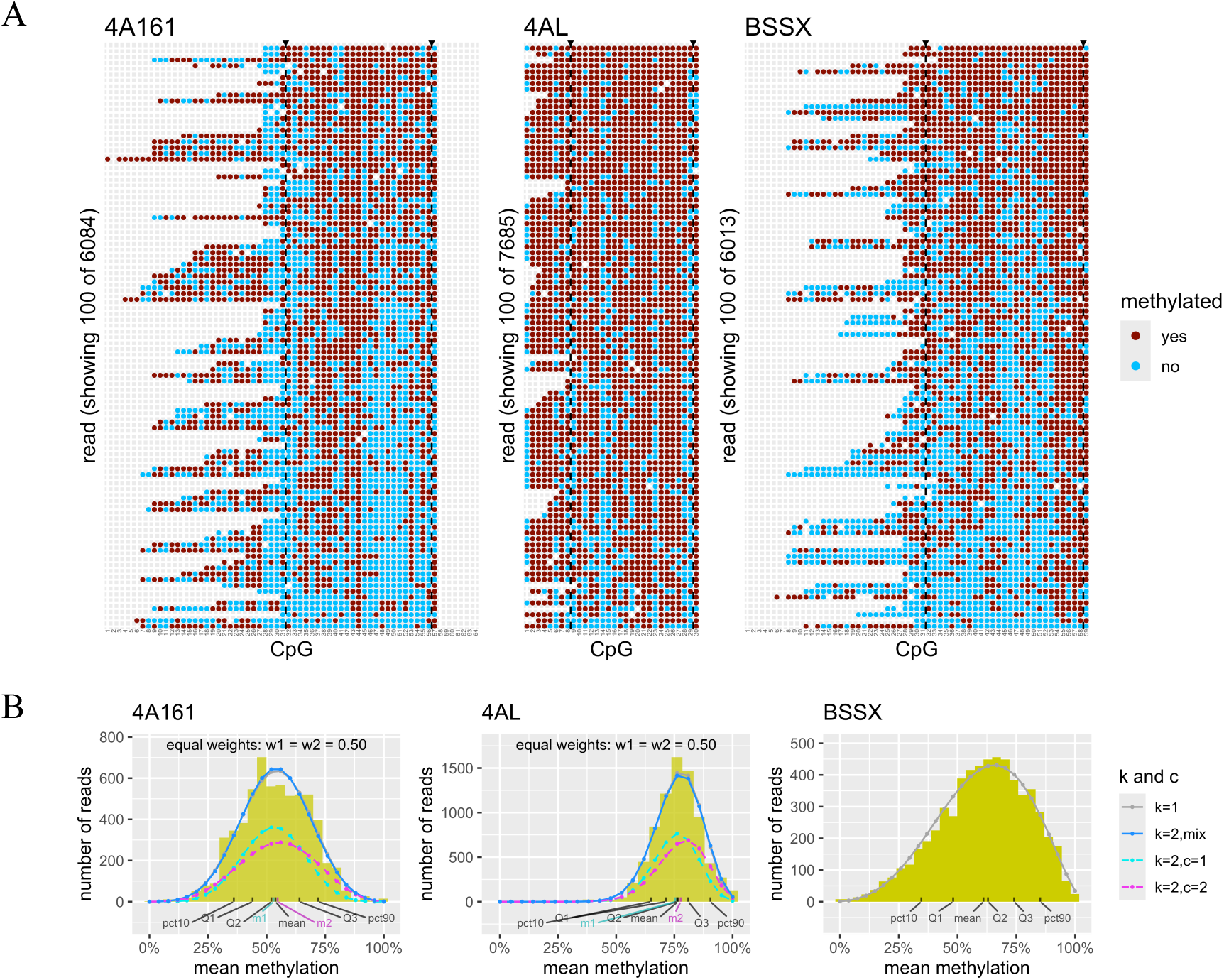
Methylation plots for an NGS sample (non-FSHD) with 4A161 and 4AL alleles. A: Grid plots for 100 randomly selected reads for BSSA (left), BSSL (center), and BSSX assays for PJL-10186 (replicate 3). Each column represents a CpG and each row represents a single read, with reads sorted by decreasing mean methylation. CpGs are colored based on methylations calls from bisulfite sequencing: red for methylated and blue for unmethylated. All CpGs in the reference amplicons are shown, and dashed vertical lines delimit the CpGs used for downstream analyses. B: Histograms of per-read mean percent CpG methylation. Curves show densities for beta distributions fitted to counts of methylated CpGs with a single beta-binomial (k = 1, gray curve) and by a mixture of two beta-binomial distributions (k = 2, blue curve), scaled to have the same area as the histograms. (Although the beta densities often closely follow the histograms, the count data has additional variance from binomial sampling.) Density curves are also shown for the two mixture components (c = 1, dashed cyan; c = 2, dashed magenta), each scaled by the weight of the components in the mixture, which is here fixed at w1 = w2 = 0.5. The means m1 and m2 for the two mixture components are indicated at the bottom border of the histogram, along with summary statistics pct10, Q1, Q2, Q3, pct90, and mean. The closeness of m1 to m2 is consistent with this subject (PLJ-10186; non-FSHD) having a single 4A161 and a single 4AL allele.

**Figure 5:**
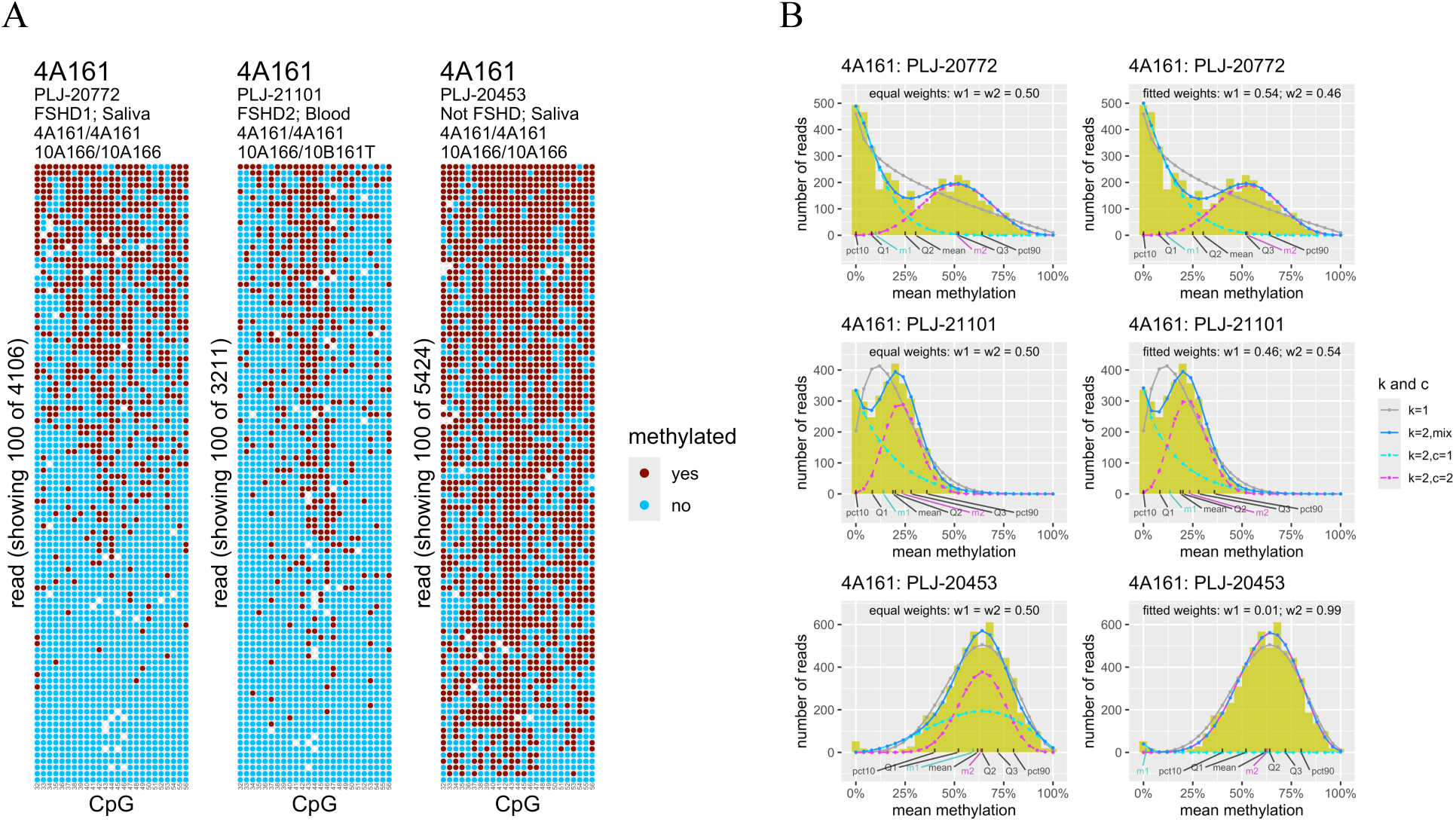
Methylation plots for three NGS samples with two 4A161 alleles. A: Grid plots for 100 randomly selected reads for BSSA assays for subjects that are genetically FSHD1 *(left)*, FSHD2 (*middle*), and non-FSHD *(right)*. Chr4 and chr10 D4Z4 array types are indicated above the plots. B: Histograms of per-read mean percent CpG methylation for the same three samples (ordered from *top to bottom*), with curves for beta probability densities from mixtures of two beta-binomial distributions overlayed (see legend for Figure 3B). In the left column the weights (w1 and w2) of the two components are each clamped at 0.50, and in the right column they are estimated from the data along with the beta-binomial parameters. Note that for sample PLJ-20453 *(bottom)* there is a small percentage of reads (∼1%) with zero percent methylation. When weights are forced to be equal *(left)*, the two components of the mixture model have similar means (m1 and m2) but the variance for one is higher to accommodate those reads. When weights are allowed to be unequal *(right)*, a component with weight w1 = 0.01 captures the reads with near-zero methylation and a component with weight w2 = 0.99 captures the remaining reads.

D4Z4caster uses the Snakemake workflow management system for scalable and reproducible analyses (52). The workflow has many components and dependencies, including Bismark (47), Bowtie2 (53), FastQC (54), Samtools (55), R and a dozen or so R libraries (including bblme for MLE), plus custom R and Python code. To simplify installation and usage all the necessary components have been “containerized” into a portable cross-platform Docker image (56), available in a DockerHub repository (https://hub.docker.com/r/kingod/d4z4caster). Source code is available in a GitHub repository (https://github.com/oliverking/d4z4caster_base) under a GNU General Public License. That repository also includes usage instructions and additional details on the computational methodology.

### Quality Control

Quality control (QC) steps were performed at multiple stages of the analysis: as pre-processing of the input files for D4Z4caster, within D4Z4caster, and as post-processing of the D4Zcaster output files. Pre-processing: Low-quality NGS reads were removed and low-quality bases were trimmed (trim-qual-cutoff = 15) with the Torrent Suite Software (Thermo Fisher Scientific) to produce the uBAM files used as input to D4Z4caster. Within D4Z4caster: FastQC was run on unaligned reads to generate a QC summary report. After the alignment and methylation-call extraction steps, additional processing was done with custom R code. Reads were removed if they had methylation calls for fewer than 90% of the CpG in the prespecified range of high-coverage CpG to use for scoring, or if over 10% of the cytosines in a non-CpG context (i.e., in CH context) were not converted to thymine. If multiple reads agreed exactly on their methylation calls at all CpG sites and their C>T conversion at all CH sites, only one of the reads was retained to reduce the potential impact of PCR duplicates. It is possible that this leads to elimination of some reads that were not PCR duplicates that happen to agree, and it is possible that some PCR duplicates remain that are not filtered out due to sequencing errors or due to different amounts of trimming leading to distinct patterns of C>T calls at CpG or CH sites. Estimates of allele-specific methylation were performed on the remaining reads and output by D4Z4caster. Post-processing: The post-processing QC steps applied to the output files for each sample are described below.

#### Filtering based on read count

Reference sequences with relatively few reads for the sample were dropped, as these may reflect low levels of sequencing errors, barcode misassignment in demultiplexing of samples in the same run, or carry-over of signal from samples from previous runs (57, 58). A minimum of 5 reads to a reference sequence was required for Sanger assays and 200 for NGS assays. Moreover, it was required that the number of reads was at least 1% that of the reference sequence with the most reads, and that the number of reads for BSSA-type reference sequence (4A161, 4A166, 10A167T, 4A161x55) was at least 10% that of the BSSA-type reference sequence with the most read.

#### Flagging potential clonal artifacts

Reference sequences were flagged as potentially having severe clonal artifacts if certain quartiles (Q1, Q2, Q3) or percentiles (pct10, pct90) of the mean per-sequence methylation were too close together (Figure 6). The criteria were as follows: pct90 – pct10 < 1; Q3 – Q1 < 1 with Q2 > 5 and at least 6 reads; (Q3 – Q2 < 1) or (Q2 – Q1 <1) with Q2 > 5 and at least 12 reads. These are simple heuristic criteria, with adjustments to reduce false alarms for samples with fewer than 12 reads or with Q2 under 5%; more complex strategies that explicitly consider the positions of methylated CpGs are also possible.

**Figure 6:**
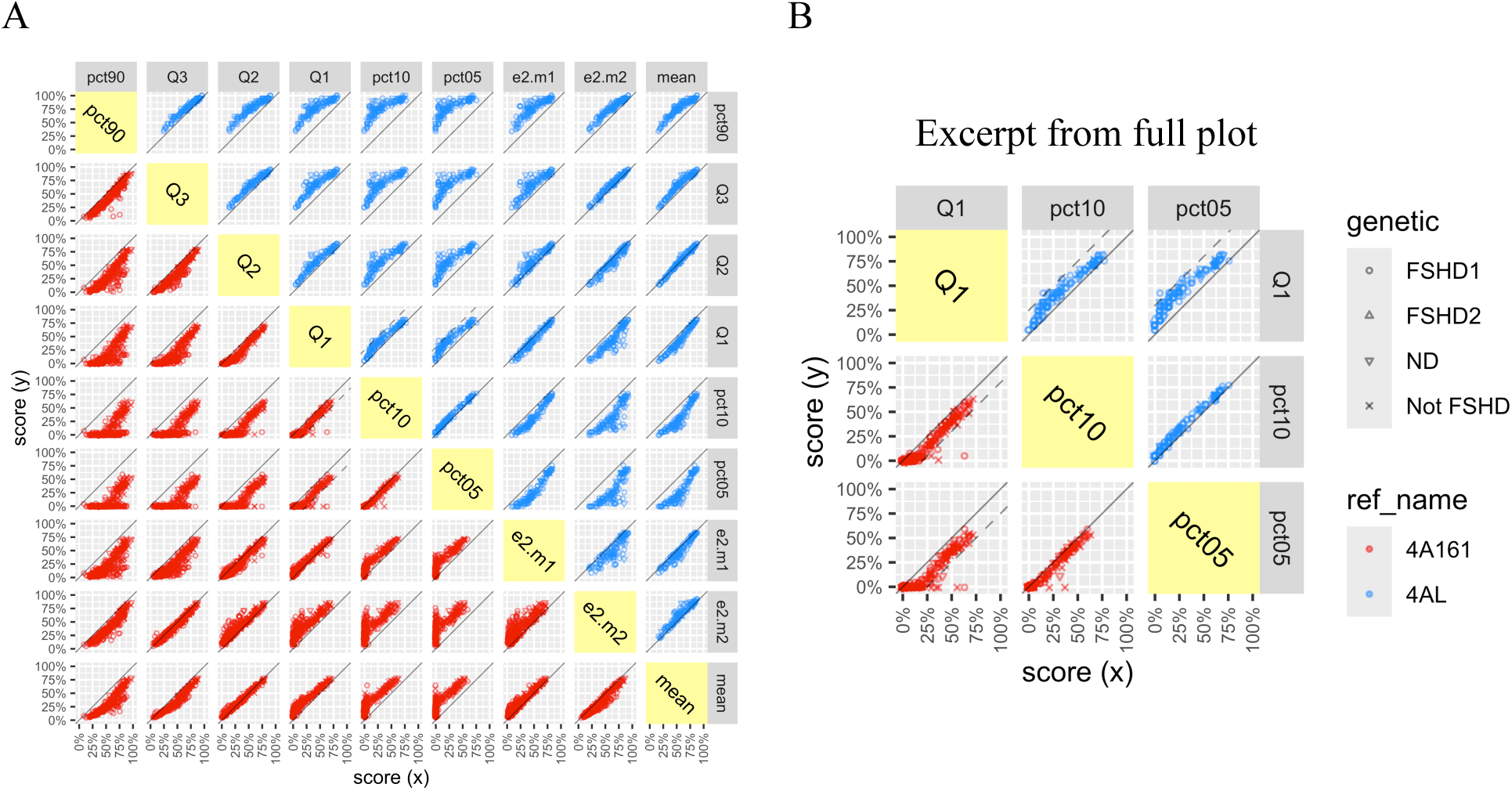
Comparison of methylation summary scores. Each dot represents an NGS run for a sample in the UNR IRB study, including some samples that were run in triplicate for the NGSx3 validation sub-study. Symbols represent FSHD genetic status (see legend, with ND= not determined). In panel A, methylation summary scores shown for each sample are the 5^th^, 10^th^, 25^th^ (Q1 [first quartile]), 50^th^ (Q2 [second quartile; median]), 75^th^ (Q3 [third quartile]), and 90^th^ percentiles, the estimated means e2.m1 and e2.m2 for a two-component beta-binomial mixture model, and overall mean. Sub-plots above the main diagonal in yellow are for 4AL methylation and sub-plots below the main diagonal are for 4A161 methylation. Scores for percentiles are jittered slightly (by up to +/- 1) to reduce overlap of points. Samples for which 4A161 assays have Q1 – pct10 > 20 or Q1 – pct05 > 25 are flagged as having potentially mosaic 4A161 alleles, and 4AL assays with Q1 – pct10 > 25 or Q1 – pct05 > 30 are flagged as having potentially mosaic 4AL alleles. These cutoffs are indicated by dashed lines, which are clearer in the excerpted plot in panel B. Note that these dashed lines are above the solid diagonal lines for 4AL and below the solid diagonal lines for 4A161 since the scores on the x and y axes are reversed.

#### Flagging potential mosaic alleles

The distribution of average percent methylation across reads for a reference sequence can be used to flag potential mosaicism of an allele. For the results presented in the study we flagged alleles that had much a higher score for Q1 than for pct10 and/or pct05. 4A161 alleles were flagged if Q1 – pct10 > 20% or Q1 – pct05 > 25% and 4AL alleles if Q1 – pct10 > 25% or Q1 – pct05 > 30% (Figure 6). A minimum of 40 reads was also required, so few alleles from Sanger samples were flagged. While one could potentially also consider pct03 or even lower percentiles to capture lower levels of mosaicism, low-levels of barcode misassignment or sample carry-over may result in false-positives (see for example PLJ-20453 in Figure 5). (Whether mosaicism for a contracted allele results in clinical FSHD may depend on both the allele frequency and length, but even a germline mosaic allele present in only 5% of cells may – if inherited – be present in all cells in offspring, so has potential relevance for genetic counseling.) Alleles can also be flagged based on getting a much better fit for a two-component mixture model with unequal weights for the components (u2 model) compared to one with the weights both set to 0.5 (e2 model), or a three-component mixture model (with [e3] or without [u3] equal weights) compared to a two-component model with equal weights. While the BIC in principle provides a way to compare models, with many thousands of reads even small violations of model assumptions can lead to more complex models having much better fit even after penalizing for addition parameters (i.e., much lower BIC). Therefore, to avoid false-positives for flagging mosaics, in addition to comparing BICs one should consider whether the difference in estimate means between alleles is large enough to be meaningful (and relevant to FSHD) and whether the weight w1 is small enough to potentially represent contamination rather than mosaicism. Such an approach will be refined in future work. The weight w1 in the u2 mixture model directly provides an estimate of the percent mosaicism for samples predicted to have a single allele of type 4A (or 4AL) based on PCR. For samples with two alleles of type 4A (or 4AL) the estimated percent mosaicism would be 2 × w1, as ∼50% of the reads would be expected to be from the allele that is not mosaic

## Statistical Analysis

The statistical methods used for estimating allele-specific methylation levels within a sample are described in the D4Zcaster section above. Below is a description of the statistical methods used for comparisons of FSHD and non-FSHD groups and for predictions of FSHD status based on methylation levels.

### Receiver Operating Characteristic (ROC) curves

ROC curves were computed based on single scores *s* taken directly from D4Z4caster output (e.g., Q1 for 4A161 for NGS assays) and for derived scores from logistic regression models (and combinations of logistic regression models). For any fixed cutoff *C*, samples are classified as FSHD if they have score *s* < *C* and non-FSHD otherwise (where the direction of the inequality may be reversed depending on the semantics of the score). This partitions the *N* samples into four sets: true positive (TP), false positive (FP), true negative (TN), and false-negative (FN), where positive means genetically FSHD and negative means genetically non-FSHD. ROC curves show the True Positive Rate [TPR = #TP/(#TP + #FN) = sensitivity] vs the False Positive Rate [FPR = #FP/(#FP + #TN) = 1 – specificity] for every cutoff *C* in the entire range of values observed for the score *s*. The area under the ROC curve (AUC) can be interpreted as the probability that a randomly-selected FSHD sample has a lower score *s* than a randomly-selected non-FSHD sample. The overall classification accuracy is given by (#TP + #TN)/*N*, though this can depend strongly on the relative proportions of FSHD and non-FSHD samples in the dataset so may be misleading. To avoid this dependence on relative class proportions we instead report G-mean, the geometric mean of the sensitivity and specificity (i.e., the square-root of their product), as an overall assessment of classification performance at any given cutoff (59). To assign confidences to class predictions and to estimate the probability of a sample belonging to class FSHD, the relative class proportions matter, serving the role of prior probabilities for Bayesian inference. For some of our datasets 80% or more of the samples are from individuals with genetic (and clinical) FSHD, whereas if one were applying this assay to randomly-selected individuals from the general population fewer than 1% of the samples would be expected to be from individuals that meet the genetic definition of FSHD. AUCs and their 95% confidence intervals (CIs; by DeLong’s method) were computed using the R package pROC (1.19.0.01) (60), separately for NGS and Sanger data and for both assay types combined. If there are differences in methylation scores depending on assay type, it can be beneficial to account for this when assigning the scores used for ROC analysis, as this may affect the rank-order of samples and thus may affect the AUC. To do this we used logistic regression models that included an additive factor for assay type as detailed below. When there is just one methylation feature in addition to assay type in the regression model, the regression is monotonic for samples of the same assay type, so the ROC, AUC, and CIs for single assay types are insensitive to the regression parameters. The CIs for the AUCs of the combined ROC curves are computed based on the fitted regression parameters and do not account for the uncertainty in these estimates.

### Logistic regression

To estimate the probability of a sample belonging to the genetic class FSHD, we fit logistic regression models to the data coded with binary response 1 for FSHD and 0 for non-FSHD, using one or more methylation-based score (e.g., Q1, e2.m1 or pct10) as predictor variables and assay type (NGS or Sanger) as an additive categorical covariate. One could also potentially include sample type (saliva, blood, muscle biopsy, etc.) as a categorical covariate, but as the UNR IRB dataset used for model training consisted almost entirely of saliva samples, we did not do so here.

### Offsets

Logistic regression models have the convenient property that the output (which is between 0 and 1) can be interpreted as an estimate of the posterior probability of a sample belonging to class 1, and the prior probability can be adjusted by changing the intercept term in the fitted model, or can be adjusted during the model fitting by including an offset term (61). We adjusted for class imbalance during model fitting by using log(#FSHD/#non-FSHD) as an offset during model fitting so that predictions from the model correspond to a scenario in which both classes have equal prior probability. (Here and elsewhere log denotes natural logarithm, with base e). One can change this offset when making predictions if one wishes to use a different prior probability, based on population demographics, context of use of the assay, pedigree information, etc., though calibration of the predicted probabilities may degrade for extreme adjustments to the prior when (as it the case here) the models only approximate reality (62). In models that included a factor for assay type (Sanger or NGS) the offset was computed separately for the two assay types, since relative proportions of FSHD to non-FSHD samples sometimes differed between the Sanger and NGS assay types, and differences in coefficient for assay types would otherwise in part reflect those differences in proportions.

### Data transformations

For scores that are on the scale of percent methylation (e.g., Q1) and are thus constrained to be between 0% and 100% we scaled the data by 100 to be in the range from 0 to 1 and performed a logit transformation *z* = log(*x*/(1 – *x*)) prior to fitting the models to avoid the compression of values for scores near zero. The motivation for this is to improve the linearity of the relationship between the predictors and the log-odds *w* = log(Pr(FSHD)/(1 – Pr(FSHD))) of a sample belonging to class FSHD, as this underlies the interpretation of f(*w*) = 1/(1 + exp(–*w*)) as a posterior probability, where the logistic function f is the inverse of the logit function. We also included a small offset of 0.01 to avoid taking the logit of 0 or 1, so formally used the transformation *z* = log(*x’*/(1 – *x’*)) where *x’* = (*x* + 0.01)/1.02. (Or equivalently x’= (*S* + 1)/102 where *S* is the score on the scale of percent methylation). In some plots, scores *x* that had been logit-transformed prior to fitting the logistic regression models have been converted back to the original linear scale to aid interpretation. The resulting curves, while still generally sigmoidal, can appear distorted compared to standard logistic curves.

### Regularization

For some datasets certain scores perfectly separated the FSHD from control samples (e.g., *s* = Q1 for 4AL in UNR IRB dataset), which can lead to instability in logistic regression fits, with coefficients approaching infinity. To combat this, we used regularized logistic regression, which strikes a balance between the goodness of fit to the data and the magnitude of the coefficients. Specifically, we used the R package glmnet (4.1-10) with elastic net mixing parameter alpha = 0.5 for an equal penalty on the *L*_1_ norm (LASSO regression) and the *L*_2_ norm (ridge regression) of the vector of coefficients (excluding the intercept and offset) (63). Categorial variables were dummy-coded as 0/1 indicator variables with tidymodels (1.4.1) (64), and variables were standardized prior to fitting but with coefficients returned on the original scale. The function cv.glmnet was used to automatically select the penalty parameter lambda for the final models based on the largest value of lambda that gave deviance within 1 standard error of the minimum (lambda.1se) by 10-fold cross validation. The dummy-coded assay type coefficient representing the difference in intercepts for NGS vs Sanger data is also penalized to stabilize estimates. As the *L*_1_ penalty favors sparseness, in some cases the fitted value of this coefficient was exactly zero, yielding a model that makes identical predictions for NGS and Sanger data (using an offset term to adjust for unequal balance in FSHD vs non-FSHD samples for NGS vs Sanger data). While we could in principle use an *L*_1_ penalty to select among the dozens of scores output by D4Z4caster (only some of which are shown in Figure 6), many of those scores are strongly correlated so the decision on which features get included will be unstable. The *L*_2_ penalty favors inclusion of all correlated variables rather than just one, but this would lead to overly complex models that offer little predictive advantage over simpler models. Thus, we fit models using one or two features selected based on our experience using these assays over many years, rather than fitting models on all features.

### Combining scores for 4 and 4AL alleles

The logistic regression models above give estimates of P_A_ = Pr(genetic FSHD from 4A allele | 4A methylation) and P_L_ = Pr(genetic FSHD from 4AL allele | 4AL methylation). As FSHD is a dominant gain of function disease, for subjects of type A/L, who have both 4A and 4AL alleles, one can take the larger of these two values, max(P_A,_ P_L_), as an estimate of Pr(genetic FSHD from either allele | 4A methylation and 4AL methylation). We also computed a refinement of this, P_A/L_ = 1 – (1– P_A_)(1– P_L_), which is nearly the same as max(P_A,_ P_L_) when one is close to zero but can be larger otherwise: for example, if P_A_ = 0.5 and P_L_ = 0.5 then their maximum is 0.5 but P_A/L_ = 1 – (1– P_A_) (1– P_L_) = 0.75: the chance of getting at least one heads when flipping two independent coins. The use of an offset to compensate for class imbalance can be viewed as implicitly assigning priors on the probabilities of A-type and L-type alleles being associated with FSHD. This formulation makes assumptions of independence between alleles that need not strictly hold, due for example to the influence of *SMCHD1* variants that affect methylation levels of both alleles, and also P_A_ and P_L_ are defined without regard to the number of 4A and 4AL alleles, so there are not simple per-allele priors. While further refinements of this approach could be used to make inferences about potential FSHD2 cases, for this we instead consider the methylation levels from the BSSX assay, as this is applicable to allele types A and L as well as A/L.

### Calibration of predicted probabilities

We used the R package CalibrationCurves (3.1.0) (65) to assess the calibration of predicted probabilities from the combined 4A/4AL scoring by fitting smooth local regression curves (LOESS) relating predicted probabilities of subjects being genetically FSHD to observed proportions of subjects that were genetically FSHD. Because the logistic regression models for 4A and 4AL alleles each included an offset term to compensate for the strong imbalance in classes sizes (with many more FSHD than non-FSHD subjects), it is expected that predicted probabilities of subjects being genetically FSHD are lower than observed proportions for these subjects. To compensate for this, calibration curves were also computed after the smaller class (non-FSHD) was randomly up-sampled to match the size of the larger class, simulating a 50% prior probability of being genetically FSHD (which was also the intent of the offset terms used in the logistic regression models). The regularization parameters lambda used in our elastic net models were the values that minimized the mean cross-validation deviance (lambda.min) and the largest values with deviance within one standard deviation of this minimum (lambda.1se). The latter provides an extra hedge against over-fitting, but it can result in conservative probability estimates in the sense that extreme predicted probabilities both above 0.5 (predicted FSHD) and below 0.5 (predicted non-FSHD) can be pulled toward 0.5.

## Results

### Sample collection, haplotyping, and targeted BSS (Sanger and NGS)

In this study we aimed to transition our targeted BS-PCR approach for FSHD analysis (Figure 1) from Sanger-based sequencing to a more efficient, accurate, and clinically relevant NGS approach. We used several large genetically confirmed cohorts (Table S1) to validate both the Sanger and NGS approaches as highly accurate for identifying DNA methylation signatures consistent with the presence of FSHD1-sized deletions or FSHD2 mutations in an FSHD-permissive genetic context.

Consistent with our published workflow for epigenetic determination of FSHD (41), the first step was again identifying the proximal SSLP and distal haplotypes for each sample. While this step could potentially be used to eliminate those samples that lack any FSHD-permissive alleles, we instead used this for confirmation of results in later steps. All gDNA samples, regardless of haplotype, were then bisulfite-converted and PCR-amplified for the BSSA, BSSL, and BSSX regions (Figure 1). Those samples undergoing Sanger sequencing to determine methylation profiles followed our published protocol (41), while those samples to be used for NGS were subjected to an alternative second nested PCR using barcodes fused to NGS-compatible versions of our previously used oligonucleotide primers (Table S2). Reaction products were visualized by agarose gel electrophoresis and amplified products were gel-purified, and then either TA cloned for direct-from-colony Sanger sequencing or used for generation of a barcoded and multiplexed NGS library (Figure 2). Sequencing results were then analyzed using the D4Z4caster program (Figures 3).

We began performing the original Sanger-based sequencing protocol in 2013 and did not transition to NGS format until 2020; therefore, we were interested to know how changing this format would affect our prior results. In addition, since each NGS run typically processes 70 samples at a time (and up to 96) to maximize cost effectiveness, there are still occasions when single samples need to be analyzed quickly, so Sanger sequencing is still utilized. Thus, we compared Sanger sequencing results with NGS results on the same samples. We assembled a set of 77 samples from a mix of genetically-confirmed FSHD1 (n = 33) and FSHD2 (n = 5) cases; genetically confirmed healthy controls without FSHD (n = 13); and individuals with undetermined FSHD genetic status (n = 26). One complication in a direct comparison of Sanger to NGS results is that the Torrent Suite Software trims low-confidence bases from NGS reads, and this typically resulted in a sharp drop-off in coverage towards the 5’ ends of the amplicons (Figure 4A). As the location at which the drop-off begins was generally consistent across samples (perhaps related to local GC content), for each reference amplicon we defined a range of CpG with high coverage to be used for NGS analyses: CpGs 32-56 for the BSSA assay, CpGs 9-29 for the BSSL assay and CpGs 32-58 for the BSSX assay. Since different regions in the amplicons can have different mean methylation – just as different regions of D4Z4 repeats can have markedly different mean methylation (39) – the most direct comparison of the Sanger and NGS approaches would use the same CpG sites for both. Using the same sites may also be useful if one wishes to use cutoffs for disease diagnostics that are the same for both approaches.

We first examined the impact of scoring Sanger samples using just these restricted ranges of CpGs (which we refer to as the 3’ CpGs), compared to using all CpGs in the amplicons, based on all 325 samples with Sanger data in the study (Figure 7A). We next compared the results from Sanger assays using the 3’ CpGs to NGS assays using the 3’ CpGs, based on 76 samples from which both Sanger and NGS assays were performed (Figure 7B). In both cases we computed correlations of mean per-sample methylation scores, separately for the BSSA, BSSL, and BSSX assays. For BSSA and BSSL assays we also computed correlations for the Q1, e2.m1, and pct10 scores that are useful in predicting disease status. Per-sample methylation scores from D4Z4caster for these samples and all other samples discussed herein are in Table S4. Pearson correlation coefficients were >0.95 for all comparisons aside from BSSX mean in NGS vs Sanger, which was 0.90. Slopes, intercepts, and residual standard deviations for linear regression fits are also shown in Figure 7A-B. Slopes were typically close to 1.0 for the NGS vs Sanger comparisons and the intercepts were typically close to zero, although for the BSSA Q1, e2.m1, and pct10 scores the intercepts ranged from 3.0 to 5.0 on the scale of percent methylation. This suggests that it may be useful to consider using different cutoffs when predicting FSHD status based on NGS vs Sanger assays, as we do in this study, although the effect of doing so may be mild.

**Figure 7:**
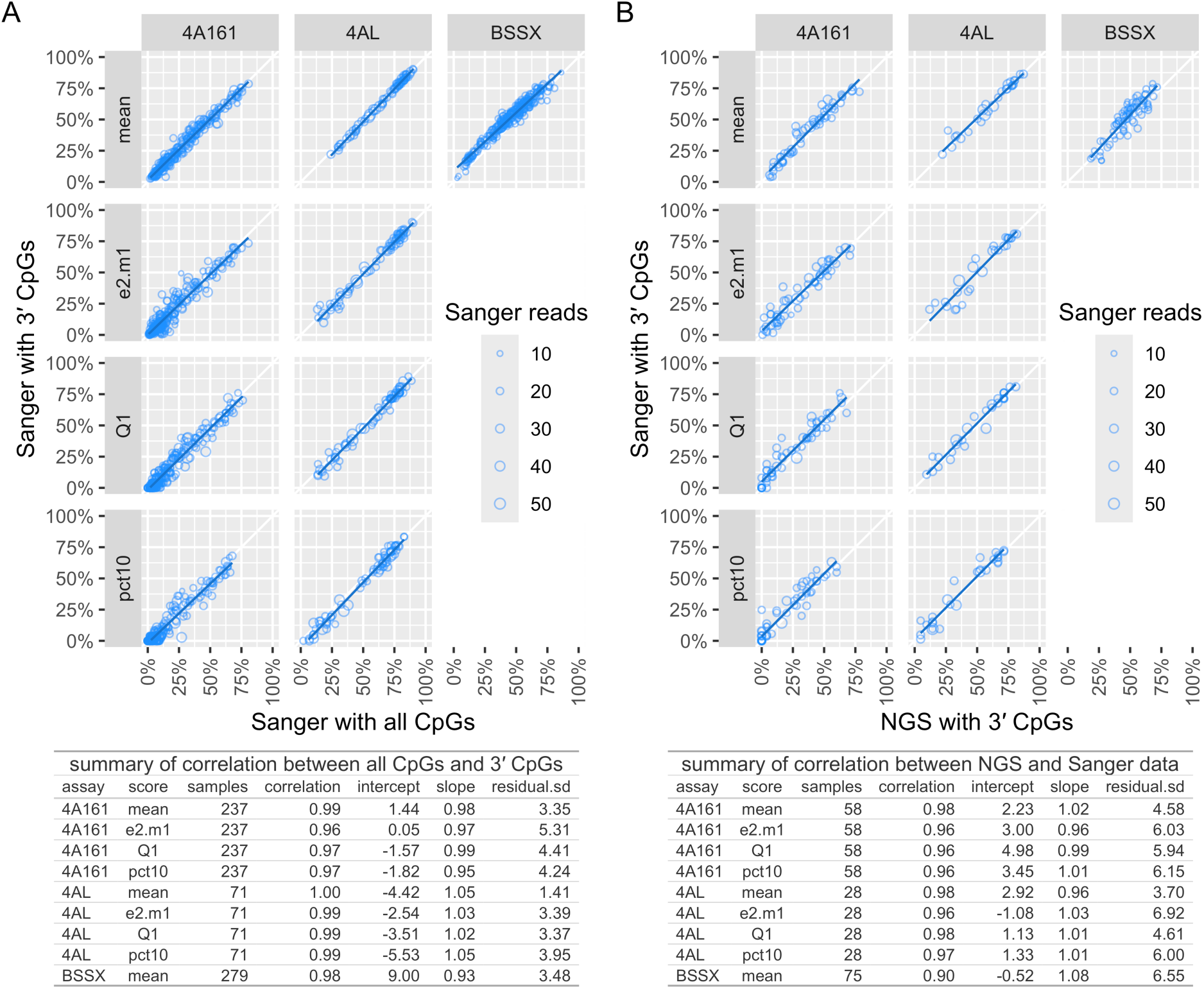
Comparison of scores from Sanger and NGS assays. Scores for the NGS assays are based on a subset of the CpGs (referred to as 3’ CpGs) from the PCR amplicons that had high coverage after trimming low-confidence bases from NGS reads. A: Comparison of scores for Sanger assays using only these 3’ CpGs vs all CpGs. Each point in the plots represents a single sample, with sample count from 71 to 279 depending on assay type (4A161, 4AL, or BSSX). Point size denotes number of Sanger reads. B: Comparison of scores from Sanger vs NGS assays, both using only the 3’ CpGs. Sample counts range from 28 to 75 depending on assay type (4A161, 4AL, or BSSX). Correlations were high for both panel A and B, in the range of 0.96–1.00 for all assay types and scores aside from 0.90 for BSSX mean in B. Slopes and intercepts of lines of best fit (from ordinary least squares regression) are as indicated, and the residual standard deviation (sd) with respect to this line was in the range of 1.4 –5.3 for A and 3.7 –6.9 for B (on the scale of % methylation).

To assess the reproducibility of this new NGS protocol, a set of 27 samples was selected for analysis in triplicate. Samples were from 12 FSHD1, 3 FSHD2, and 10 healthy control (not FSHD) individuals, all genetically confirmed, and 2 with undetermined genetic status. Each of the three replicates started with a new bisulfite conversion of the gDNA sample and the replicates were sequenced on independent NGS runs. The D4Z4caster results showed excellent reproducibility for our NGS approach, with the coefficient of determination r_2_ ranging from 0.94 to 1.0 and residual standard deviation ranging from 1.4 to 4.7 depending on the score (mean, Q1, e2.m1, or pct10) and assay (BSSA, BSSL, or BSSX) (Figure 8; Table S4). One sample showed appreciably higher between-replicate variance than the others: PLJ-10382, due to replicate 1 disagreeing with replicates 2 and 3 in the BSSA assay, most notably for the Q1 and pct10 scores. As discussed further later, this sample appears to be mosaic for a 4A161 contraction (Figure 9), which can result in high variance of methylation percentiles near the frequency of the mosaic allele.

**Figure 8:**
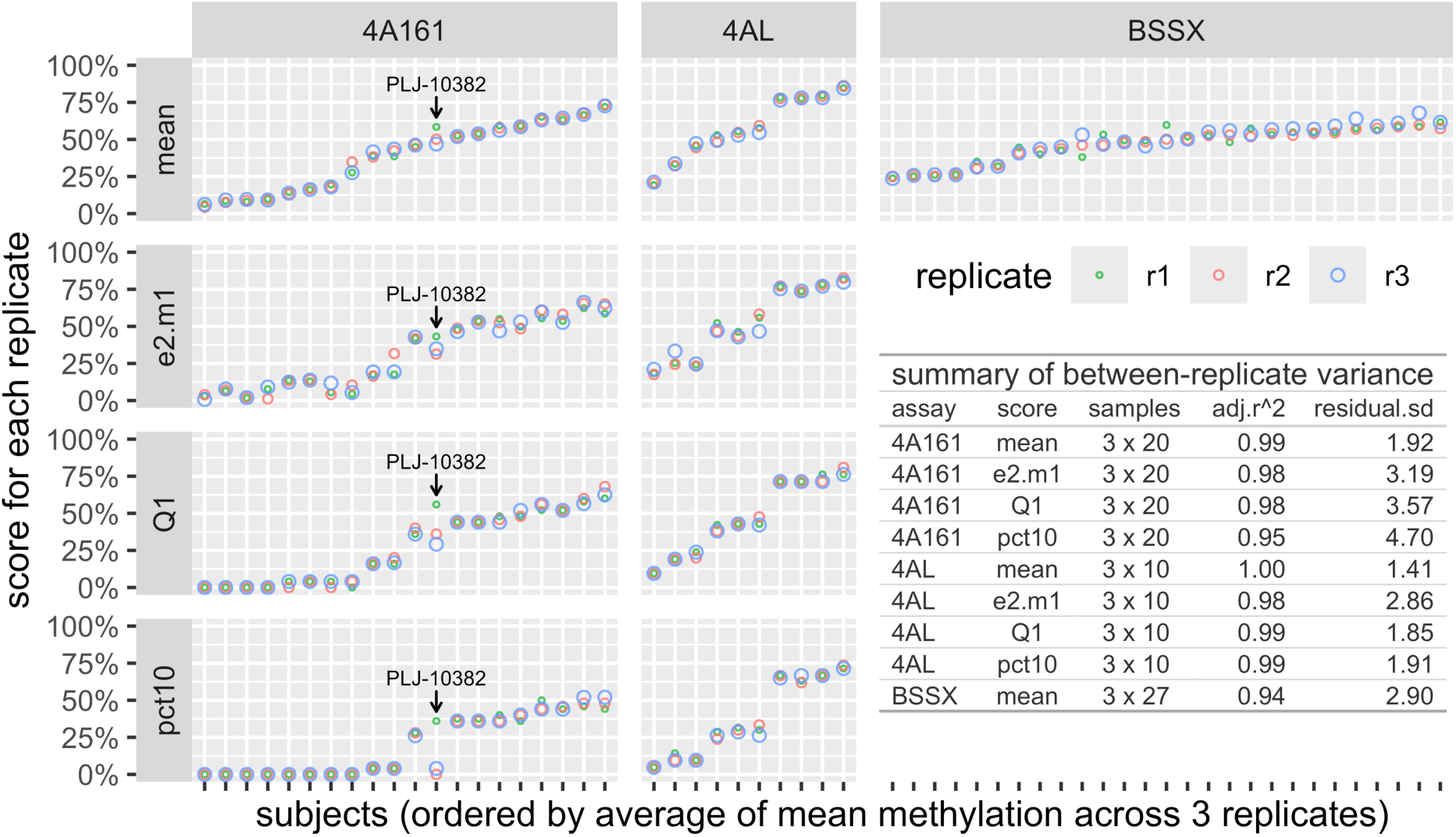
Technical reproducibility of scores from NGS assays. Comparison of methylation results from three NGS replicates for each of 27 samples. Each column represents a sample and each point in that column represents a replicate. The number of samples is 20 for the 4A161 assay, 10 for the 4AL assay, and 27 for the BSSX assay (with overlapping samples for those three assays). A linear model with a factor for subject was fit to the data for each assay and score type, and the coefficient of determination r^2^ (adjusted for the number of predictors) was high (0.94 –1.00) for all assay types (4A161, 4AL, or BSSX) and scores (mean, e2.m1, Q1, or pct10). The residual standard deviation with respect to the fitted model was in the range of 1.4 to 4 .7 (on the scale of % methylation). The most overt example of scores that disagree between replicates is 4A161 methylation in sample PLJ-10382 (*labeled*), particularly for Q1 and pct10. As discussed in the main text, this sample appears to be mosaic for a 4A161 contraction, which can result in high variance of methylation percentiles near the frequency of the mosaic allele.

**Figure 9:**
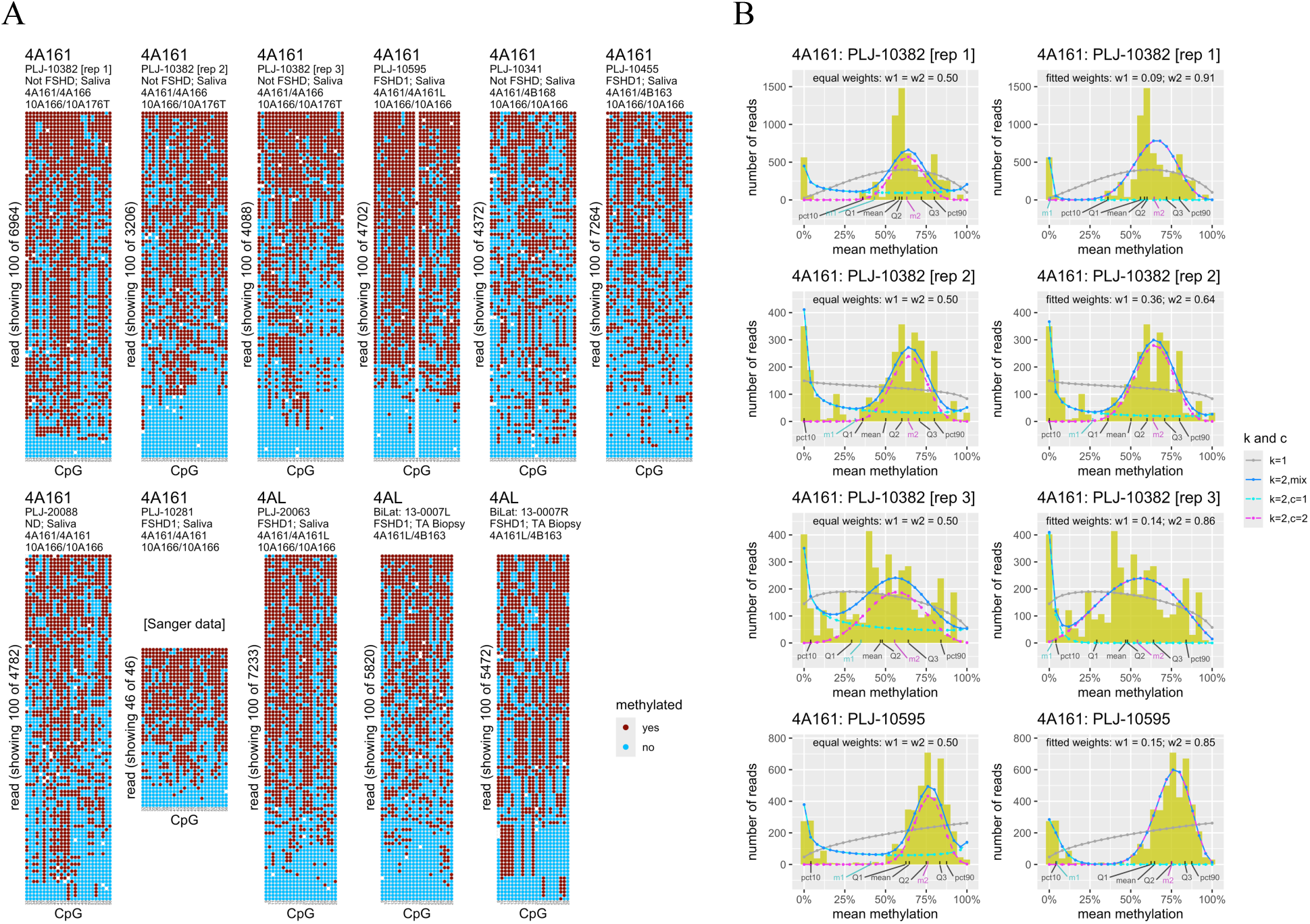
Methylation plots for alleles flagged as potentially mosaic. Flagging was based on an unusually large difference in the 25^th^ vs 10^th^ percentile (Q1 vs pct05) and/or the 25^th^ vs 5^th^ percentile (Q1 vs pct10) (see Figure 6 and Methods). A: Grid plots of 100 randomly selected reads. All plots are from NGS assays except for the plot with only 46 reads (sample PJL-10281), which was from Sanger sequencing. Plots are for 4A161 alleles aside from the last three plots, which are for 4AL alleles. The first three plots are replicates for PLJ-10382, an outlier for Q1 and pct10 in the NGSx3 plots in Figure 8. The unusual missing 44th CpG for PJL-10595 suggests mosaicism of a single 4A161 allele rather than a mixture of 4A161 reads from different alleles (and the sample has a 4AL allele). Of the last two plots, only the first, 13-0007L, was flagged as potentially mosaic, and 13-0007R is included for context as these are from left and right TA of the same donor. This does *not* mean that mosaicism is present only in the left TA, as the flagging can have false positives and false negatives. The right TA just slightly missed the cutoff for flagging 4AL alleles based on Q1 – pct05 > 30. B: Fits of beta-binomial mixture models for the first four samples shown in A, one per row. The format of these plots is described in the legends to Figures 4B and 5B. Plots on the left show mixtures in which both components are forced to have equal weight (w1 = w2 = 0.5) and plots on the right show mixtures in which these weights are estimated along with the beta-binomial parameters.

We next examined how well several methylation summary scores could distinguish between samples from genetic FSHD and non-FSHD individuals. The scores we considered for the BSSA and BSSL assays were the following:

1. **mean**: the mean (over all bisulfite reads) of the mean (over all high-coverage 3’ CpGs) percent methylation. This has the conceptual weakness that high methylation from a non-contracted allele can mask low methylation from a contracted allele. It would, for example, give the same score (50%) to an FSHD subject with a contracted 4A allele with 20% mean methylation and non-contracted 4A allele with 80% mean methylation as it would for a non-FSHD subject with two non-contracted 4A alleles each with 50% methylation (or one 4A allele with 50% methylation and one 4B allele not picked up by the assay). It is nonetheless interesting to consider this score, as it shares a limitation of methylation assays based on single CpG sites, in which one cannot use information from neighboring CpGs on the same read or DNA molecule to estimate allele-specific methylation. Using different cutoffs for 4A/4A and 4A/4B samples has been proposed as a strategy for improving sample classification accuracy based on mean methylation at other CpGs (66), but this does not address the example scenario described above.
2. **Q2:** the median of the distribution of mean percent CpG methylation for all reads. This estimate can be unstable when sampling from a strongly bimodal distribution in which each mode has ∼50% frequency, as may occur for samples with two 4A or two 4AL alleles. In the example scenario above, depending on how tightly-clustered the scores for each allele are about their respective means (or medians, which can differ from means due to asymmetry in distributions), the overall sample median could be much under 50% (as low at ∼20%) or much over 50% (as high at ∼80%), and which of those is observed could be essentially random, determined by which allele happens to be sampled more frequently during sequencing. (It is unlikely for both alleles to be sampled *exactly* the same number of times: for *N* = 2*n* reads the probability scales roughly as 1/sqrt(*N*).) We include results for Q2 mainly because it has been used as a summary statistic in other studies (67, 68).
3. **Q1**: The first quartile (25^th^ percentile) of the distribution of mean percent CpG methylation for all reads, as in our initial work (41). In the example scenario above, Q1 could be in the vicinity of 20%, since if the methylation distributions for the two alleles have little overlap then Q1 for both alleles combined is approximately the median (Q2) for the shorter allele. A conceptual weakness of this approach is that if the distributions for the two alleles have a lot of overlap (or if there is just one allele of this type), then Q1 for the combined alleles will be close to Q1 rather than Q2 of the first allele. Thus, as an allele-specific estimate for the less-methylated allele, the score can be biased by the methylation of the more-methylated allele. It is also biased by the number of alleles of the type 4A (or 4AL). Although we often have information on the multiplicity of allele types from PCR assays, such information may be incomplete or inaccurate (e.g., due to translocations between chromosomes 4 and 10 or the presence of multiple *cis* D4Z4 arrays on chromosome 4 (33, 48)); this information is currently used for QC and for flagging unexpected findings but not to adjust scores or cutoffs. It is possible to use Q2 to score samples with one allele of type 4A (or 4AL) and Q1 to score samples with two alleles of the same type, provided that this information is known (68), though Q1 is still biased by the amount of overlap between the distributions of the two alleles.
4. **e2.m1**: The estimated mean methylation of the allele with lower mean methylation in a two-component beta-binomial mixture model, as in our previous work (13). This can mitigate the biases for the Q1 score by modeling the data as a mixture of two distributions whose overlap can range from minimal to extensive. When only one allele is present, even if a two-component model gives a significantly better fit to the data, the means of the components can be quite similar, mitigating the dependence of Q1 on allele number. A weakness of the e2.m1 score is that estimates are based on model assumptions – beta-binomial distributions with equal number of reads expected from each allele – that can be violated to various degrees, for both technical (e.g., clonal amplification) and biological (e.g., mosaicism) reasons. We discuss extensions to address some of the issues, by fitting mixture models with not-necessarily-equal proportions of reads from two alleles (u2) and mixture models with three alleles (e3 and u3).
5. **ex.m1**: The estimated mean methylation of the allele with lower mean methylation in a two-component beta-binomial mixture model, regularized by Bayesian model averaging with the mean methylation in a one-component beta-binomial model (see Methods). This typically has a bigger effect for Sanger data, which has many fewer reads and more uncertainty in estimated parameters.
6. **pct10:** The 10^th^ percentile of the distribution of mean percent CpG methylation for all reads. As *DUX4* expression in FSHD muscle is sporadic and expression may be more likely in nuclei that happen to have lower D4Z4 methylation, scores based on the low tail of the methylation distribution may have conceptual advantages over scores based on means or medians, whether allele-specific or not. Interpretation of pct10 as an allele-specific score depends on whether a sample has one or two alleles of the type being assayed, as it does for Q1 and other percentile-based scores. *(Note: allele-specific estimates of percentiles based on beta-binomial mixture models are included as an experimental feature in D4Z4caster and may be further explored in future work after additional refinement. One can compute percentiles directly from beta distributions or from the estimated posterior probabilities of each read originating from each allele. There is the technical issue that when mixture components have different variances, extreme points from both tails can have higher posterior probability for the component with higher variance, which may or may not reflect reality and can make estimated allele-specific tail percentiles sensitive to the fitted beta-binomial parameters in a way that the estimated allele-specific means are not.)*
7. **pct05**: The 5^th^ percentile of the distribution of mean percent CpG methylation for all reads. The same considerations apply as for pct10, but here the focus is on the more extreme lower tail.
8. **pct03**: The 3^rd^ percentile of the distribution of mean percent CpG methylation for all reads. This focuses on the even more extreme lower tail than pct05. There is the potential for diminishing returns from looking too far in the tail, both due to floor effects (i.e., loss of discriminative power if many healthy control samples have a score of zero) and due to the potential for the extreme tails to be strongly affected by even low levels of technical artifacts (e.g., rare sequencing errors resulting in reads from a ∼99% homologous 4q or 10q locus being misassigned; a small fraction of barcodes within a multiplexed NGS run being misassigned; or a small amount of barcode carry-over from a previous NGS run). For example, note that 1% of the 4A161 reads (52/5430) for the non-FSHD sample PLJ-20453 in Figure 5 had zero methylation, which is likely due to carry-over from the sample (not included in this study) with the same barcode in the previous NGS run, for which ∼70% of the 4A161 reads had zero methylation (itself potentially a clonal artifact). Keeping a record or results from prior NGS runs can be helpful for diagnosing potential carry-over artifacts.

For each allele type (4A or 4AL) and each score (mean, Q2, Q1, e2.m1, ex.m1, pct10, pct05, or pct03; Table S4) we fit a regularized logistic regression model to predict whether subjects with just one or the other of those allele types, but not both, were genetically FSHD or not based on that score (see Methods). These models included an additive factor for assay type to allow for differences in intercepts for NGS and Sanger assays, though similar intercepts were favored because the elastic net regularization penalty was applied to this factor as well as the methylation score. Models also included offset terms to account for differences in proportions of FSHD and control samples for the two assay types, and between 4A and 4AL allele types. The subjects used for fitting these models may also have nonpermissive 4B alleles or 4A166 alleles (which are typically not pathogenic even if contracted), but samples with both 4A and 4AL alleles were not used for fitting the models since we don’t know *a priori* which of them is the FSHD-associated contracted/hypomethylated allele (but see note below **[*]**). Rather, we used the models fit on subjects with just one or the other allele type to predict the FSHD status of the subjects based on just their 4A methylation [P_A_ = Prob(FSHD|4A methylation)] or just their 4AL methylation [P_L_ = Prob(FSHD|4AL methylation)]. We considered two strategies for combining these scores: P_A/L_ = max(P_A_, P_L_) and P_A/L_ = 1 – (1 – P_A_)(1 – P_L_). These scores agree closely when one of P_A_ and P_L_ is close to zero, and overall gave similar classification performance, perhaps due to the paucity of subjects with borderline predictions based on both the 4A and 4AL alleles. In what follows we mainly discuss results for the simpler max(P_A_, P_L_) score, though this choice can be revisited later as more data becomes available. By setting P_A_ and P_L_ to zero for subjects with no 4A and 4AL alleles (respectively), the score P_A/L_ can be interpreted as the predicted probability of a subject being genetically FSHD regardless of allele type: Pr(FSHD). *(**[*]** Note: one could in principle iteratively estimate the probability of the 4A and 4AL alleles in 4A/4AL samples being FSHD-associated and refit remodels for 4A and 4AL allele types including alleles from 4A/4AL samples weighted by these probabilities, but because most samples had only one or the other allele type we did not bother with this.)*

ROC curves show the tradeoffs between sensitivity and specificity that may be obtained by varying the cutoff on Pr(FSHD). The geometric mean of the sensitivity and specificity (G-mean) combines them into a single performance metric for each cutoff, and the area-under-the ROC curve (AUC) provides a classifier performance metric that is independent of any given cutoff. We used offsets in our regression models to compensate for class imbalances so that the output can be interpreted as the predicted probability of a sample being genetically FSHD in a situation in which the prior probability of that is 50%. That prior can be adjusted if desired by shifting the intercept term of the fitted models, but the important point here is that it puts predictions for different assay type (NGS or Sanger) and allele types (A, L, or A/L) on a common scale even if the corresponding methylation scores are quite different, which allows us to assess classification performance on all samples combined. For example, the methylation score corresponding to Pr(FSHD) = 0.50 is much higher for L alleles than for A alleles, which is likely due at least in part to different regions of the distal copy of *DUX4* being assayed, as there are fine-scale fluctuations in average CpG methylation in D4Z4 repeats that may be associated with local sequence features or nucleosome positioning (39).

In Figure 10, ROC curves are shown and AUCs are reported for each assay type (NGS or Sanger) and each allele type combination (A, L, or A/L) separately, and for one or both of those factors combined, all based on the e2.m1 score. Analogous figures for the Q1, ex.m1, pc10, pct05, pct03, mean, and median (Q2) scores are shown in Figures S1-S7. Figure S8 shows results for the e2.m1 score using PA/L = 1 – (1– PA)(1– PL) rather than max(PA, PL) for scoring the A/L samples, which results in curves rather than rectilinear contours of equal probability in panel C. (The overall AUCs were similar for both approaches so we focus on max(PA, PL) for simplicity, but this could be reassessed as data from more A/L samples becomes available.)

**Figure 10:**
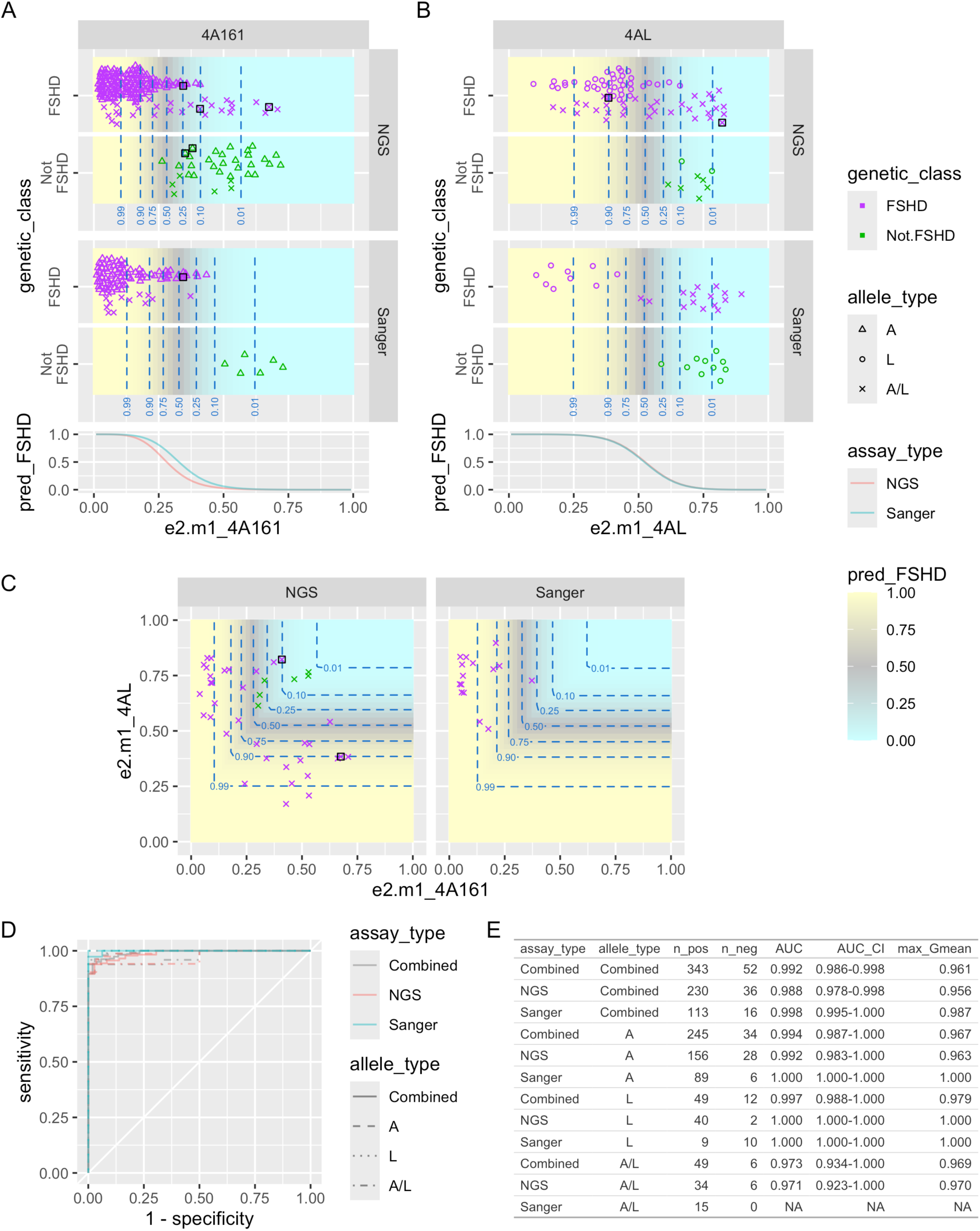
e2.m1 methylation scores for 4A161 and 4AL assays in UNR IRB samples. *(Caption is on next page and applies mutatis mutandis to Figures S1-S7 that show other scores.)* **A**: e2.m1 scores for 4A161 allele (e2.m1_4A161) in genetically FSHD (violet) and non-FSHD (green) samples, divided by assay type (NGS and Sanger). Symbol indicates types of alleles present in the sample: A for only 4A alleles (one or more), L for only 4AL alleles (one or more), A/L for one or more of each. These samples may also have non-permissive 4B alleles and 4A166 alleles (not targeted by BSSA assay and typically not pathogenic even if contracted). Black squares indicate alleles flagged as potentially mosaic. Curves in the bottom panel and yellow-to-cyan gradient in the upper panels show predicted probability of sample being genetically FSHD based on its e2.m1 score for 4A161, from a regularized logistic regression model with an offset used to compensate for class imbalances. The model was fit using only the samples of type A, since for A/L samples one does not know *a priori* which is the disease-associated allele. **B**: Analogous to panel A but for 4AL rather than 4A161 alleles, with the model fit only to the samples of type L. (Note that curves for the two assay types can exactly coincide due to the regularization penalty.) **C**: e2.m1_4A161 (horizontal axis) and e2.m1_4AL (vertical axis) scores for A/L samples for NSG (left) and Sanger (right) assays and predicted probability of a sample being genetically FSHD based on the scores from panel A (for A alleles) and panel B (for L alleles). Dashed blue lines indicate contours of probability 0.01, 0.1, 0.25, 0.5, 0.75, 0.9, and 0.99. **D**: ROC curves showing true positive rate (sensitivity) vs false positive rate (1 – specificity) for classification of samples as genetic FSHD (positive) or not FSHD (negative) based on pred_FSHD scores from A-C. There are separate curves for each combination of allele type (denoted by line type) and assay type (denoted by color), as well as curves for levels of one or both of these factors combined (solid lines for allele types, gray for assay type). **E**: Table of statistics for each of the ROC curves shown in D. These include the number of positive (n_pos) and negative (n_neg) samples used to construct the curve; the area-under-the-curve (AUC); the 95% confidence interval for this area (AUC_CI); and the maximum value of the G-mean (geometric mean of the sensitivity and specificity) for any point on the curve.

The AUCs from the figures above do not depend on a particular cutoff on methylation scores or estimated probability of being genetically FSHD. Overall AUCs (based on the combined allele types and combined assay types) for the seven scores listed above were as follows: 0.987 for mean; 0.934 for Q2; 0.991 for Q1; 0.992 for e2.m1; 0.991 for ex.m1; 0.987 for pct10; 0.969 for pct05; 0.971 for pct03. This supports our primary focus on e2.m1 and Q1 in the current work, which is also consistent with our focus on allele-specific estimated means (akin to e2.m1) and Q1 in our previous work. We note that despite the known weakness of mean for scoring 4A/4A (or 4AL/4AL) samples, its AUC is nonetheless high (lower than for Q1 and e2.m1, but within the 95% confidence intervals for these), which is consistent with the validation of mean methylation from the distal D4Z4 repeat as a highly accurate marker of FSHD disease status and severity in a study from Germany (69). Likewise for Q2, which was used in a study in China that found distal 4A methylation to be highly accurate at distinguishing FSHD from control samples (68). (That study discusses the advantages of Q1 over Q2 for 4A/4A samples, though it is not clear if a hybrid strategy was used for the ROC analysis, with Q2 used to score 4A/4B samples and Q1 to score 4A/4A samples.) In our study the AUC decreases from pct10 to pct05 to pct03, consistent with floor effects as discussed for pct03 above. We include tracks for pct10 and pct05 in many figures as they can help highlight samples that are potentially mosaic for contractions (dropping pct05 for Sanger data, as pct05 is poorly determined with <20 reads).

The value of G-mean shown in the column max_Gmean is based on whatever cutoff on Pr(FSHD) maximizes this score, which need not be 0.5, although using Pr(FSHD) > 0.5 as the cutoff typically resulted in a Gmean nearly as high as max_Gmean (e.g., 0.949 vs 0.957 for Q1 and 0.949 vs 0.961 for e2.m1, both based on the combined allele types and combined assay types). The overall max_Gmean for Q1 is attained at sensitivity = 0.953 and specificity = 0.962 and for e2.m1 it is attained at sensitivity = 0.942 and specificity = 0.981, but in both cases other tradeoffs are possible. The optimal cutoff to use may depend on context, including population demographics of those being assessed, pedigree information, or relative costs of false-positive vs false-negative predictions. Beyond cutoffs, decision-making can depend on the precise estimate of probabilities. Calibration curves indicate that values of Pr(FSHD) over 0.5 in our models may underestimate the actual probability of being genetically FSHD (Figure S9). Using a milder regularization penalty (lambda.min rather than lambda.1se) appears to provide better calibration overall, with the caveat that there were few samples with intermediate predicted probabilities when using milder regularization, limiting confidence in the calibration at intermediate values even if this is suggested by the LOESS curve (which uses default span of 0.75). The milder regularization penalty lambda.min also gave slightly better AUC and G-Mean scores than lambda.1se (Figure S10 vs Figure 10). Except where noted, the results discussed herein are for lambda.1se, as it in principle provides an extra hedge against model overfitting (beyond the cross-validation used to select lambda.min), but it appears that this may be over-cautious here. Over-moderation of predicted probabilities using lambda.1se may also be exacerbated if one adjusts the offset term in the fitted models to specify a smaller prior probability of being genetic FSHD. For these reasons we provide coefficients for models using lambda.min as well as lambda.1se; regardless of which approach one uses, models can be recalibrated as more data becomes available.

As discussed in the Methods section, we do not directly report overall accuracy simply because it depends on relative sizes of the FSHD and non-FSHD classes and can be misleading when there is strong imbalance in class sizes, which is the case for the UNR IRB study. However, the overall accuracy is a weighted average of sensitivity and specificity (which are both between sqrt(G-mean) and 1.0), so when both sensitivity and specificity are high, the overall accuracy will be high regardless of relative class sizes.

There are several caveats to the interpretation of these AUC and G-mean scores, which should be considered if comparing results from this study to results from other studies:

1)Although scores are provided separately for allele types A, L, and A/L, the scores for those three allele types combined are based on the relative proportions for these particular subjects, and their relative frequencies can be quite different in different populations: the 4A161L allele is common in European populations (∼20% as frequent as 4A161S alleles) but is absent in the Asian and African HAPMAP populations (70), and not observed in recent FSHD D4Z4 methylation studies in China (68, 71). However, because the classification performance based on AUC and max_Gmean was similarly high regardless of allele type, the relative haplotype frequencies may have only a mild impact on overall performance. (Note that AUCs are less precisely determined for allele types with fewer samples.)
2)We did not include subjects with 4B/4B genotypes in evaluating the overall performance. If such subjects are regarded (by definition) as not being genetically FSHD, then they would all be classified correctly and their inclusion would increase the overall AUC and G-mean. Such subjects may account for roughly 25% of the overall population if (as is often stated) 4A alleles are roughly as common as 4B alleles, although the frequency varies by population (33).
3)The “ground truth” genetic FSHD status is based on heterogeneous genetic testing, and some test reports may have errors, due for example to unusual D4Z4 array configurations that are blind-spots for standard PFGE testing. The genetic status of some participants may also be inherently ambiguous due to mosaicism for a contracted allele, and in some cases the clinical status of individuals with negative FSHD genetic tests is unknown.
4)The participants in the UNR IRB study do not represent random samples from the populations of either FSHD or non-FSHD subjects. Nor are they independent samples from these populations, as multiple members of some families participated in the study, and this may potentially lead to an enrichment of participants with unusual or complex genetics. These factors may impact both the AUCs and the widths of the confidence intervals for them. (We do not report detailed pedigree information for the participants, nor their ages and sexes, to reduce the risk of re-identification.)
5)One limitation of this analysis is the lack of borderline D4Z4 deletion cases (e.g., 11 or 12RUs), as there is some debate if these could in fact represent FSHD1 – i.e., result in FSHD in the absence of an FSHD2-associated mutation in *SMCHD1*, or more rarely in *LRIF1* or *DNMT3*, or in unknown factors that similarly result in decreased methylation for all D4Z4 arrays (suggestive of idiopathic FSHD2). Since the goal of this study was to validate this epigenetic diagnostic procedure and the D4Z4caster program, we used cohorts with more clearly defined FSHD genetics. Analysis of ambiguous borderline genetic FSHD1 cases with clinical evaluations will be included in a follow-up study.

For each score and assay type (NGS or Sanger), the coefficients for the logistic regression models for 4A and 4AL alleles (which are for logit-transformed input; see Methods) are given in Table S5, along with methylation values corresponding to estimated Pr(FSHD) of 0.25, 0.5, and 0.75. Figures 11, 12 and 13B collectively show all the samples from the IRB study (aside from two Sanger samples with unknown clinical status), separated into samples of known genetic status with NGS data (Figure 11), samples of known genetic status with Sanger data (Figure 12), and samples of unknown genetic status with NGS data (Figure 13B). In these figures, the score corresponding to Pr(FSHD) = 0.5 is denoted by a dashed line, red for 4A and blue for 4AL. The interval from Pr(FSHD) = 0.25 to 0.75 is shaded, again red for 4A and blue for 4AL, but in both cases representing a “gray area” in which predictions of genetic FSHD status based on the methylation score are not confident one way or the other. For the BSSX track, the dashed green line is at 30% mean methylation, an approximate threshold beneath which individuals are predicted to have FSHD2 if they are predicted to have FSHD based on 4A- or 4AL-specific methylation levels in e2.m1 or Q1 tracks. There is again expected to be a gray area for such predictions, although in this study we do not attempt to provide precise estimates for Pr(FSHD2) using logistic regression because the ground truth for many subjects is not known: for example, subjects with clinical FSHD that are found by genetic testing to have contracted D4Z4 alleles consistent with FSHD1 may not be subsequently tested for mutations in *SMCHD1* (or *LRIF1* or *DNMT3*), making it unknown which are genetically FSHD1 + FSHD2.

**Figure 11:**
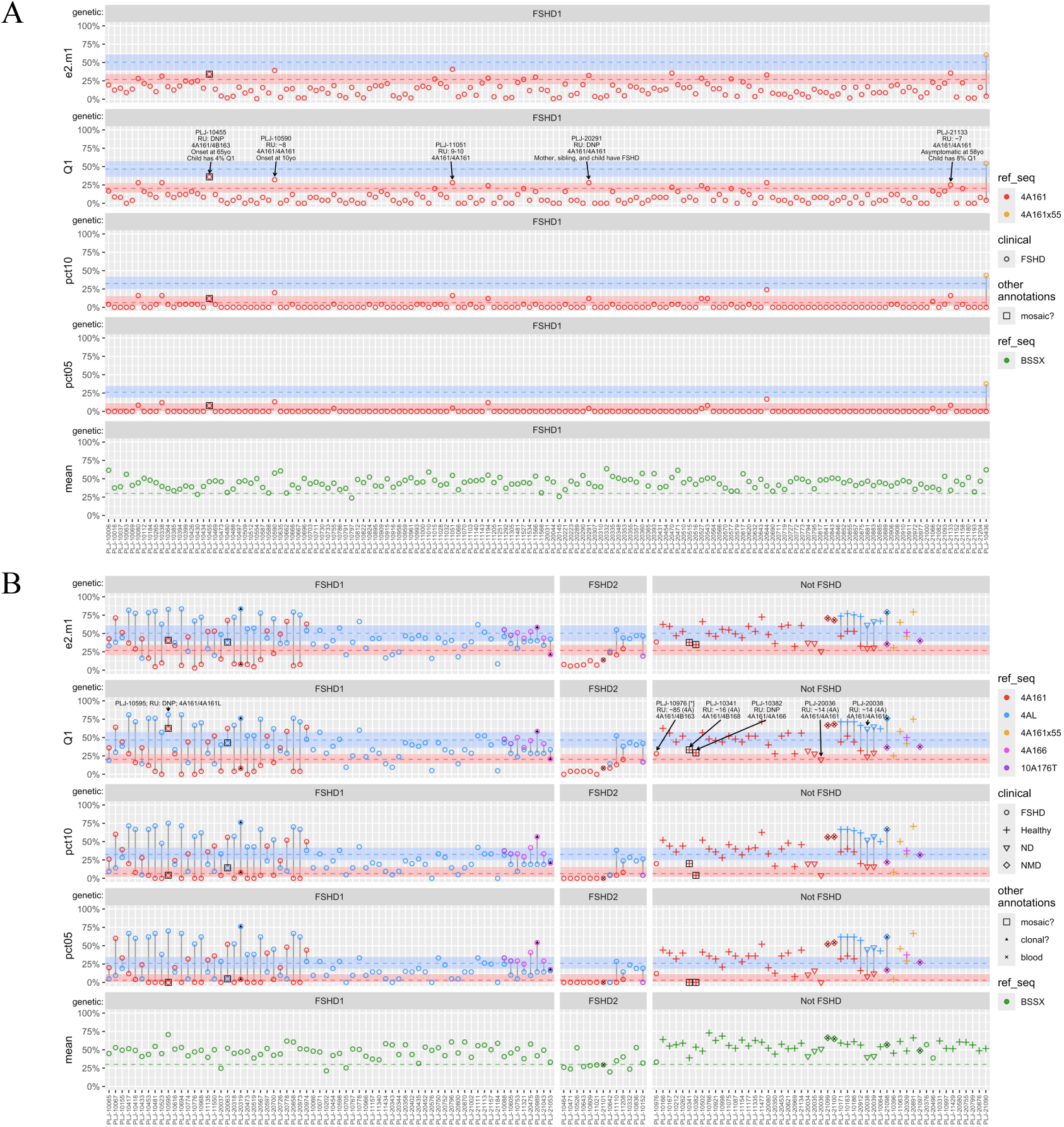
Methylation data for samples from the UNR IRB study with known FSHD genetic status from NGS assays. Samples are from saliva except for a few from blood (small black x; see legend). Samples are grouped based on genetic diagnosis (FSHD1, FSHD2, Not FSHD), and shapes indicate clinical group (FSHD, healthy, NMD=other neuromuscular disorder). Colors denote reference sequences. Dashed blue and red horizontal lines indicate cutoffs for classifying subjects as FSHD based on corresponding 4A161 and 4AL scores, respectively, and shaded blue and red regions indicate the “gray area” of 25%-75% confidence in classification. Dashed gray line for mean BSSX score is at 30%, an approximate cutoff for FSHD2 for subjects that are also below FSHD cutoffs for a 4A161 and/or 4AL allele. **A:** FSHD1 samples with 4A161 but not 4AL reads. **B:** All other samples. Some of the labeled samples in the Q1 track for A and B are discussed in the main text.

**Figure 12:**
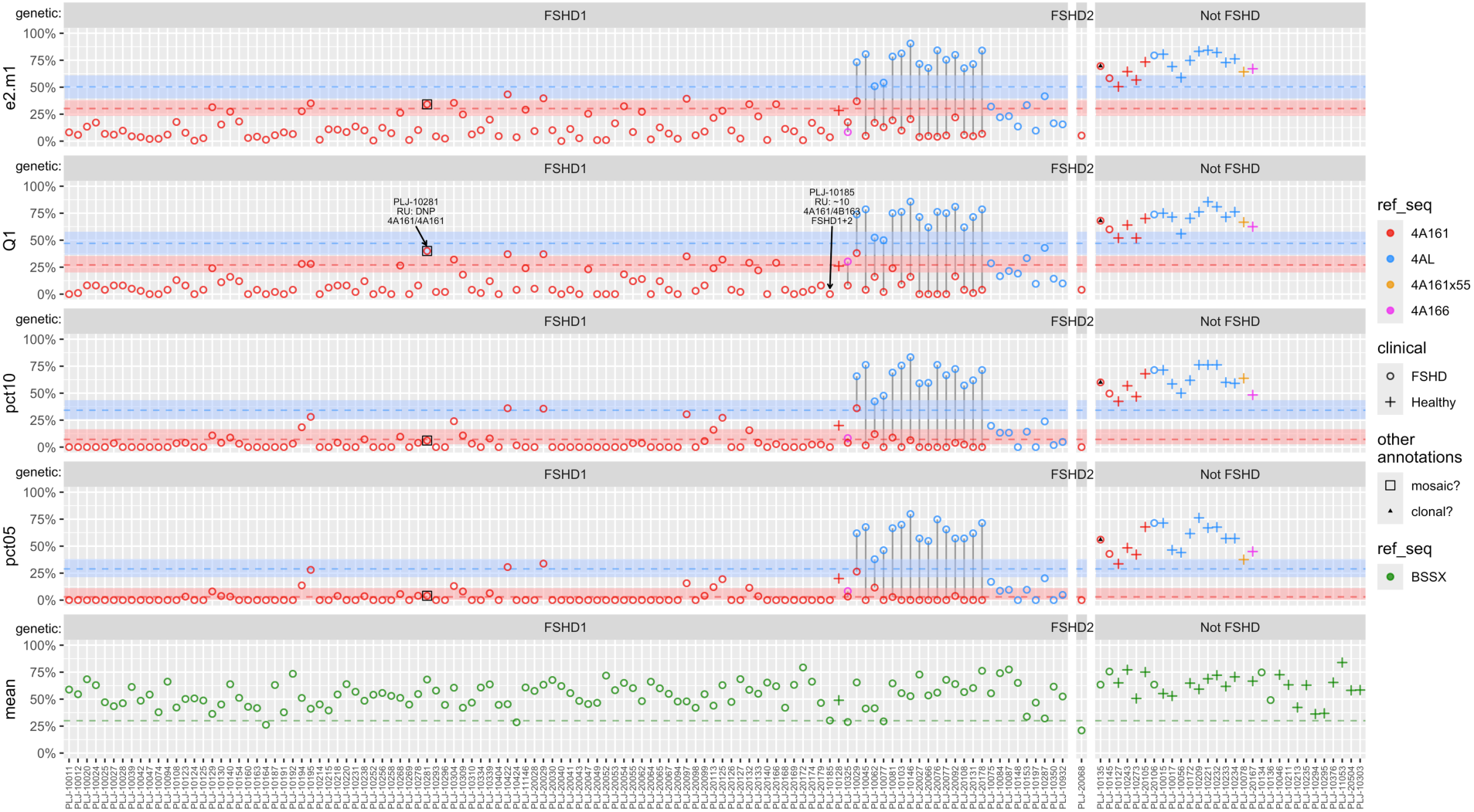
Methylation data for samples from the UNR IRB study with known FSHD genetic status from Sanger assays. See caption of Figure 11 for details.

**Figure 13:**
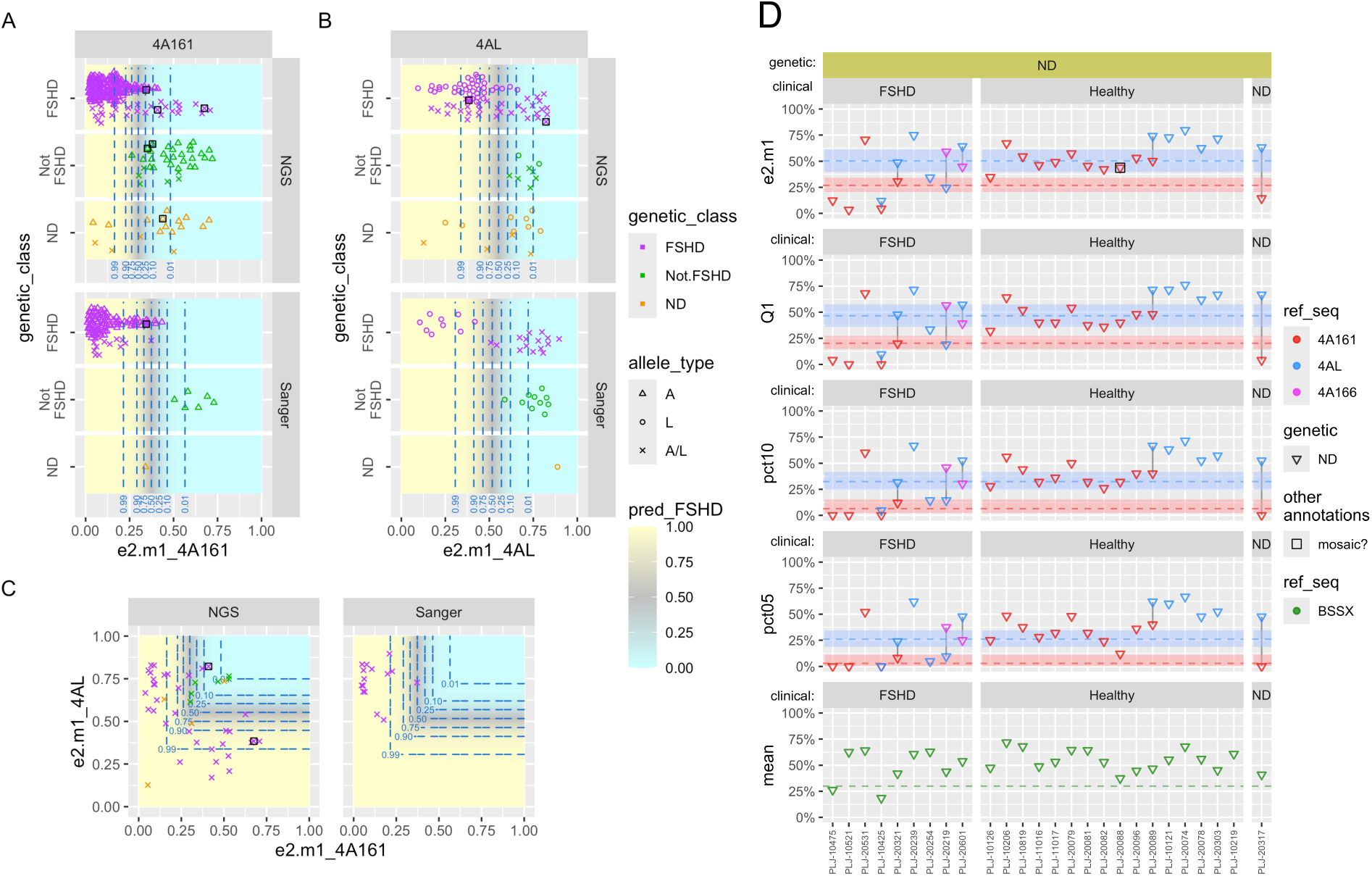
Methylation data for samples from the UNR IRB study with unknown FSHD genetic status. All are from NGS assays on saliva samples. Panels **A-C** are similar to panels A-C in Figure 10 but also show genetic ND (not determined) samples (orange). Panel **D** shows details of the genetic ND samples with NGS data. Colors denote reference sequences and classification cutoffs, as described for Figure 11; samples are grouped based on clinical diagnosis (FSHD1, healthy, or ND); and symbol shapes indicate genetic diagnosis, which is ND for all these subjects. Details of the two Sanger ND samples (one from panel A and one from panel B) are not shown in D, but both are clinically healthy.

Consistent with the high AUCs and G-means, most samples in Figures 11 and 12 are on the expected side of the cutoffs given their genetic status, and when this is not the case they are often in the gray area is which predictions are not confident. Note that for FSHD samples with both 4A and 4AL alleles (i.e., A/L samples), only one of the alleles needs to be below the cutoff for that allele for the predicted genetic class to be correct when using max(PA, PL) for scoring the A/L samples. In these plots samples that are flagged for being potentially mosaic (Figure 9) or for having potentially severe clonal artifacts (Figure S11) based on larger- or smaller-than-usual differences between certain percentiles (see Methods and Figure 6) are annotated with special symbols. In the Q1 track several samples with incorrect or borderline predicted genetic status are labelled by name, accompanied by brief clinical or genetic details for context. These include:

1)Several of the samples predicted to be mosaic (PLJ-10455, PLJ-1095, PLJ-10341, and PLJ-10382, which was also the outlier in the NGSx3 validation plots in Figure 8). Although we do not have direct experimental verification that these samples are mosaic (using for example optical genome mapping or nanopore sequencing, both of which would require HMW gDNA), there is indirect evidence in support of mosaicism for several of the flagged samples. For example, subject PLJ-10455 has disease onset at age 65 and 36% Q1 4A161 methylation but has a child with 4% Q1 4A161 methylation, which is consistent with the child inheriting a contracted allele that is mosaic in the parent. Or, in the case of PLJ-1095, which has both 4A and 4AL alleles, the 4A methylation grid plot shows missing methylation calls for CpG 44 in almost all of the reads, both hypomethylated and hypermethylated (Figure 9). Based on the bisulfite sequencing reads, this appears to be due to a rare *DUX4* intronic G>A variant at chr4:190175512 (GRCh38.p14; rs1415465659) that changes CpG 44 to CA. The rareness of this variant, which has allele frequency < 0.001 in each genetic ancestry group in gnomAD 4.1.0 (72), together with the strongly bimodal methylation pattern, suggests that the reads arise from two alleles that differ in the number of D4Z4 RUs but share the same distal repeat sequence. Although a partial *cis* duplication of the 4A array could also explain a rare variant being present in distal repeats for both contracted and noncontracted arrays, reads from the distal repeats would be expected to occur in roughly equal frequencies in this scenario, whereas the 4A methylation data for PLJ-1095 suggests the hypomethylated and hypermethylated reads have frequencies of ∼15% and ∼85%, respectively, which is consistent with mosaicism.
2)Several samples with 4A methylation in the gray area that are classified as genetically non-FSHD based on having a 4A allele of ∼14RU, but for which the clinical status is not known (PLJ-20036 and PLJ-20038). Whole exome sequencing did not identify pathogenic mutations in known FSHD2 genes, and the mean BSSX methylation levels are not strongly suggestive of FSHD2.
3)Sample PLJ-10976 (marked with an asterisk), which has Q1 for 4A just above the gray area and mean BSSX methylation just above the 30% threshold. This subject is the only genetically non-FSHD subject in Figure 11B that is reported as being clinically FSHD (although as noted above several subjects with 14RU 4A arrays have unknown clinical status). This subject has a *SMCHD1* c.769del mutation and a contracted 4B allele of 8RU but is considered genetically non-FSHD since the shortest permissive 4A or 4AL allele is 85RU, well above the semi-short array lengths typically seen for FSHD2. This subject also has a rare genetic disorder (not FSHD but not disclosed here to reduce risk of re-identification) for which muscle weakness is one of the symptoms. This raises the possibilities that overlapping symptoms confounded the clinical diagnosis of FSHD or that the other disorder resulted in FSHD symptoms manifesting in a genetic context in which they ordinarily would not, as with the frequent reports of “double trouble” conditions that modulate clinical severity of FSHD (73).

Figure 13 highlights samples from the UNR IRB study that have unknown FSHD genetic status, including them as a separate group (ND) in panels A-C, which are otherwise the same as Figure 10 A-C. Panel D shows that in most cases the predicted genetic status of these individuals is consistent with the clinical status, though there are two strong exceptions to this, PLJ-20351 and PLJ-20239, which could potentially represent clinical misdiagnoses of FSHD.

### Correlation between methylation and repeat lengths

A strong positive correlation between number of 4A RUs and 4A methylation is well-established in comparisons that include both genetic FSHD and non-FSHD subjects, but this can be driven largely by subjects with FSHD1 typically having both shorter 4A arrays and lower methylation levels than subjects without FSHD (12, 13, 74). Correlation between number of RUs and methylation just among subjects with FSHD1 is a more delicate matter. Estimates based on methylation-sensitive restriction sites proximal to the D4Z4 array give averages over both 4q alleles, or both 4q and also both 10q alleles; these average are broadly consistent with methylation increasing with array lengths, and deviations from the expected average given the array lengths (Delta1 score) can be computed if all array lengths are known (12), but this does not establish what proportion of the deviation is on the pathogenic contracted D4Z4 array. Interpretation can also be complicated for methylation estimates based on bisulfite sequencing of the DR1 region within each D4Z4 RU (40). This again represents an average over alleles, but an average that is weighted by the number of RU in the arrays. This can confound direct comparisons between DR1 methylation levels for FSHD1 subjects with different number of RUs (75), as shorter contracted alleles get less weight in the weighted averages. Recent studies using Nanopore sequencing give allele-specific methylation estimates by sequencing entire D4Z4 arrays (39, 76) but these require HWM gDNA.

Our BSSA and BSSL assays measure methylation on only the distal repeat, and unlike the methylation-sensitive restriction site assays that assess single CpGs, bisulfite sequencing of multiple CpGs allows for allele-specific estimates based on mixture models even when there are two alleles of the same haplotype. Here we computed correlations with the number of RUs only for the BSSA assay, which picks up 4A161-type but not 4AL alleles, as there were many more subjects with 4A161-type alleles. Subjects with both 4A and 4AL alleles were excluded since for them the 4AL rather than 4A allele may be contracted. We observed a moderate positive correlation (Pearson correlation coefficient r = 0.38-0.40) between number of RUs for the contracted 4A allele and its allele-specific estimated methylation score (e2.m1) among the FSHD1 subjects for which the RU data was available (Figure S12). Correlations were similar for subjects with two 4A alleles (group AA: r = 0.40; p = 0.03; n = 29), subjects with one 4A allele and one 4B or 4A166 allele (group An: r = 0.39; p = 0.02; n = 35), and both those groups combined (r = 0.38; p = 0.002; n = 64). We also used a linear model to fit e2.m1 to the number of RUs and an additive categorical factor for group (An vs AA). The coefficient for RU was 1.46 (p = 0.0015), indicating an average increase of ∼1.5% methylation per RU within the limited range of ∼1-10 RUs for subjects with FSHD1. (Over the full range of 1-100 or more RUs a nonlinear relation is expected as methylation levels are bounded above by 100%). The intercept was 6.5, which in this parametrization is the estimate for the AA group. The coefficient for An vs AA was -3.78, indicating lower scores for samples with one 4A allele than samples with two 4A alleles on average (while adjusting for the number of RUs), but this difference was not significant (p = 0.08). In contrast, for the mean methylation score the coefficient for An vs AA was -8.80 and was highly significant (p = 0.00007). This is consistent with the estimated allele-specific methylation e2.m1 removing much of the confounding effect of methylation on the non-contracted allele. (The correlations above used fractional RUs when converting from restriction fragment lengths in kb, but results were similar with RUs rounded to the nearest integer or rounded down.)

We next used the cutoffs and gray areas defined based on the UNR IRB study to evaluate methylation data from several additional studies. The samples in the UNR IRB study were predominantly from saliva, with just a few from blood (as indicated in Figure 11), and samples from some of the studies below are from fibroblasts, myoblasts, PBMCs, and muscle biopsies. Our previous work shows that D4Z4 methylation levels are broadly similar for saliva, PBMCs, and cultured muscle cells (13, 41), although it is possible that the cutoffs, calibrated almost entirely on saliva samples, will not be optimal for other sample types. D4Z4 methylation data for some of these studies have been previously published but here they are uniformly re-analyzed using D4Z4caster. Results for each study are shown in figures that are similar in format to Figures 11, 12 and 13B for the UNR IRB study, but with different study-specific groupings and subgroupings of samples.

### BiLat study

Figure 14 shows D4Z4caster results for samples from the BiLat study, which compares left and right tibialis anterior (TA) muscle biopsies from 32 subjects, all with genetically-confirmed FSHD. Sample metadata is in Table S1; D4Z4caster results are in Table S4. Data from the same NGS runs of these samples has been published previously (67). Whereas the previous study reported only per-sample median methylation levels of the assayed CpGs, here we present the results of analyses using D4Z4caster, which include allele-specific methylation estimates and predictions of genetic FSHD status based on methylation scores. As in the previous study, we found strong concordance between scores from left and right TA biopsies from the same subject: the mean absolute value of the difference in scores for paired bilateral samples ranged from 1.5 to 6% methylation depending on assay type (4A161, 4AL, BSSX) and score type (e2.m1, Q1, pct10, mean). D4Z4caster correctly predicted all samples to be from genetically FSHD individuals based on both the e2.m1 and Q1 methylation scores, with none even in the gray area. It should, however, be noted that there are no non-FSHD muscle biopsies for comparisons here, and it is possible that methylation scores are systematically lower for muscle biopsies than for saliva or blood specifically for subjects with FSHD or regardless of FSHD status. If skeletal muscle does have particularly low D4Z4 methylation levels, this may provide one explanation as to why, in FSHD, both *DUX4* expression and pathology are most prominent in skeletal muscle. In future studies we plan to compare methylation levels from muscle biopsies to those from saliva in the same subjects to directly address this.

**Figure 14:**
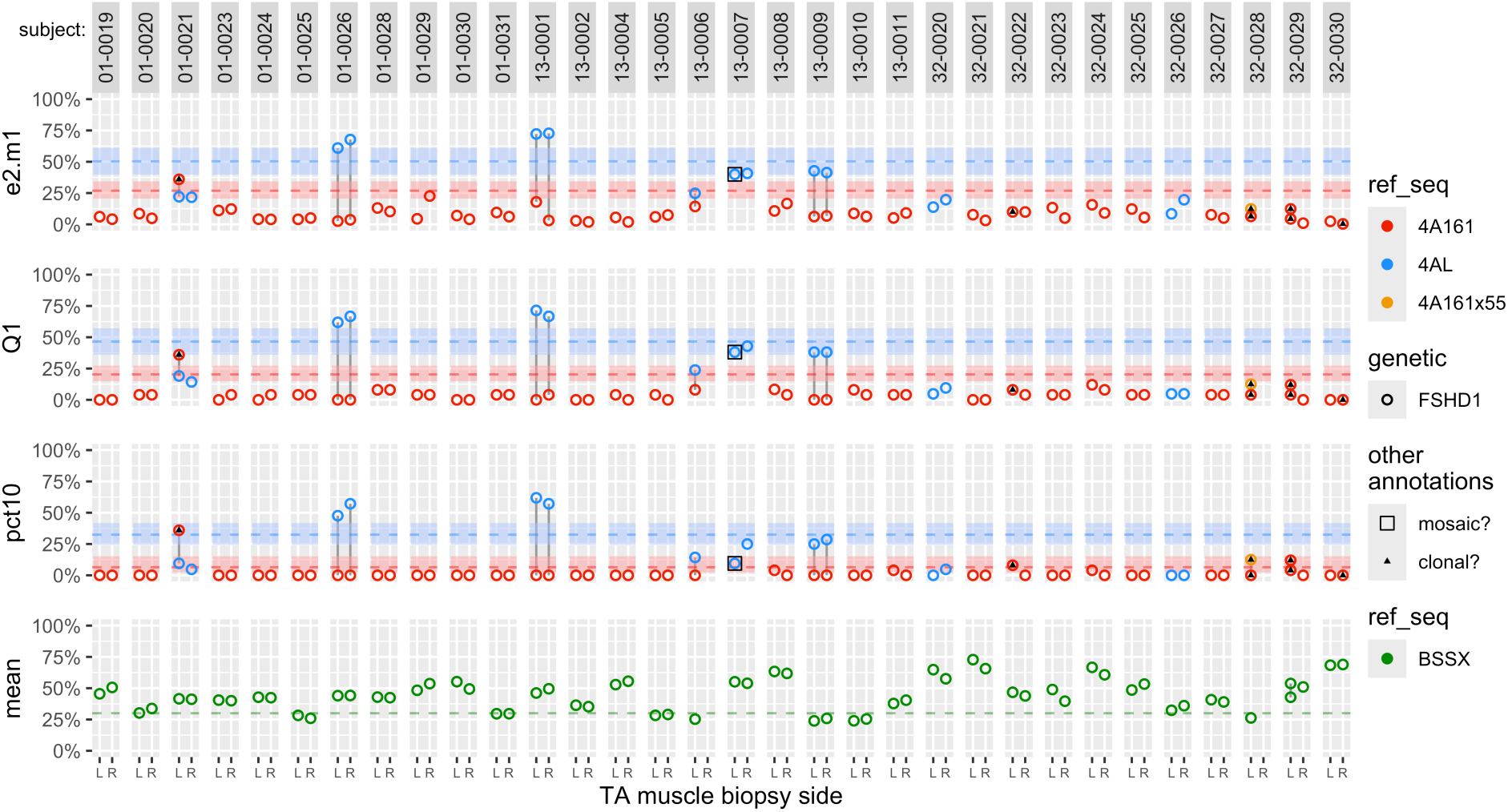
Methylation data for samples in the BiLat study. All are based on NGS assay. Colors denote reference sequences and classification cutoffs, as described for Figure 11. All samples are from subjects (32 total) with genetic FSHD1. Samples are grouped by subject (labeled at top of plot) and for each subject the methylation level from bilateral muscle biopsies of tibialis anterior (TA) are shown (L for left, R for right at bottom of plot). Subjects 13-0006 and 32-0028 have data only from the left side and subject 32-0029 has data from two samples from the left side, from different sections of the same biopsy. Small black triangles indicate assays that were flagged for potential clonal artifacts, possibly due to relatively low input DNA from the muscle biopsy samples. The mean absolute value of the difference in scores for paired bilateral samples ranged from 1.5 to 6.1 (on the scale of % methylation) depending on assay type (4A161, 4AL, or BSSX) and score type (e2.m1, Q1, pct10, or mean). (Only the first of the two L samples for 32-0029 was used in this calculation, 32-0029L48.) Data from the same NGS runs of these samples was presented in an earlier report (as per-sample median methylation) (Ref 67).

The samples in the BiLat study also have a higher rate of assays being flagged as having potentially severe clonal artifacts (indicated by small black triangles in plots) than samples in the UNR IRB study. This may be due to the smaller amount of input DNA being used for muscle biopsy samples. For saliva samples, the flagged assays were typically for alleles that were not targeted for PCR amplification (e.g., 4A166 alleles that may be amplified with low efficiency with the BSSA assay if there are no 4A161 alleles present) or alleles that are targeted but not predicted to be present (e.g., reads for 4A161 in samples with predicted 4AL/4A166 haplotypes). Figure 12B has examples of both. For the BiLat study there were flagged assays of this type, which can be regarded as likely irrelevant (as with the examples above), but there was also several flagged BSSA assays for samples in which 4A161 alleles are predicted to be present (such as 32-0022L and 32-0030L) based on genetic test results and/or PCR. Methylation scores for the flagged left TA samples were typically similar to the (unflagged) paired right TA samples, so these may represent examples in which clonal amplification did not necessarily introduce a strong bias in scores. If the chromosomes that are clonally amplified are randomly selected then the estimated mean methylation is not biased at all due to the linearity of expectations, though this does not apply for percentile-based scores or e2.m1, and even for mean methylation the variance of the estimate is higher (potentially much higher) as it is based on fewer independently-sampled chromosomes. For these reasons, caution is recommended in interpreting scores for flagged assays.

### Wellstone MDCRC study

Figure 15 shows D4Z4caster results using samples from a UMass Medical School Wellstone MDCRC study, in a reanalysis of previously published Sanger sequencing data, with myoblast and PBMC samples (13). Sample metadata is in Table S1; D4Z4caster results are in Table S4. The exact scores differ somewhat from those published previously due to three factors: 1) Here scores are based on just the 3’ CpG sites that typically have high coverage with NGS, rather than all CpGs in the amplicons, for consistency with the NGS scoring; 2) Here we used D4Z4caster rather than BISMA (77) (for which the webserver appears to be long-defunct) for aligning and filtering of the Sanger reads; 3) Here we used an MLE approach (implemented as part of D4Z4caster) rather than a full Bayesian approach for estimating the parameters of the beta-binomial mixture models (for e2.m1 and ex.m1). Results are described further in the legend to Figure 15.

**Figure 15:**
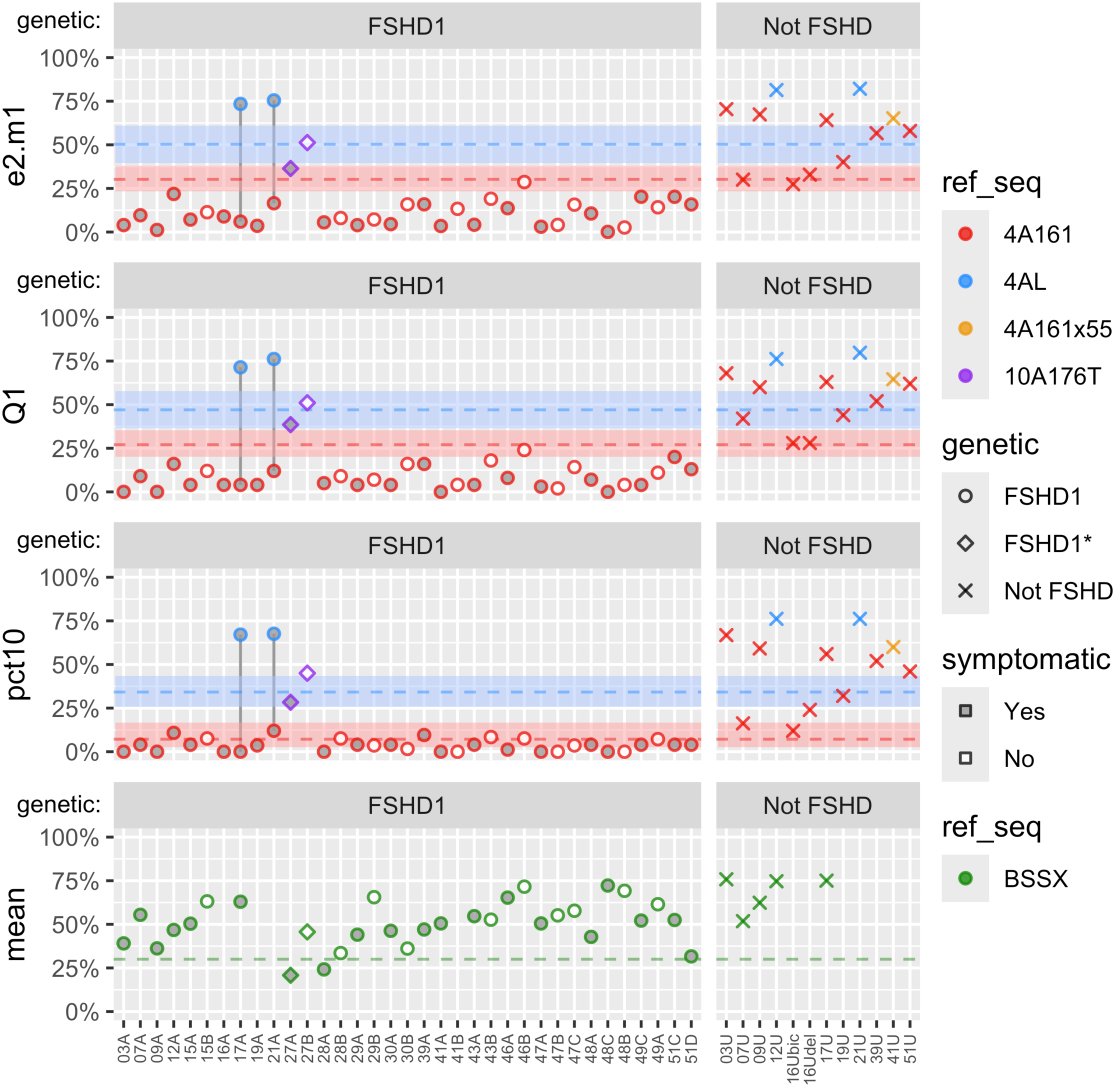
Methylation data for the Wellstone MDSRC samples. All are based on Sanger assays. Samples with IDs beginning 03-30 are from myoblasts derived from muscle biopsies (biceps, aside from 16U which has samples from both biceps and deltoid). Samples with IDs beginning 39-51 are from PBMCs. Colors denote reference sequences and classification cutoffs, as described for Figure 11. Samples are grouped based on genetic diagnosis (FSHD1 or Not FSHD) and ordered by sample name within groups so that FSHD samples from the same family (numeric prefix in sample name) are adjacent. Symbol shapes indicate genetic diagnosis, with FSHD1* indicating two likely misdiagnoses (27A and 27B, which appear to have contracted non-permissive 10A176T alleles), and hollow symbols indicate genetic FSHD1 cases that were asymptomatic. Note that such cases typically have higher 4A161 methylation scores than the symptomatic family members to their immediate left, as we previously noted for these same samples (Ref 13). BSSX assays were not done for several of these samples, so there are some missing scores in the mean BSSX panel.

### Arhinia and/or FSHD2 study

Figure 16 shows D4Z4caster results for samples provided by the University of Rochester Medical Center, University of Iowa Muscular Dystrophy Specialized Research Center, and Boston Children’s Hospital. Sample metadata is in Table S1; D4Z4caster results are in Table S4. The primary focus is a comparison of D4Z4 methylation profiles for subjects with Bosma arhinia microphthalmia syndrome (BAMS) and subjects with FSHD2, as both are associated with mutations in *SMCHD1* (78, 79). The spectrum of pathogenic *SMCHD1* mutations observed for the two disorders is quite different, with mutations for FSHD2 (missense and nonsense variants, splice-site variants, and indels) broadly distributed across the SMCHD1 protein, whereas mutations for BAMS (exclusively missense variants) are largely restricted to the extended ATPase domain (80). There is, however, at least some overlap, as at least two *SMCHD1* mutations have been implicated in both disorders, one in different individuals and the other in the same individual (79). Notably, BAMS does not have the same requirement for a permissive 4A or 4AL allele that FSHD2 does (whether semi-short or not). In Figure 16, subjects with BAMS that have FSHD-permissive alleles are listed as genetically Arhinia/FSHD2, although the clinical diagnosis for all of them is BAMS without mention of FSHD2. Subjects in the genetically Arhina, Arhinia/FSHD2, and FSHD2 groups all show hypomethylation of the D4Z4 arrays based on the BSSX assay, with mean methylation typically below, or in a few cases just slightly above, the ∼30% cutoff for FSHD2 mentioned previously. This is consistent with prior reports of hypomethylated D4Z4 arrays in BAMS (79, 80), although because the samples shown in Figure 16 overlap those in prior reports, the purpose here is not independent validation but rather presentation of the methylation profiles using the same uniform D4Z4caster analysis as for the other studies presented here.

**Figure 16:**
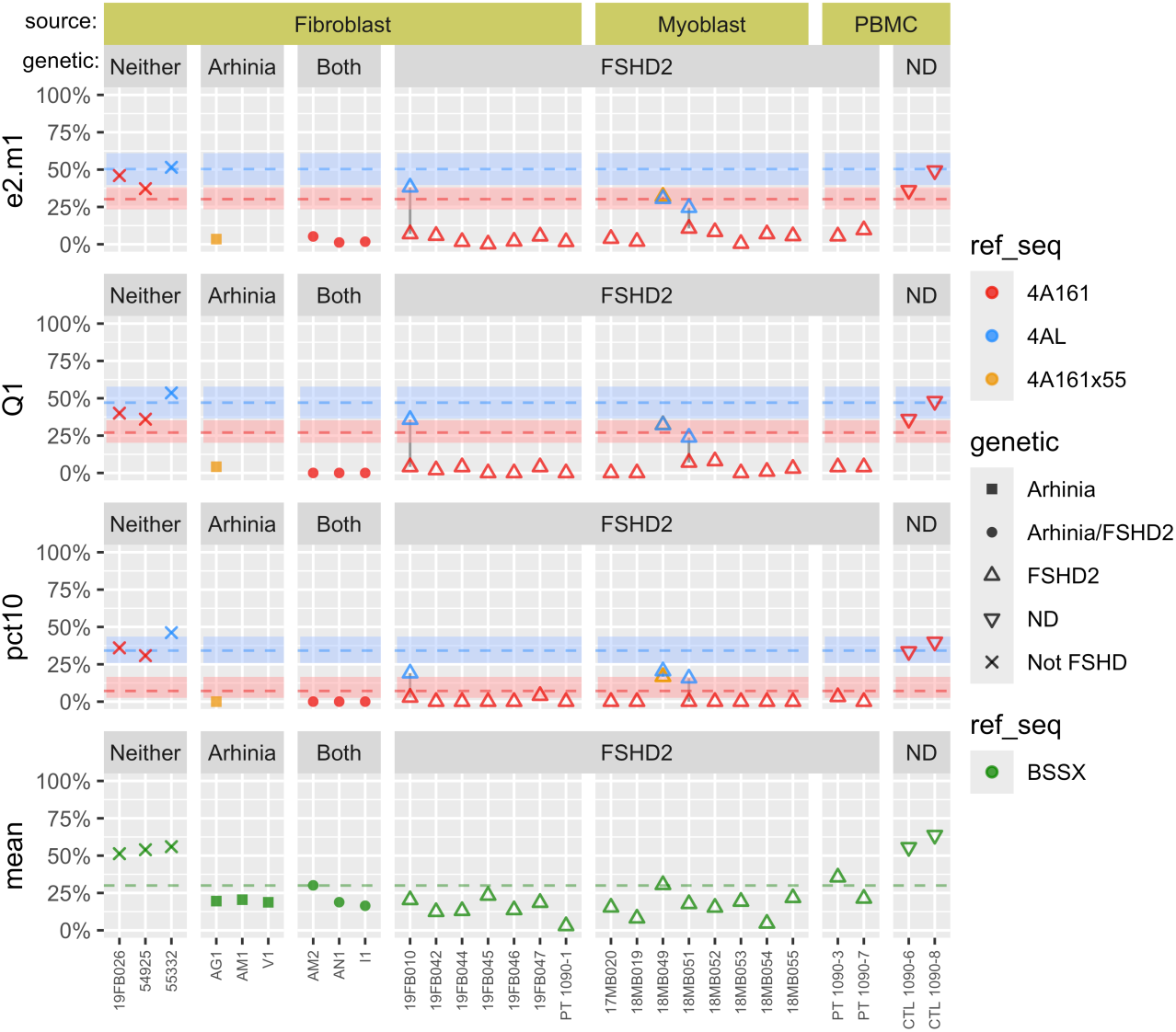
Methylation data for Arhinia, FSHD2, and control samples. All are based on Sanger assay. Colors denote reference sequences and classification cutoffs, as described for Figure 11. Samples are grouped based on sample source (Fibroblast, Myoblast, and PBMC) and by genetic group (Arhinia, FSHD2, both, neither, or ND [not-determined]). Symbols also denote genetic group. “Not FSHD” (“neither”) samples do not have data provided on *SMCHD1* mutations but are clinically healthy. Note that the three samples indicated as having genetic Arhinia but not Arhinia/FSHD2 do not have any points shown for the 4A161 or 4AL reference sequences in the upper three panels, which is as expected since two (AM1 and V1) have 4B/4B haplotypes and the third (AG1) has 4A166/4B haplotypes (here with the 4A166 allele being picked up as 4A161x55 rather than 4A166).

## Discussion

FSHD is considered the third most prevalent muscular dystrophy (2). However, due to complications regarding genetic diagnostics, the true prevalence is not known. Many clinically affected individuals don’t have access to genetic testing. In addition, there are many asymptomatic and disease nonmanifesting individuals, both within FSHD families and in the general population, who meet the genetic criteria for FSHD1, suggesting there is a large unknown at-risk population (9–12, 14, 81). Interestingly, some clinical FSHD cases do not meet the currently accepted genetic criteria for FSHD1 or FSHD2 or display global reduction of D4Z4 methylation suggestive of idiopathic FSHD2 yet may in fact be true FSHD cases due to currently unknown epigenetic modifiers. Better access to accurate FSHD molecular diagnostics that can identify all forms of FSHD is critically needed.

FSHD1 accounts for >95% of genetically confirmed FSHD (5, 82). Current clinically accepted FSHD1 diagnostic techniques all measure the physical sizes of the chromosome 4q35 and 10q26 D4Z4 arrays and thus require specially prepared HMW gDNA typically isolated from PBMCs (4, 34, 37, 83). Recent long-read DNA sequencing technologies being developed for FSHD diagnostics have similar requirements (36, 38, 39, 76). While new technologies for measuring D4Z4 repeats, and including haplotype information, offer some improvements to diagnostic workflow and can be more readily established in a standard diagnostic lab setting compared with traditional pulse-field gel electrophoresis (PFGE) followed by Southern blotting (34), they do not overcome a key limitation to wide accessibility and utility: the requirement for HMW gDNA specifically prepared for this type of testing. Typical processing of DNA for candidate gene panel arrays, whole exome sequencing (WES), and whole genome sequencing (WGS) commonly used in neuromuscular disease (NMD) diagnostic labs is not appropriate for FSHD1 testing and a specialized workflow starting with fresh DNA isolation and processing is required. Similarly, biobanked gDNAs cannot be re-analyzed for FSHD1 by these technologies. Additional concerns include the requirement for nonstandard equipment for analysis and cost per sample analyzed. The situation for FSHD2 is somewhat different. Although genetic mutations in *SMCHD1* or other known FSHD2 genes can be identified by candidate gene panel, WES, and WGS approaches, FSHD2 also has two genetic requirements at the D4Z4 locus: 1) an FSHD-permissive 4qA or 4qAL allele that is 2) in *cis* with a D4Z4 RU that is typically ∼20RUs or less (21). Thus, even for individuals with FSHD2-associated mutations in *SMCHD1* or other genes, a genetic diagnosis of FSHD2 (or a prognosis for clinical FSHD) still requires information about the D4Z4 locus for which the limitations of FSHD1 genetic testing still apply.

However, since all forms of FSHD are caused by the epigenetic dysregulation and subsequent aberrant increased expression of the *DUX4* gene from a distal 4q35-derived D4Z4 RU with a *cis* distal FSHD-permissive haplotype, we hypothesized that assaying the epigenetic status of this distal region would be diagnostic for genetic FSHD and those at risk for developing clinical FSHD. A BS-PCR approach proved useful for identifying FSHD2 patterns of D4Z4 hypomethylation (40), but not for FSHD1, which is epigenetically distinct (5, 74). Therefore, in 2013 we established a set of targeted BS-PCR assays to specifically determine the methylation state of the relevant FSHD-permissive distal 4q35-derived D4Z4 RUs (13, 41, 45). Significantly, this approach has been performed successfully on gDNA, isolated using standard methods, from saliva samples, PBMCs, cultured myoblasts and fibroblasts, and muscle biopsy samples, including both frozen biobanked gDNAs and gDNAs isolated from organic tissue extractions (13, 41, 42, 67). Here, we have validated our targeted BS-PCR FSHD analysis against several large genetically confirmed cohorts representing diverse populations and show that it very accurately identifies subjects that fit the current genetic definitions of FSHD1 and FSHD2 and clearly distinguishes them from control non-FSHD subjects. Importantly, this protocol separately analyzes the epigenetics of 4A (short) and 4AL alleles, which have very different methylation profiles in FSHD (likely due at least in part from testing different CpG sites). In addition, we have improved the workflow, throughput, and data analysis for this technology with the development of the D4Z4caster program for transition to a clinically relevant NGS diagnostic setting.

Since our 2014 report documenting that epigenetic analysis using targeted BS-PCR was apparently diagnostic for FSHD1 and FSHD2, others have adopted our approach (68, 69) and an alternative BS-PCR method has been developed, albeit focused on a different region of the FSHD locus, the 4q35 PAS (46, 84). The PAS BS-PCR assay has the advantage of directly assessing only FSHD-permissive chromosomes; however, this strategy assesses combined total methylation of the sample being assayed at two individual CpG sites and does not distinguish individual chromosomes or the two 4q alleles when both are permissive, nor does it distinguish between 4A and 4AL. In addition, permissive alleles containing SNPs that do not typically result in FSHD even if contracted, such as some 4A166 sub-haplotypes, are not excluded from the analysis, thus complicating the interpretation (30). Regardless, the PAS assay has proven fairly accurate for distinguishing FSHD from non-FSHD when tested against defined cohorts (71, 84). However, the limitations to this approach become more apparent when performing large population screening or using epigenetic status for prognostic purposes, when obtaining an accurate epigenetic assessment of the pathogenic allele(s) is critical (85, 86).

Even within the framework of our general BS-PCR approach (whether assessing exactly the same CpG sites or not) there are decisions to be made on how to transform the resulting methylation data (bisulfite sequencing reads) into classifications or predictions for FSHD diagnosis or prognosis. In the Results section we outline the conceptual pros and cons of different ways of distilling methylation data into scalar scores that can be used for classification or prediction. This includes scores we used in our previous Sanger-based studies (Q1 and allele-specific estimates from beta-binomial mixture models, here updated to e2.m1), and also scores that have been used by other groups in their NGS-based studies that are also reported by D4Z4caster (mean (69) and median (68)). For the samples in the UNR IRB study, the Q1 and e2.m1 scores outperformed the mean and pct10 scores based on AUC and G-mean, though the differences were modest and may be within sampling error. Even a small increase in specificity can result in a profound increase in the positive-predictive value of a test (i.e., the probability that a subject predicted to have FSHD really has FSHD) when the tested non-FSHD subjects vastly outnumber the tested FSHD subjects (as would be the case for broad screening), but firmly establishing such a difference may require both larger sample sizes and more confidence in the ground truth used to evaluate accuracy.

The examples of misclassified samples that are highlighted in the Results section reinforce the notion that the genetic FSHD status may in some cases be ambiguous, as may be the clinical FSHD status, so it should not be surprising that predictions of FSHD status based on methylation levels also sometimes fall into a gray area. The logistic regression models we use attempt to quantify this uncertainty, and some misclassifications of samples in the gray area are to be expected. Less expected are misclassifications of samples outside the gray area, which represent confident but incorrect predictions, although as discussed many of the most striking cases of this are predicted to be mosaic for contractions. Some of these samples are reported as genetically FSHD and some as genetically not-FSHD, and there may not be a consensus on what percent mosaicism should be required for a genetic diagnosis of FSHD, particularly in the absence of clinical symptoms. We flag such samples so that additional information (e.g., from pedigrees, when available) may be used to aid interpretation of the results. In future work we will further refine and validate D4Z4caster’s estimation of percent mosaicism from methylation data.

Although the focus of this paper was on diagnosis of genetic FSHD based on D4Z4 methylation profiles, such profiles have also been demonstrated to have value in predicting disease severity or prognosing disease progression among subjects with genetic FSHD (12, 13, 46, 69, 71). We expect that the allele-specific estimates of methylation provided by D4Z4caster will again be beneficial in these contexts, as reducing the influence of the non-contracted allele on the methylation scores can improve the power to detect disease-associated relationships with methylation levels from the pathogenic contracted allele.

## Author Contributions

TIJ, PLJ, and ODK conceived of the study, developed the technology, and wrote the manuscript. TIJ, BZE, MNF, TG, and PLJ performed experiments. ODK wrote the software code and performed the statistical analysis.

## Institutional Review Board Statement

The study was conducted according to the guidelines of the Declaration of Helsinki, and approved by the University of Nevada, Reno Institutional Review Board (#1316095, approved on 9 October 2018).

## Informed Consent Statement

All qualified subjects provided written informed consent.

## Data Availability Statement

Data for this project will be deposited in Zenodo (10.5281/zenodo.20187613). This includes D4Z4caster reports and other processed data for each sample. It also includes bisulfite sequencing reads (FASTA files for Sanger; BAM files for NGS, down-sampled to a maximum of 10000 reads per reference amplicon).

## Acknowledgements

We dedicate this work to the memory of Mr. Daniel Paul Perez, a fierce advocate for FSHD research, great supporter of our labs, and close personal friend. We thank the FSHD patients and families for participating in our research program to improve and make FSHD diagnostics accessible to everyone around the world. We thank Dr. Rabi Tawil and Dr. Steve Moore for providing cell, saliva, and gDNA samples from their patients and repositories.

## List of Supplemental Figures

1. IRB study: prediction of genetic FSHD status based on Q1 score.
2. IRB study: prediction of genetic FSHD status based on ex.m1 score.
3. IRB study: prediction of genetic FSHD status based on pct10 score.
4. IRB study: prediction of genetic FSHD status based on pct05 score.
5. IRB study: prediction of genetic FSHD status based on pct03 score.
6. IRB study: prediction of genetic FSHD status based on mean score.
7. IRB study: prediction of genetic FSHD status based on median (Q2) score.
8. IRB study: prediction of genetic FSHD status with alternate score for A/L samples.
9. IRB study: calibration of predicted probabilities.
10. IRB study: prediction of genetic FSHD status with milder regularization penalty lambda.
11. Samples flagged for potential clonal artifacts.
12. IRB study: methylation scores vs number of 4A RUs for FSHD1 samples.

## List of Supplemental Tables

1. Sample characteristics.
2. NGS BSS Primers.
3. Sequences of reference amplicons.
4. Methylation summary statistics for each sample.
5. Coefficients and cutoffs for regularized logistic regression models.

## References

1. Deenen JC, Arnts H, van der Maarel SM, Padberg GW, Verschuuren JJ, Bakker E, et al. Population-based incidence and prevalence of facioscapulohumeral dystrophy. Neurology. 2014;83:1056–9.

2. Orphanet. Prevalence and incidence of rare diseases: Bibliographic data: Inserm; 2021 [cited 2021 2021]. Available from: http://www.orpha.net/orphacom/cahiers/docs/GB/Prevalence_of_rare_diseases_by_alphabetical_list.pdf.

3. Lemmers RJ, O’Shea S, Padberg GW, Lunt PW, van der Maarel SM. Best practice guidelines on genetic diagnostics of Facioscapulohumeral muscular dystrophy: workshop 9th June 2010, LUMC, Leiden, The Netherlands. Neuromuscul Disord. 2012;22(5):463–70.

4. Montagnese F, de Valle K, Lemmers R, Mul K, Dumonceaux J, Voermans N, et al. 268th ENMC workshop - Genetic diagnosis, clinical classification, outcome measures, and biomarkers in Facioscapulohumeral Muscular Dystrophy (FSHD): Relevance for clinical trials. Neuromuscul Disord. 2023;33(5):447–62.

5. Himeda CL, Jones PL. The Genetics and Epigenetics of Facioscapulohumeral Muscular Dystrophy. Annu Rev Genomics Hum Genet. 2019;20:265–91.

6. Padberg GW. Facioscapulohumeral Disease [thesis]. Leiden, the Netherlands: Leiden University; 1982.

7. Mul K. Facioscapulohumeral Muscular Dystrophy. Continuum (Minneap Minn). 2022;28(6):1735–51.

8. Zatz M, Marie SK, Cerqueira A, Vainzof M, Pavanello RC, Passos-Bueno MR. The facioscapulohumeral muscular dystrophy (FSHD1) gene affects males more severely and more frequently than females. American journal of medical genetics. 1998;77(2):155–61.

9. Tonini MM, Passos-Bueno MR, Cerqueira A, Matioli SR, Pavanello R, Zatz M. Asymptomatic carriers and gender differences in facioscapulohumeral muscular dystrophy (FSHD). Neuromuscul Disord. 2004;14(1):33–8.

10. Arashiro P, Eisenberg I, Kho AT, Cerqueira AM, Canovas M, Silva HC, et al. Transcriptional regulation differs in affected facioscapulohumeral muscular dystrophy patients compared to asymptomatic related carriers. Proc Natl Acad Sci U S A. 2009;106(15):6220–5.

11. Jones TI, Chen JC, Rahimov F, Homma S, Arashiro P, Beermann ML, et al. Facioscapulohumeral muscular dystrophy family studies of DUX4 expression: evidence for disease modifiers and a quantitative model of pathogenesis. Hum Mol Genet. 2012;21(20):4419–30.

12. Lemmers RJ, Goeman JJ, Van Der Vliet PJ, Van Nieuwenhuizen MP, Balog J, Vos-Versteeg M, et al. Inter-individual differences in CpG methylation at D4Z4 correlate with clinical variability in FSHD1 and FSHD2. Hum Mol Genet. 2015;24(3):659–69.

13. Jones TI, King OD, Himeda CL, Homma S, Chen JC, Beermann ML, et al. Individual epigenetic status of the pathogenic D4Z4 macrosatellite correlates with disease in facioscapulohumeral muscular dystrophy. Clinical epigenetics. 2015;7(1):37.

14. Scionti I, Greco F, Ricci G, Govi M, Arashiro P, Vercelli L, et al. Large-scale population analysis challenges the current criteria for the molecular diagnosis of fascioscapulohumeral muscular dystrophy. Am J Hum Genet. 2012;90(4):628–35.

15. Lemmers RJ, Tawil R, Petek LM, Balog J, Block GJ, Santen GW, et al. Digenic inheritance of an SMCHD1 mutation and an FSHD-permissive D4Z4 allele causes facioscapulohumeral muscular dystrophy type 2. Nat Genet. 2012;44(12):1370–4.

16. van den Boogaard ML, Lemmers RJ, Balog J, Wohlgemuth M, Auranen M, Mitsuhashi S, et al. Mutations in DNMT3B Modify Epigenetic Repression of the D4Z4 Repeat and the Penetrance of Facioscapulohumeral Dystrophy. Am J Hum Genet. 2016;98(5):1020–9.

17. Flanigan KM, Coffeen CM, Sexton L, Stauffer D, Brunner S, Leppert MF. Genetic characterization of a large, historically significant Utah kindred with facioscapulohumeral dystrophy. Neuromuscul Disord. 2001;11(6-7):525–9.

18. Jones TI, Himeda CL, Perez DP, Jones PL. Large family cohorts of lymphoblastoid cells provide a new cellular model for investigating facioscapulohumeral muscular dystrophy. Neuromuscul Disord. 2017;27(3):221–38.

19. Gabriels J, Beckers MC, Ding H, De Vriese A, Plaisance S, van der Maarel SM, et al. Nucleotide sequence of the partially deleted D4Z4 locus in a patient with FSHD identifies a putative gene within each 3.3 kb element. Gene. 1999;236(1):25–32.

20. Kowaljow V, Marcowycz A, Ansseau E, Conde CB, Sauvage S, Matteotti C, et al. The DUX4 gene at the FSHD1A locus encodes a pro-apoptotic protein. Neuromuscul Disord. 2007;17(8):611–23.

21. Lemmers RJ, van der Vliet PJ, Klooster R, Sacconi S, Camano P, Dauwerse JG, et al. A unifying genetic model for facioscapulohumeral muscular dystrophy. Science. 2010;329(5999):1650–3.

22. Snider L, Geng LN, Lemmers RJ, Kyba M, Ware CB, Nelson AM, et al. Facioscapulohumeral dystrophy: incomplete suppression of a retrotransposed gene. PLoS Genet. 2010;6(10):e1001181.

23. Wijmenga C, Hewitt JE, Sandkuijl LA, Clark LN, Wright TJ, Dauwerse HG, et al. Chromosome 4q DNA rearrangements associated with facioscapulohumeral muscular dystrophy. Nat Genet. 1992;2(1):26–30.

24. van Deutekom JC, Wijmenga C, van Tienhoven EA, Gruter AM, Hewitt JE, Padberg GW, et al. FSHD associated DNA rearrangements are due to deletions of integral copies of a 3.2 kb tandemly repeated unit. Hum Mol Genet. 1993;2(12):2037–42.

25. van Overveld PG, Lemmers RJ, Sandkuijl LA, Enthoven L, Winokur ST, Bakels F, et al. Hypomethylation of D4Z4 in 4q-linked and non-4q-linked facioscapulohumeral muscular dystrophy. Nat Genet. 2003;35(4):315–7.

26. van Overveld PG, Enthoven L, Ricci E, Rossi M, Felicetti L, Jeanpierre M, et al. Variable hypomethylation of D4Z4 in facioscapulohumeral muscular dystrophy. Annals of neurology. 2005;58(4):569–76.

27. Hamanaka K, Sikrova D, Mitsuhashi S, Masuda H, Sekiguchi Y, Sugiyama A, et al. Homozygous nonsense variant in LRIF1 associated with facioscapulohumeral muscular dystrophy. Neurology. 2020;94(23):e2441–e7.

28. Giardina E, Camano P, Burton-Jones S, Ravenscroft G, Henning F, Magdinier F, et al. Best practice guidelines on genetic diagnostics of facioscapulohumeral muscular dystrophy: Update of the 2012 guidelines. Clin Genet. 2024;106(1):13–26.

29. Lemmers RJ, de Kievit P, Sandkuijl L, Padberg GW, van Ommen GJ, Frants RR, et al. Facioscapulohumeral muscular dystrophy is uniquely associated with one of the two variants of the 4q subtelomere. Nat Genet. 2002;32(2):235–6.

30. Lemmers RJ, Wohlgemuth M, van der Gaag KJ, van der Vliet PJ, van Teijlingen CM, de Knijff P, et al. Specific sequence variations within the 4q35 region are associated with facioscapulohumeral muscular dystrophy. Am J Hum Genet. 2007;81(5):884–94.

31. Lemmers R, van der Vliet PJ, Blatnik A, Balog J, Zidar J, Henderson D, et al. Chromosome 10q-linked FSHD identifies DUX4 as principal disease gene. J Med Genet. 2022;59(2):180–8.

32. Lemmers RJ, Wohlgemuth M, Frants RR, Padberg GW, Morava E, van der Maarel SM. Contractions of D4Z4 on 4qB subtelomeres do not cause facioscapulohumeral muscular dystrophy. Am J Hum Genet. 2004;75(6):1124–30.

33. Lemmers RJ, van der Vliet PJ, van der Gaag KJ, Zuniga S, Frants RR, de Knijff P, et al. Worldwide population analysis of the 4q and 10q subtelomeres identifies only four discrete interchromosomal sequence transfers in human evolution. Am J Hum Genet. 2010;86(3):364–77.

34. Lemmers RJ. Analyzing Copy Number Variation Using Pulsed-Field Gel Electrophoresis: Providing a Genetic Diagnosis for FSHD1. Methods Mol Biol. 2017;1492:107–25.

35. Vasale J, Boyar F, Jocson M, Sulcova V, Chan P, Liaquat K, et al. Molecular combing compared to Southern blot for measuring D4Z4 contractions in FSHD. Neuromuscul Disord. 2015;25(12):945–51.

36. Mitsuhashi S, Nakagawa S, Takahashi Ueda M, Imanishi T, Frith MC, Mitsuhashi H. Nanopore-based single molecule sequencing of the D4Z4 array responsible for facioscapulohumeral muscular dystrophy. Sci Rep. 2017;7(1):14789.

37. Dai Y, Li P, Wang Z, Liang F, Yang F, Fang L, et al. Single-molecule optical mapping enables quantitative measurement of D4Z4 repeats in facioscapulohumeral muscular dystrophy (FSHD). J Med Genet. 2020;57(2):109–20.

38. Hiramuki Y, Kure Y, Saito Y, Ogawa M, Ishikawa K, Mori-Yoshimura M, et al. Simultaneous measurement of the size and methylation of chromosome 4qA-D4Z4 repeats in facioscapulohumeral muscular dystrophy by long-read sequencing. J Transl Med. 2022;20(1):517.

39. Butterfield RJ, Dunn DM, Duvall B, Moldt S, Weiss RB. Deciphering D4Z4 CpG methylation gradients in fascioscapulohumeral muscular dystrophy using nanopore sequencing. bioRxiv. 2023.

40. Hartweck LM, Anderson LJ, Lemmers RJ, Dandapat A, Toso EA, Dalton JC, et al. A focal domain of extreme demethylation within D4Z4 in FSHD2. Neurology. 2013;80(4):392–9.

41. Jones TI, Yan C, Sapp PC, McKenna-Yasek D, Kang PB, Quinn C, et al. Identifying diagnostic DNA methylation profiles for facioscapulohumeral muscular dystrophy in blood and saliva using bisulfite sequencing. Clinical epigenetics. 2014;6(1):23.

42. Mitsuhashi S, Boyden SE, Estrella EA, Jones TI, Rahimov F, Yu TW, et al. Exome sequencing identifies a novel SMCHD1 mutation in facioscapulohumeral muscular dystrophy 2. Neuromuscul Disord. 2013;23:975–80.

43. Landrum MJ, Lee JM, Riley GR, Jang W, Rubinstein WS, Church DM, et al. ClinVar: public archive of relationships among sequence variation and human phenotype. Nucleic Acids Res. 2014;42(Database issue):D980–5.

44. Gerard L, Delourme M, Tardy C, Ganne B, Perrin P, Chaix C, et al. SMCHD1 genetic variants in type 2 facioscapulohumeral dystrophy and challenges in predicting pathogenicity and disease penetrance. Eur J Hum Genet. 2025;33(6):784–92.

45. Gould T, Jones TI, Jones PL. Precise Epigenetic Analysis Using Targeted Bisulfite Genomic Sequencing Distinguishes FSHD1, FSHD2, and Healthy Subjects. Diagnostics. 2021;11:1469.

46. Calandra P, Cascino I, Lemmers RJ, Galluzzi G, Teveroni E, Monforte M, et al. Allele-specific DNA hypomethylation characterises FSHD1 and FSHD2. J Med Genet. 2016;53(5):348–55.

47. Krueger F, Andrews SR. Bismark: a flexible aligner and methylation caller for Bisulfite-Seq applications. Bioinformatics. 2011;27(11):1571–2.

48. Lemmers R, Butterfield R, van der Vliet PJ, de Bleecker JL, van der Pol L, Dunn DM, et al. Autosomal dominant in cis D4Z4 repeat array duplication alleles in facioscapulohumeral dystrophy. Brain. 2024;147(2):414–26.

49. van der Maarel SM, Deidda G, Lemmers RJ, van Overveld PG, van der Wielen M, Hewitt JE, et al. De novo facioscapulohumeral muscular dystrophy: frequent somatic mosaicism, sex-dependent phenotype, and the role of mitotic transchromosomal repeat interaction between chromosomes 4 and 10. Am J Hum Genet. 2000;66(1):26–35.

50. Lemmers RJ, van der Wielen MJ, Bakker E, Padberg GW, Frants RR, van der Maarel SM. Somatic mosaicism in FSHD often goes undetected. Annals of neurology. 2004;55(6):845–50.

51. Hoeting JA, Madigan D, Raftery AE, Volinsky CT. Bayesian Model Averaging: A Tutorial. Statistical Science. 1999;14(4):382–401.

52. Molder F, Jablonski KP, Letcher B, Hall MB, van Dyken PC, Tomkins-Tinch CH, et al. Sustainable data analysis with Snakemake. F1000Res. 2021;10:33.

53. Langmead B, Salzberg SL. Fast gapped-read alignment with Bowtie 2. Nat Methods. 2012;9(4):357–9.

54. Andrews S. FastQC: A Quality Control Tool for High Throughput Sequence Data 2010.

55. Li H, Handsaker B, Wysoker A, Fennell T, Ruan J, Homer N, et al. The Sequence Alignment/Map format and SAMtools. Bioinformatics. 2009;25(16):2078–9.

56. Merkel D. Docker: lightweight linux containers for consistent development and deployment. Linux Journal. 2014;239(2).

57. Kircher M, Sawyer S, Meyer M. Double indexing overcomes inaccuracies in multiplex sequencing on the Illumina platform. Nucleic Acids Res. 2012;40(1):e3.

58. Seitz V, Schaper S, Droge A, Lenze D, Hummel M, Hennig S. A new method to prevent carry-over contaminations in two-step PCR NGS library preparations. Nucleic Acids Res. 2015;43(20):e135.

59. Kubat M, Matwin S. Addressing the Curse of Imbalanced Training Sets: One-Sided Selection. Proceedings of the 14th International Conference on Machine Learning. 1997:179–86.

60. Robin X, Turck N, Hainard A, Tiberti N, Lisacek F, Sanchez JC, et al. pROC: an open-source package for R and S+ to analyze and compare ROC curves. BMC Bioinformatics. 2011;12:77.

61. Prentice RL, Pyke R. Logistic disease incidence models and case-control studies. Biometrika. 1979;66(3):403–11.

62. Rose S, van der Laan MJ. Simple optimal weighting of cases and controls in case-control studies. Int J Biostat. 2008;4(1):Article 19.

63. Friedman J, Hastie T, Tibshirani R. Regularization Paths for Generalized Linear Models via Coordinate Descent. J Stat Softw. 2010;33(1):1–22.

64. Kuhn M, Wickham H. Tidymodels: a collection of packages for modeling and machine learning using tidyverse principles. 2020.

65. Van Calster B, Nieboer D, Vergouwe Y, De Cock B, Pencina MJ, Steyerberg EW. A calibration hierarchy for risk models was defined: from utopia to empirical data. J Clin Epidemiol. 2016;74:167–76.

66. Strafella C, Megalizzi D, Trastulli G, Proietti Piorgo E, Colantoni L, Tasca G, et al. Integrating D4Z4 methylation analysis into clinical practice: improvement of FSHD molecular diagnosis through distinct thresholds for 4qA/4qA and 4qA/4qB patients. Clinical epigenetics. 2024;16(1):148.

67. Wong CJ, Friedman SD, Snider L, Bennett SR, Jones TI, Jones PL, et al. Regional and bilateral MRI and gene signatures in facioscapulohumeral dystrophy: implications for clinical trial design and mechanisms of disease progression. Hum Mol Genet. 2024;33(8):698–708.

68. Xia X, Cheng N, Liu Y, Yue D, Gao M, Hu C, et al. 4qA D4Z4 Methylation Test as a Valuable Complement for Differential Diagnosis in Patients with a Facioscapulohumeral Muscular Dystrophy-Like Phenotype. J Mol Diagn. 2025;27(5):405–18.

69. Erdmann H, Scharf F, Gehling S, Benet-Pages A, Jakubiczka S, Becker K, et al. Methylation of the 4q35 D4Z4 repeat defines disease status in facioscapulohumeral muscular dystrophy. Brain. 2023;146(4):1388–402.

70. Lemmers RJ, van der Vliet PJ, Balog J, Goeman JJ, Arindrarto W, Krom YD, et al. Deep characterization of a common D4Z4 variant identifies biallelic DUX4 expression as a modifier for disease penetrance in FSHD2. Eur J Hum Genet. 2018;26(1):94–106.

71. Zheng F, Qiu L, Chen L, Zheng Y, Lin X, He J, et al. Association of 4qA-Specific Distal D4Z4 Hypomethylation With Disease Severity and Progression in Facioscapulohumeral Muscular Dystrophy. Neurology. 2023.

72. Chen S, Francioli LC, Goodrich JK, Collins RL, Kanai M, Wang Q, et al. A genomic mutational constraint map using variation in 76,156 human genomes. Nature. 2024;625(7993):92–100.

73. Puma A, Tammam G, Ezaru A, Slioui A, Torchia E, Tasca G, et al. Double trouble: a comprehensive study into unrelated genetic comorbidities in adult patients with Facioscapulohumeral Muscular Dystrophy Type I. Eur J Hum Genet. 2025;33(8):1006–14.

74. de Greef JC, Lemmers RJ, van Engelen BG, Sacconi S, Venance SL, Frants RR, et al. Common epigenetic changes of D4Z4 in contraction-dependent and contraction-independent FSHD. Hum Mutat. 2009;30(10):1449–59.

75. Sacconi S, Briand-Suleau A, Gros M, Baudoin C, Lemmers R, Rondeau S, et al. FSHD1 and FSHD2 form a disease continuum. Neurology. 2019;92(19):e2273–e85.

76. Xiao LC, Semwal A, St John B, Zeglinski K, Su S, Lancaster J, et al. D4Z4End2End: complete genetic and epigenetic architecture of D4Z4 macrosatellites in FSHD, BAMS and reference cohorts. medRxiv. 2025:2025.04.24.25326320.

77. Rohde C, Zhang Y, Reinhardt R, Jeltsch A. BISMA--fast and accurate bisulfite sequencing data analysis of individual clones from unique and repetitive sequences. BMC Bioinformatics. 2010;11:230.

78. Gordon CT, Xue S, Yigit G, Filali H, Chen K, Rosin N, et al. De novo mutations in SMCHD1 cause Bosma arhinia microphthalmia syndrome and abrogate nasal development. Nat Genet. 2017;49(2):249–55.

79. Shaw ND, Brand H, Kupchinsky ZA, Bengani H, Plummer L, Jones TI, et al. SMCHD1 mutations associated with a rare muscular dystrophy can also cause isolated arhinia and Bosma arhinia microphthalmia syndrome. Nat Genet. 2017;49(2):238–48.

80. Lemmers R, van der Stoep N, Vliet PJV, Moore SA, San Leon Granado D, Johnson K, et al. SMCHD1 mutation spectrum for facioscapulohumeral muscular dystrophy type 2 (FSHD2) and Bosma arhinia microphthalmia syndrome (BAMS) reveals disease-specific localisation of variants in the ATPase domain. J Med Genet. 2019;56(10):693–700.

81. Zatz M, Marie SK, Passos-Bueno MR, Vainzof M, Campiotto S, Cerqueira A, et al. High proportion of new mutations and possible anticipation in Brazilian facioscapulohumeral muscular dystrophy families. Am J Hum Genet. 1995;56(1):99–105.

82. Lunt PW. 44th ENMC International Workshop: Facioscapulohumeral Muscular Dystrophy: Molecular Studies 19-21 July 1996, Naarden, The Netherlands. Neuromuscul Disord. 1998;8(2):126–30.

83. Nguyen K, Walrafen P, Bernard R, Attarian S, Chaix C, Vovan C, et al. Molecular combing reveals allelic combinations in facioscapulohumeral dystrophy. Annals of neurology. 2011;70(4):627–33.

84. Caputo V, Megalizzi D, Fabrizio C, Termine A, Colantoni L, Bax C, et al. D4Z4 Methylation Levels Combined with a Machine Learning Pipeline Highlight Single CpG Sites as Discriminating Biomarkers for FSHD Patients. Cells. 2022;11(24).

85. Zheng F, Qiu L, Chen L, Zheng Y, He Q, Lin X, et al. An epigenetic basis for genetic anticipation in facioscapulohumeral muscular dystrophy type 1. Brain. 2023.

86. Erdmann H, Scharf F, Hallermayr A, Barsegehyan H, Walter MC, Holinski-Feder E, et al. Reply: An epigenetic basis for genetic anticipation in facioscapulohumeral muscular dystrophy type 1. Brain. 2023.

