## Supplemental Figures for "The D4Z4caster DNA methylation signature identifies individuals at epigenetic risk for developing facioscapulohumeral muscular dystrophy (FSHD)"

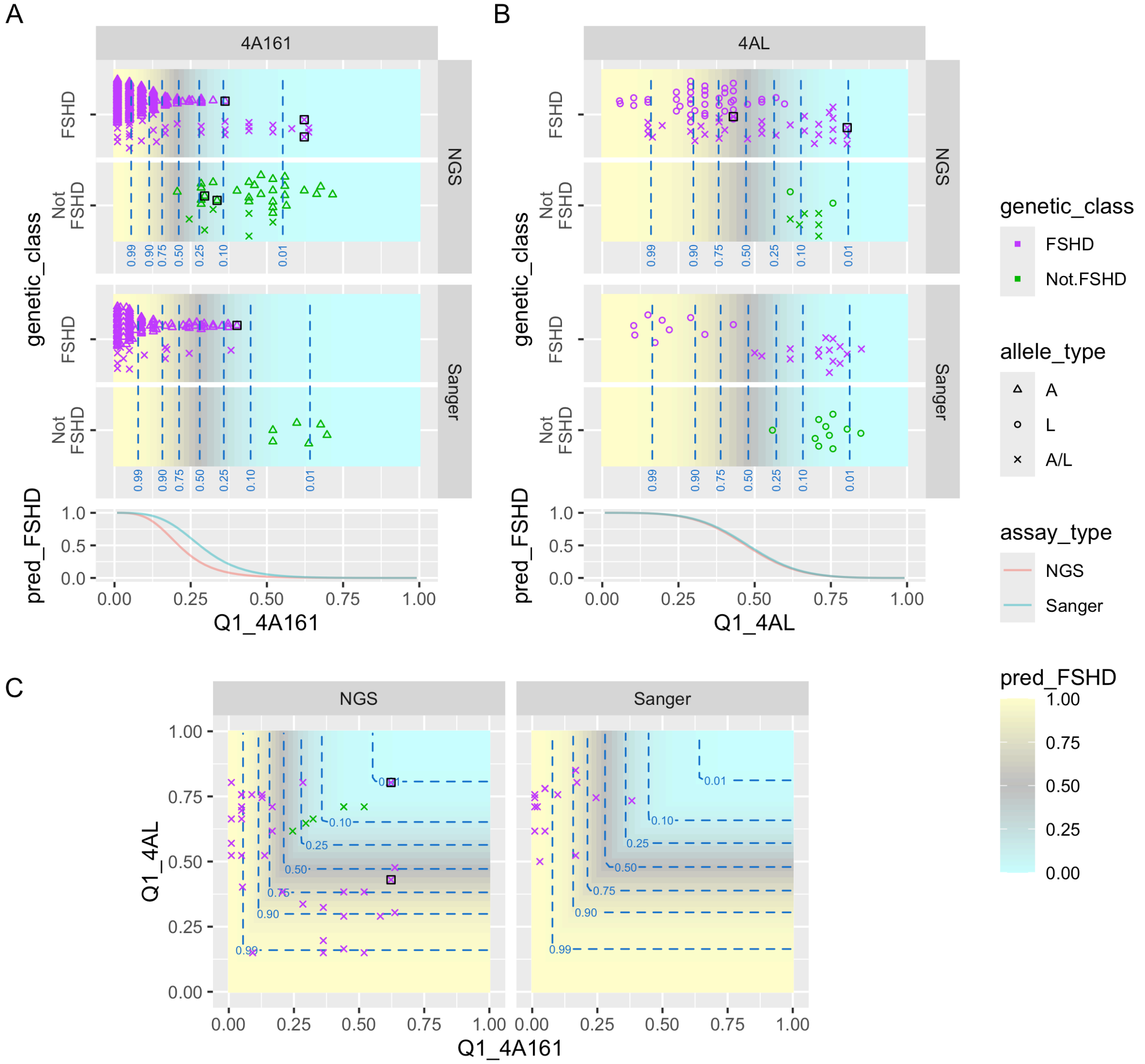


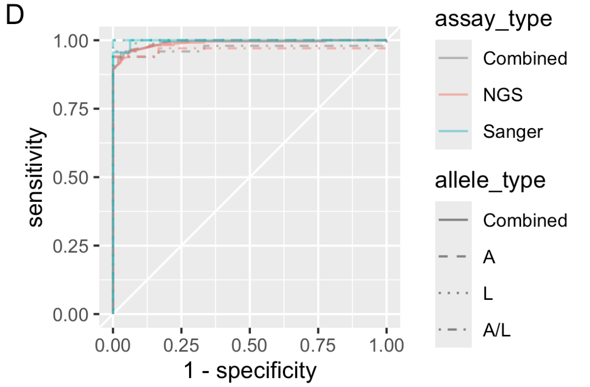


E

| assay_type | allele_type | n_pos | n_neg | AUC | AUC_CI | max_Gmean |
|---|---|---|---|---|---|---|
| Combined | Combined | 343 | 52 | 0.991 | 0.984-0.998 | 0.957 |
| NGS | Combined | 230 | 36 | 0.988 | 0.978-0.998 | 0.951 |
| Sanger | Combined | 113 | 16 | 0.997 | 0.991-1.000 | 0.978 |
| Combined | A | 245 | 34 | 0.993 | 0.986-1.000 | 0.965 |
| NGS | A | 156 | 28 | 0.992 | 0.983-1.000 | 0.960 |
| Sanger | A | 89 | 6 | 1.000 | 1.000-1.000 | 1.000 |
| Combined | L | 49 | 12 | 0.997 | 0.988-1.000 | 0.979 |
| NGS | L | 40 | 2 | 1.000 | 1.000-1.000 | 1.000 |
| Sanger | L | 9 | 10 | 1.000 | 1.000-1.000 | 1.000 |
| Combined | A/L | 49 | 6 | 0.969 | 0.925-1.000 | 0.969 |
| NGS | A/L | 34 | 6 | 0.966 | 0.907-1.000 | 0.970 |
| Sanger | A/L | 15 | 0 | NA | NA | NA |

**Figure S1:** Q1 methylation scores for 4A161 and 4AL assays in UNR IRB samples. Caption is as for Figure 10 but with Q1 score rather than e2.m1 score.

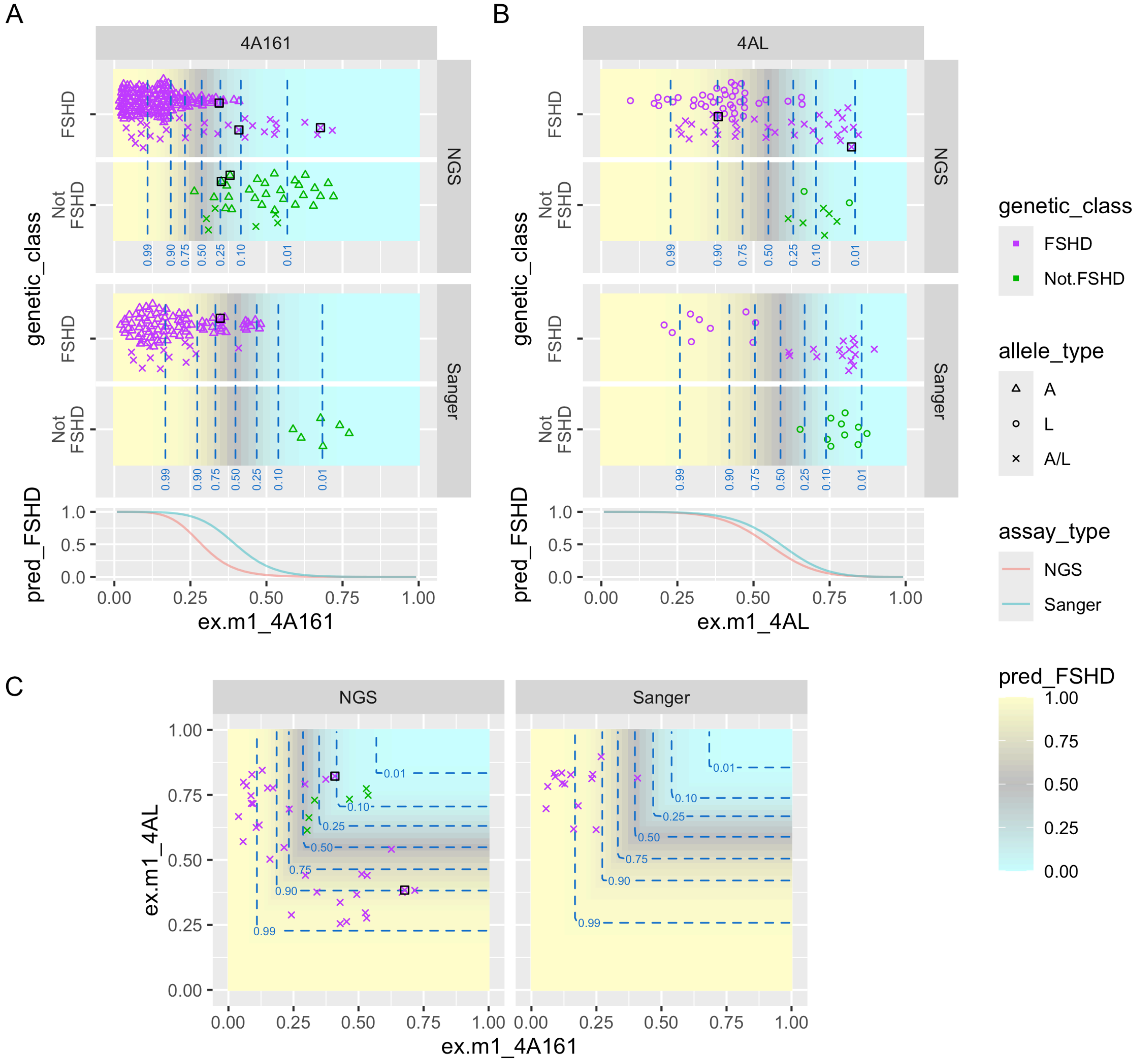


D

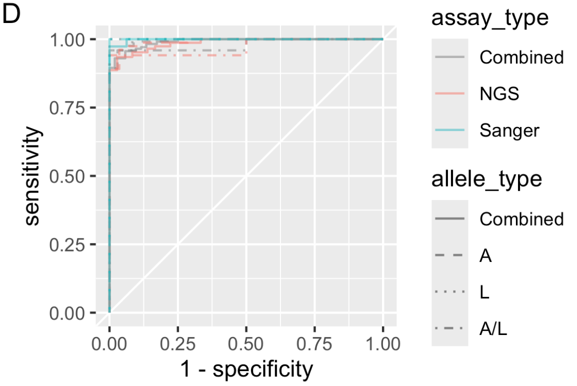


E

| assay_type | allele_type | n_pos | n_neg | AUC | AUC_CI | max_Gmean |
|---|---|---|---|---|---|---|
| Combined | Combined | 343 | 52 | 0.991 | 0.984-0.998 | 0.955 |
| NGS | Combined | 230 | 36 | 0.987 | 0.976-0.997 | 0.953 |
| Sanger | Combined | 113 | 16 | 0.998 | 0.995-1.000 | 0.987 |
| Combined | A | 245 | 34 | 0.994 | 0.986-1.000 | 0.965 |
| NGS | A | 156 | 28 | 0.992 | 0.982-1.000 | 0.963 |
| Sanger | A | 89 | 6 | 1.000 | 1.000-1.000 | 1.000 |
| Combined | L | 49 | 12 | 0.997 | 0.988-1.000 | 0.979 |
| NGS | L | 40 | 2 | 1.000 | 1.000-1.000 | 1.000 |
| Sanger | L | 9 | 10 | 1.000 | 1.000-1.000 | 1.000 |
| Combined | A/L | 49 | 6 | 0.980 | 0.946-1.000 | 0.979 |
| NGS | A/L | 34 | 6 | 0.971 | 0.923-1.000 | 0.970 |
| Sanger | A/L | 15 | 0 | NA | NA | NA |

**Figure S2:** ex.m1 methylation scores for 4A161 and 4AL assays in UNR IRB samples. Caption is as for Figure 10 but with ex.m1 score rather than e2.m1 score.

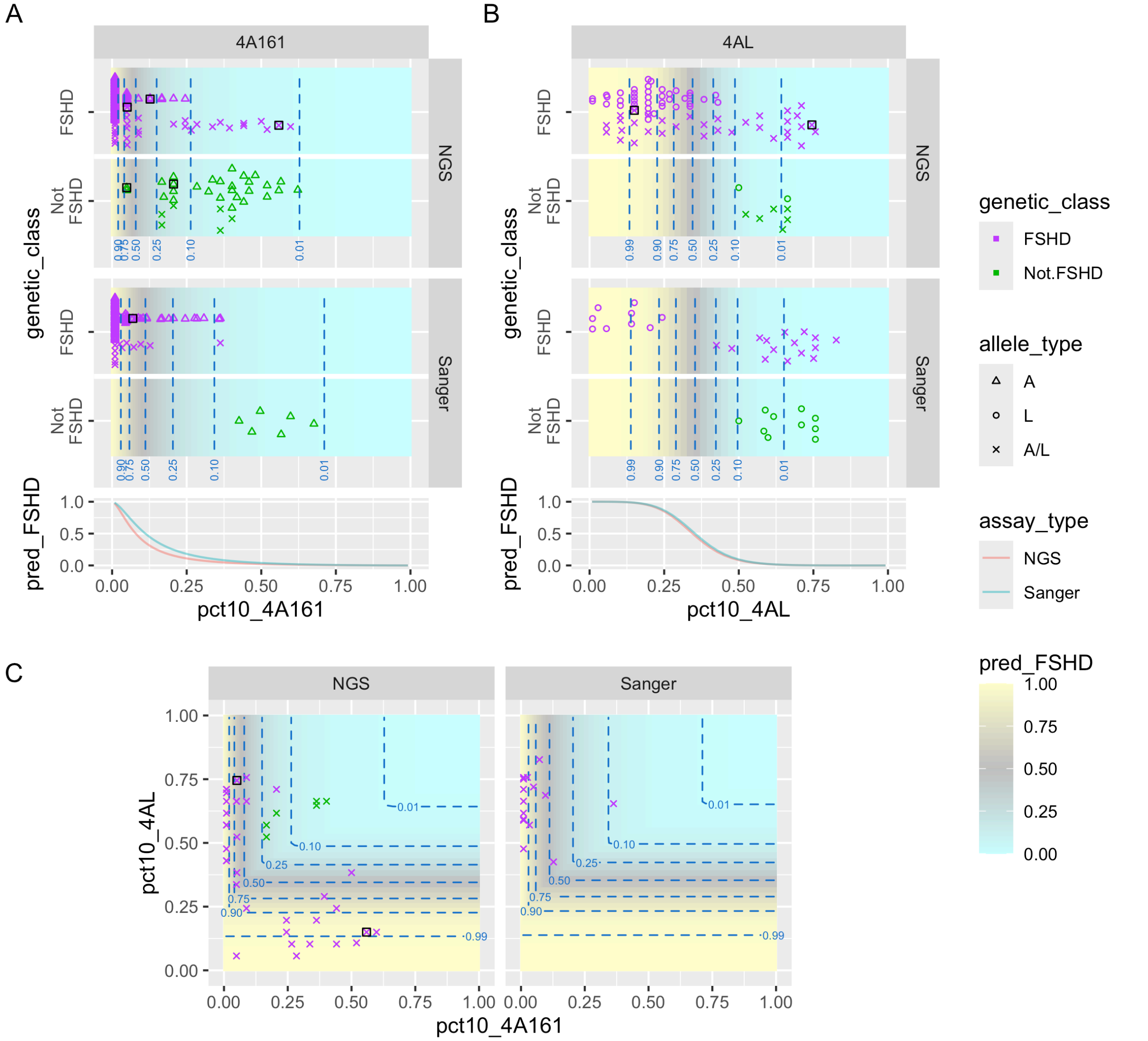


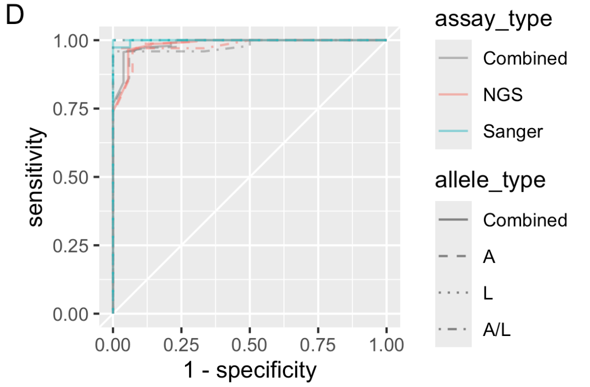


E

| assay_type | allele_type | n_pos | n_neg | AUC | AUC_CI | max_Gmean |
|---|---|---|---|---|---|---|
| Combined | Combined | 343 | 52 | 0.987 | 0.976-0.998 | 0.959 |
| NGS | Combined | 230 | 36 | 0.985 | 0.969-1.000 | 0.955 |
| Sanger | Combined | 113 | 16 | 0.998 | 0.995-1.000 | 0.987 |
| Combined | A | 245 | 34 | 0.984 | 0.968-1.000 | 0.946 |
| NGS | A | 156 | 28 | 0.984 | 0.964-1.000 | 0.945 |
| Sanger | A | 89 | 6 | 1.000 | 1.000-1.000 | 1.000 |
| Combined | L | 49 | 12 | 1.000 | 1.000-1.000 | 1.000 |
| NGS | L | 40 | 2 | 1.000 | 1.000-1.000 | 1.000 |
| Sanger | L | 9 | 10 | 1.000 | 1.000-1.000 | 1.000 |
| Combined | A/L | 49 | 6 | 0.981 | 0.951-1.000 | 0.979 |
| NGS | A/L | 34 | 6 | 0.988 | 0.961-1.000 | 0.985 |
| Sanger | A/L | 15 | 0 | NA | NA | NA |

**Figure S3:** pct10 methylation scores for 4A161 and 4AL assays in UNR IRB samples. Caption is as for Figure 10 but with pct10 score rather than Q1 score.

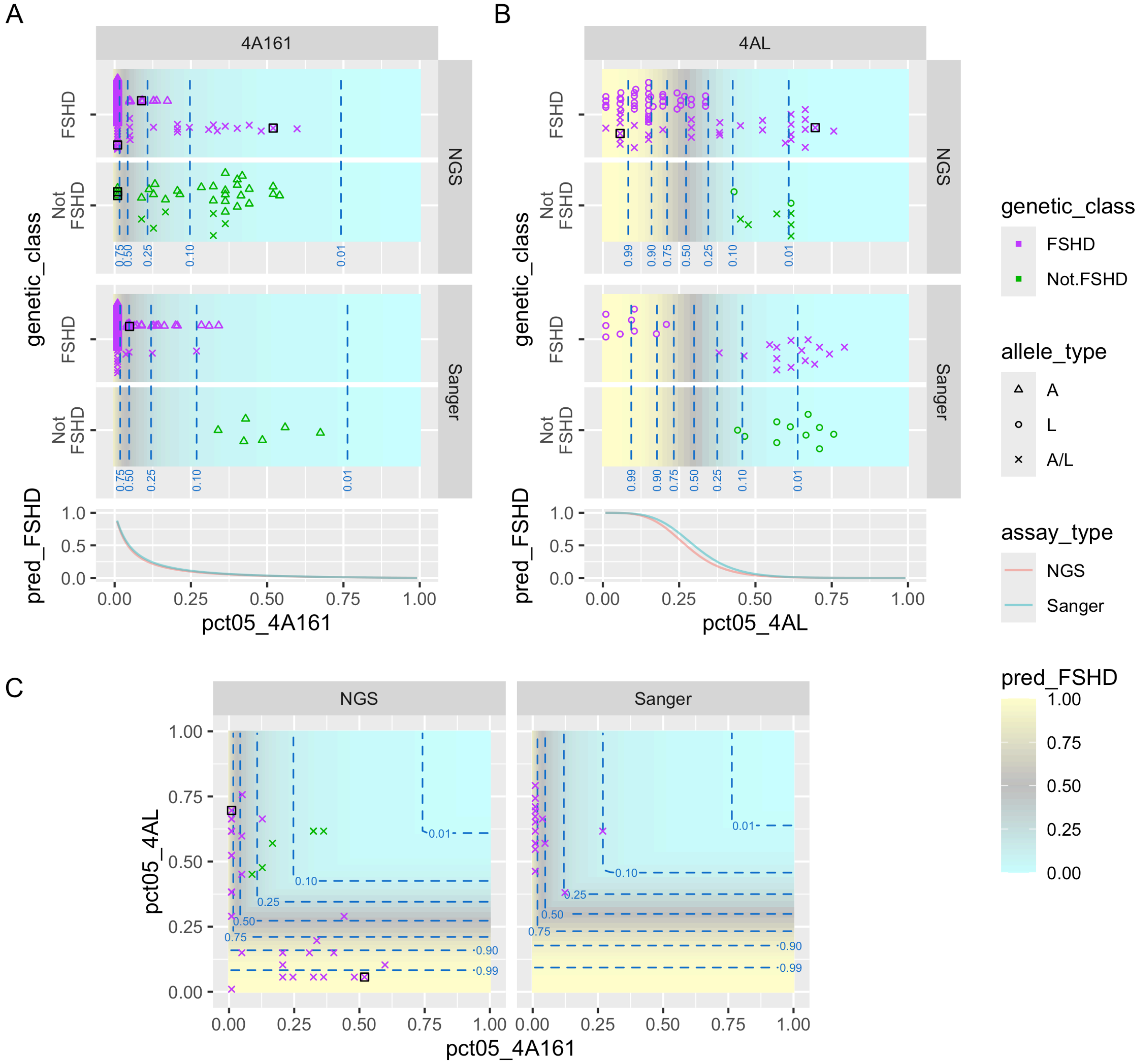


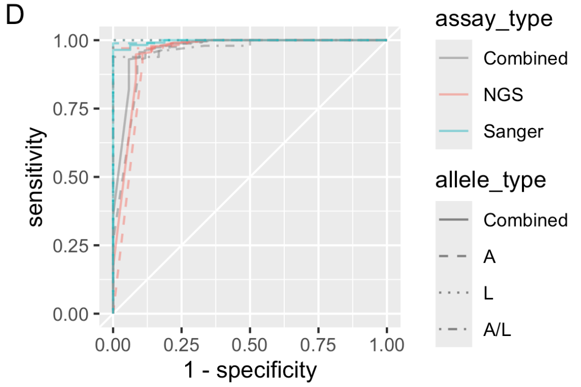


E

| assay_type | allele_type | n_pos | n_neg | AUC | AUC_CI | max_Gmean |
|---|---|---|---|---|---|---|
| Combined | Combined | 343 | 52 | 0.969 | 0.943-0.996 | 0.936 |
| NGS | Combined | 230 | 36 | 0.955 | 0.910-0.999 | 0.932 |
| Sanger | Combined | 113 | 16 | 0.996 | 0.990-1.000 | 0.982 |
| Combined | A | 245 | 34 | 0.953 | 0.910-0.996 | 0.919 |
| NGS | A | 156 | 28 | 0.938 | 0.875-1.000 | 0.920 |
| Sanger | A | 89 | 6 | 0.998 | 0.993-1.000 | 0.994 |
| Combined | L | 49 | 12 | 1.000 | 1.000-1.000 | 1.000 |
| NGS | L | 40 | 2 | 1.000 | 1.000-1.000 | 1.000 |
| Sanger | L | 9 | 10 | 1.000 | 1.000-1.000 | 1.000 |
| Combined | A/L | 49 | 6 | 0.981 | 0.951-1.000 | 0.969 |
| NGS | A/L | 34 | 6 | 0.993 | 0.975-1.000 | 0.985 |
| Sanger | A/L | 15 | 0 | NA | NA | NA |

**Figure S4:** pct05 methylation scores for 4A161 and 4AL assays in UNR IRB samples. Caption is as for Figure 10 but with pct05 score rather than e2.m1 score.

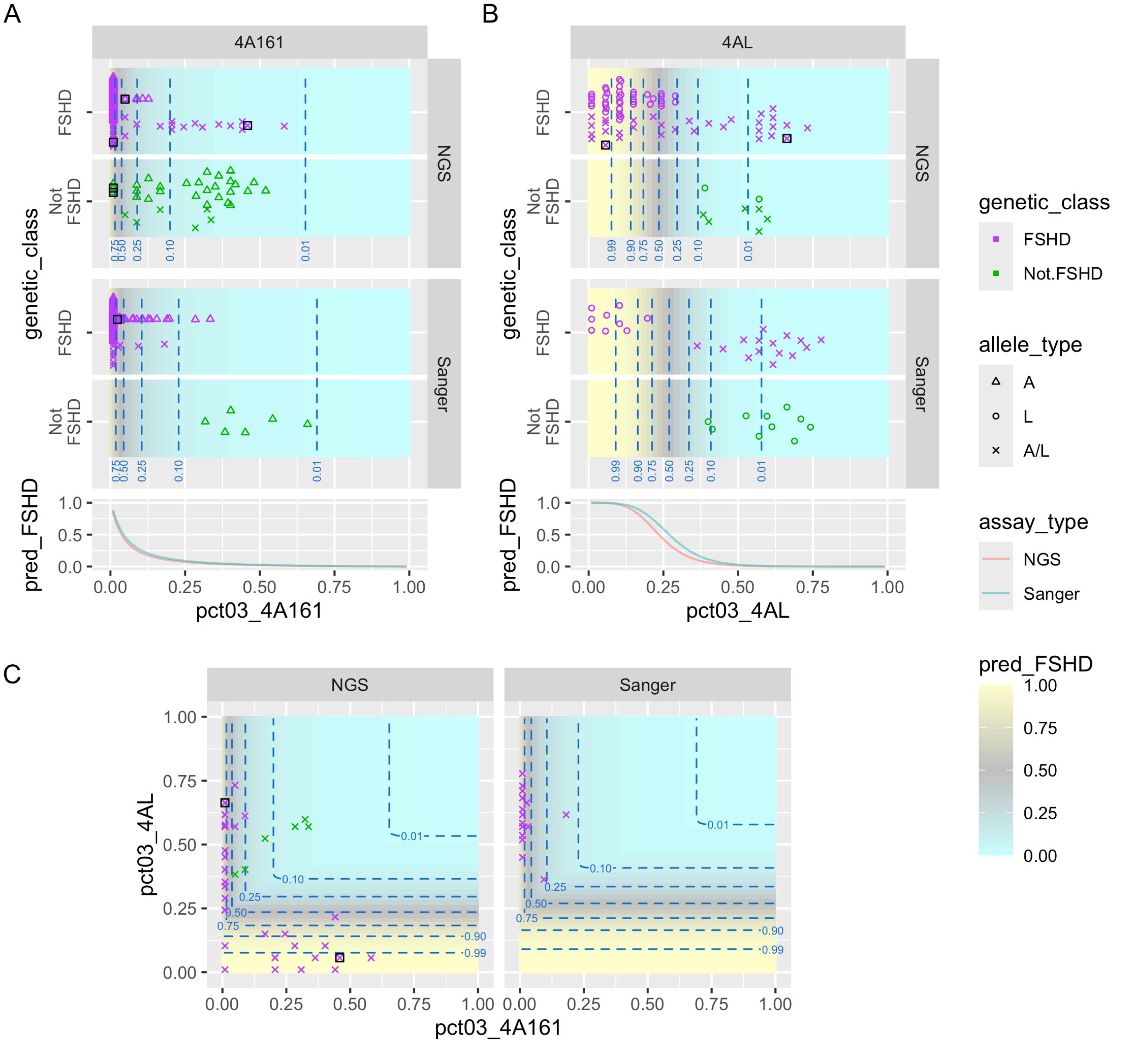


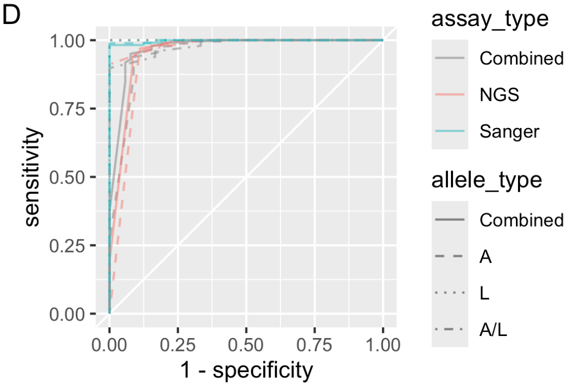


E

| assay_type | allele_type | n_pos | n_neg | AUC | AUC_CI | max_Gmean |
|---|---|---|---|---|---|---|
| Combined | Combined | 343 | 52 | 0.971 | 0.946-0.996 | 0.937 |
| NGS | Combined | 230 | 36 | 0.958 | 0.917-1.000 | 0.928 |
| Sanger | Combined | 113 | 16 | 0.997 | 0.992-1.000 | 0.991 |
| Combined | A | 245 | 34 | 0.954 | 0.912-0.997 | 0.925 |
| NGS | A | 156 | 28 | 0.940 | 0.879-1.000 | 0.927 |
| Sanger | A | 89 | 6 | 0.998 | 0.993-1.000 | 0.994 |
| Combined | L | 49 | 12 | 1.000 | 1.000-1.000 | 1.000 |
| NGS | L | 40 | 2 | 1.000 | 1.000-1.000 | 1.000 |
| Sanger | L | 9 | 10 | 1.000 | 1.000-1.000 | 1.000 |
| Combined | A/L | 49 | 6 | 0.981 | 0.949-1.000 | 0.948 |
| NGS | A/L | 34 | 6 | 0.988 | 0.963-1.000 | 0.955 |
| Sanger | A/L | 15 | 0 | NA | NA | NA |

**Figure S5:** pct03 methylation scores for 4A161 and 4AL assays in UNR IRB samples. Caption is as for Figure 10 but with pct03 score rather than e2.m1 score.

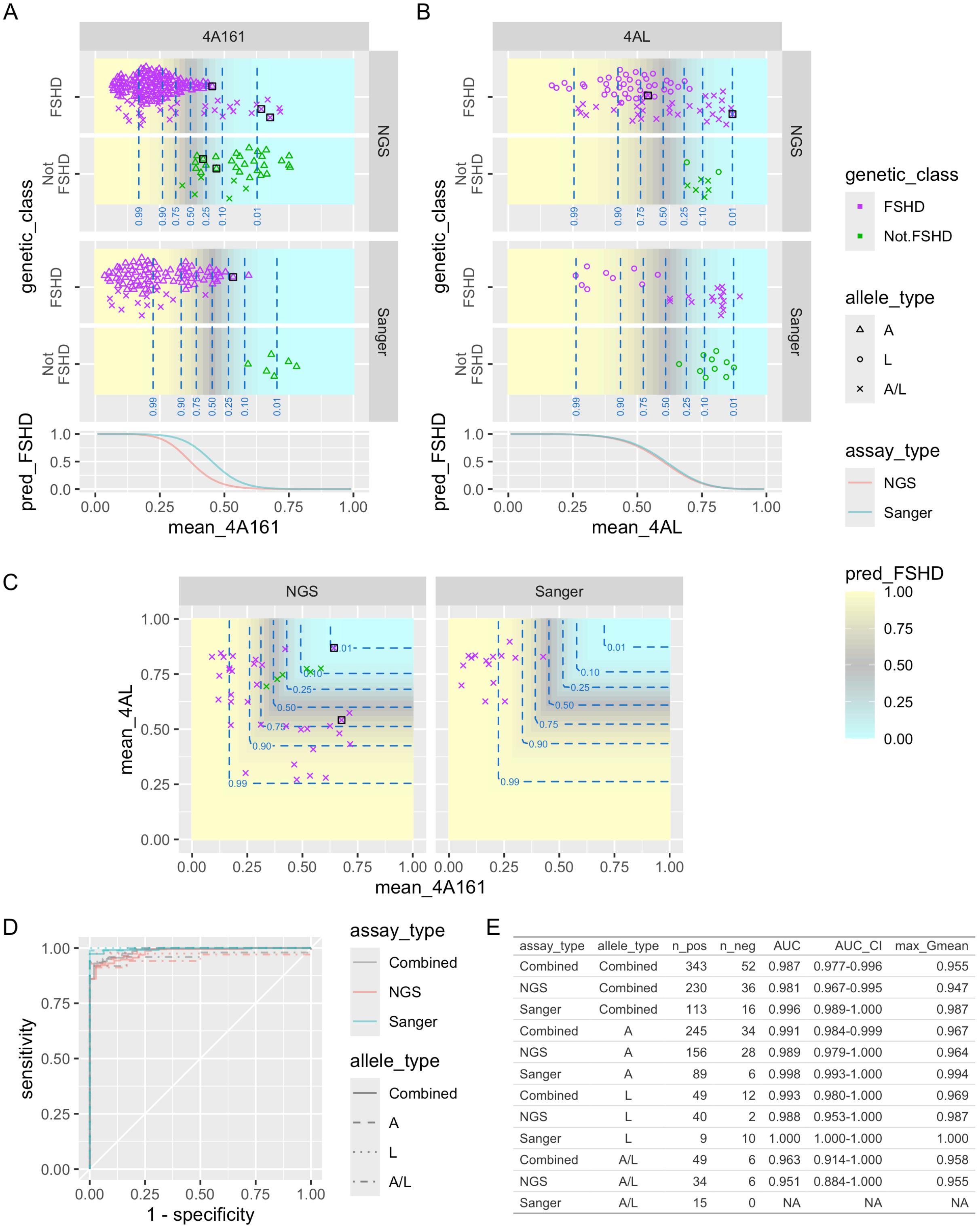


| assay_type | allele_type | n_pos | n_neg | AUC | AUC_CI | max_Gmean |
|---|---|---|---|---|---|---|
| Combined | Combined | 343 | 52 | 0.987 | 0.977-0.996 | 0.955 |
| NGS | Combined | 230 | 36 | 0.981 | 0.967-0.995 | 0.947 |
| Sanger | Combined | 113 | 16 | 0.996 | 0.989-1.000 | 0.987 |
| Combined | A | 245 | 34 | 0.991 | 0.984-0.999 | 0.967 |
| NGS | A | 156 | 28 | 0.989 | 0.979-1.000 | 0.964 |
| Sanger | A | 89 | 6 | 0.998 | 0.993-1.000 | 0.994 |
| Combined | L | 49 | 12 | 0.993 | 0.980-1.000 | 0.969 |
| NGS | L | 40 | 2 | 0.988 | 0.953-1.000 | 0.987 |
| Sanger | L | 9 | 10 | 1.000 | 1.000-1.000 | 1.000 |
| Combined | A/L | 49 | 6 | 0.963 | 0.914-1.000 | 0.958 |
| NGS | A/L | 34 | 6 | 0.951 | 0.884-1.000 | 0.955 |
| Sanger | A/L | 15 | 0 | NA | NA | NA |

**Figure S6:** mean methylation scores for 4A161 and 4AL assays in UNR IRB samples. Caption is as for Figure 10 but with mean score rather than e2.m1 score.

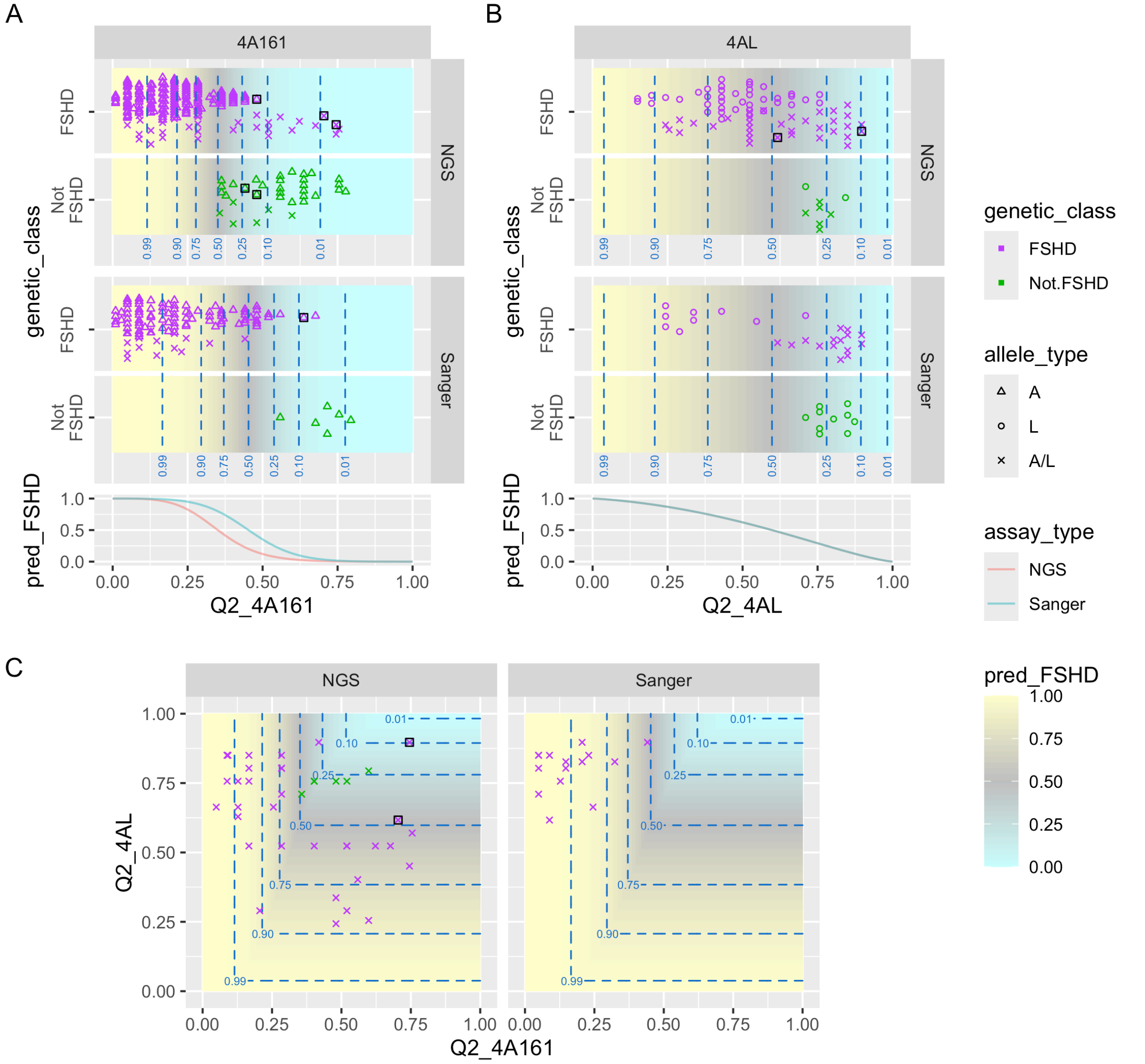


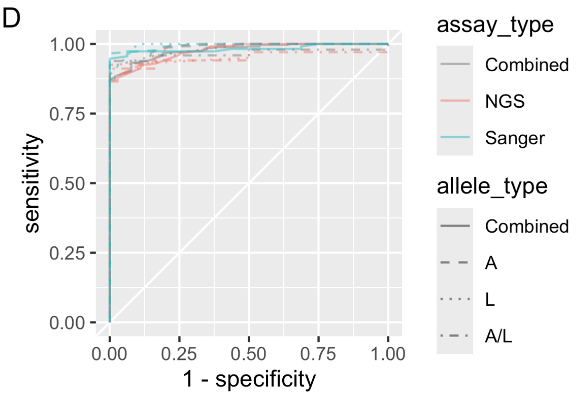


E

| assay_type | allele_type | n_pos | n_neg | AUC | AUC_CI | max_Gmean |
|---|---|---|---|---|---|---|
| Combined | Combined | 343 | 52 | 0.978 | 0.966-0.990 | 0.934 |
| NGS | Combined | 230 | 36 | 0.975 | 0.959-0.991 | 0.930 |
| Sanger | Combined | 113 | 16 | 0.983 | 0.963-1.000 | 0.973 |
| Combined | A | 245 | 34 | 0.984 | 0.971-0.997 | 0.935 |
| NGS | A | 156 | 28 | 0.983 | 0.968-0.998 | 0.934 |
| Sanger | A | 89 | 6 | 0.994 | 0.983-1.000 | 0.983 |
| Combined | L | 49 | 12 | 0.983 | 0.960-1.000 | 0.958 |
| NGS | L | 40 | 2 | 0.969 | 0.898-1.000 | 0.962 |
| Sanger | L | 9 | 10 | 0.994 | 0.979-1.000 | 0.949 |
| Combined | A/L | 49 | 6 | 0.966 | 0.919-1.000 | 0.969 |
| NGS | A/L | 34 | 6 | 0.951 | 0.884-1.000 | 0.955 |
| Sanger | A/L | 15 | 0 | NA | NA | NA |

**Figure S7:** median (Q2) methylation scores for 4A161 and 4AL assays in UNR IRB samples. Caption is as for Figure 10 but with Q2 score rather than e2.m1 score.

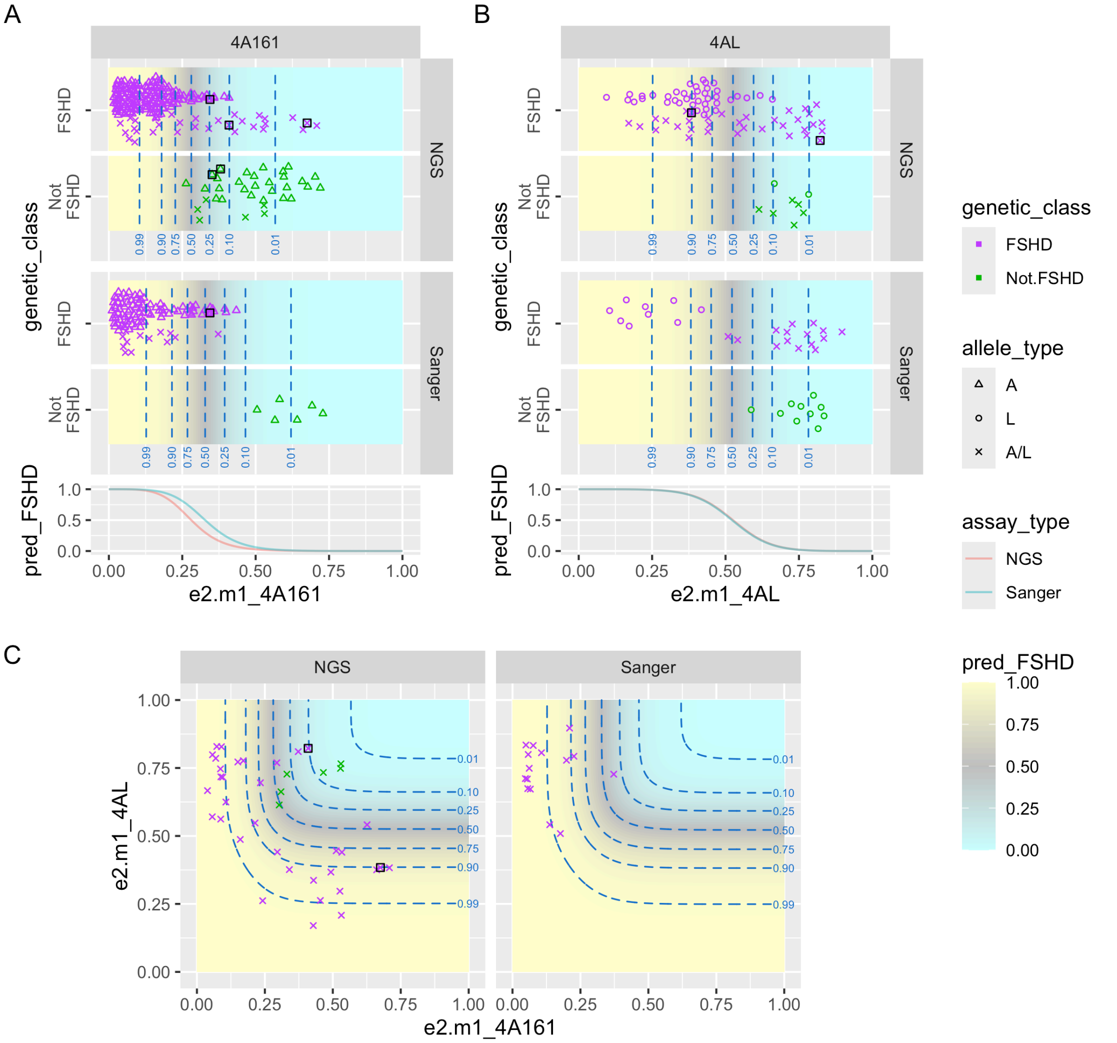


D

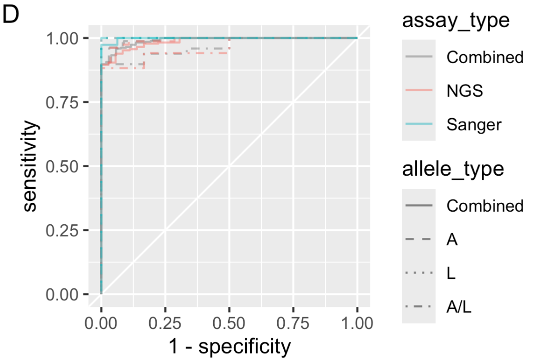


E

| assay_type | allele_type | n_pos | n_neg | AUC | AUC_CI | max_Gmean |
|---|---|---|---|---|---|---|
| Combined | Combined | 343 | 52 | 0.991 | 0.984-0.998 | 0.951 |
| NGS | Combined | 230 | 36 | 0.987 | 0.976-0.997 | 0.946 |
| Sanger | Combined | 113 | 16 | 0.998 | 0.995-1.000 | 0.987 |
| Combined | A | 245 | 34 | 0.994 | 0.987-1.000 | 0.967 |
| NGS | A | 156 | 28 | 0.992 | 0.983-1.000 | 0.963 |
| Sanger | A | 89 | 6 | 1.000 | 1.000-1.000 | 1.000 |
| Combined | L | 49 | 12 | 0.997 | 0.988-1.000 | 0.979 |
| NGS | L | 40 | 2 | 1.000 | 1.000-1.000 | 1.000 |
| Sanger | L | 9 | 10 | 1.000 | 1.000-1.000 | 1.000 |
| Combined | A/L | 49 | 6 | 0.966 | 0.920-1.000 | 0.948 |
| NGS | A/L | 34 | 6 | 0.961 | 0.904-1.000 | 0.939 |
| Sanger | A/L | 15 | 0 | NA | NA | NA |

**Figure S8:** e2.m1 methylation scores for 4A161 and 4AL assays in UNR IRB samples. Caption is as for Figure 10 but with predicted prob(FSHD) for samples with allele type A/L taken to be $P_{A/L} = 1 - (1 - P_A)(1 - P_L)$ rather than $\max(P_A, P_L)$ (see Methods).

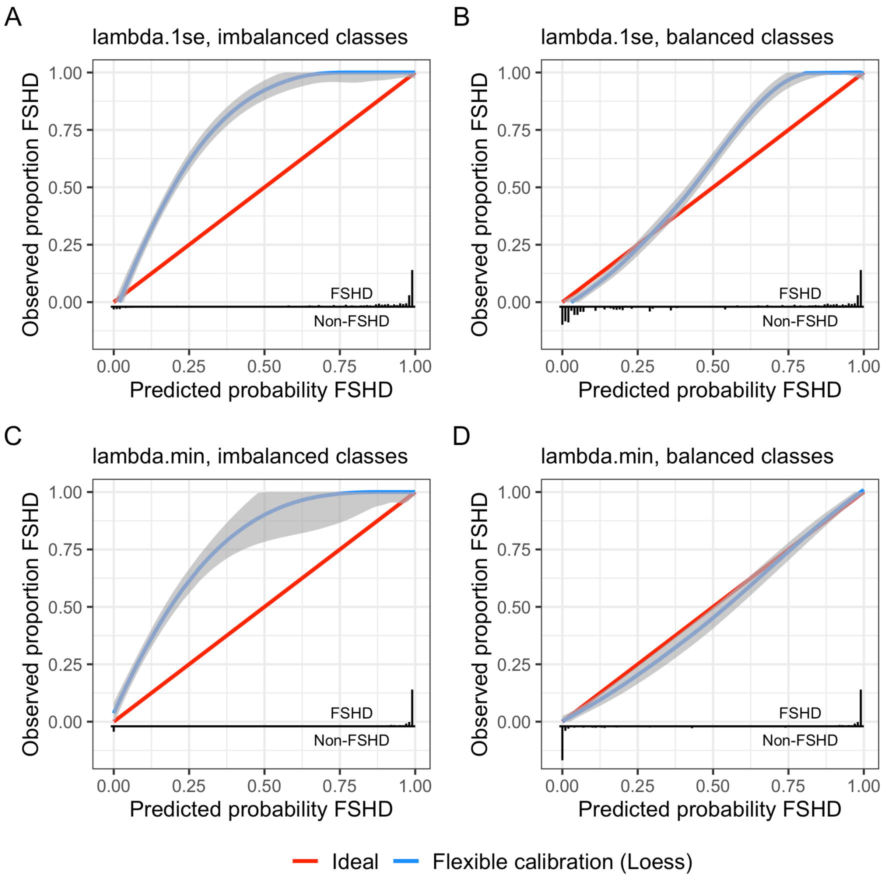


**Figure S9:** Calibration of predicted probabilities. Horizontal axes indicate predicted probabilities of subjects being genetic FSHD based on e2.m1 methylation values of 4A and/or 4AL alleles. Histograms of probabilities are shown in the lower margin (positive for FSHD and negative for non-FSHD). Vertical axes indicate observed proportions of subjects with FSHD. The blue curve shows the LOESS-smoothed relationship between predicted probabilities and observed proportions (with 95% confidence interval shaded gray), and the red diagonal line represents ideal calibration. Panel **A** shows calibration on 343 FSHD vs 52 non-FSHD samples in the UNR IRB study. Because the logistic regression models used an offset to adjust for the class imbalance, it is expected that predicted probabilities are lower than observed proportions in A. In panel **B**, non-FSHD samples were randomly up-sampled to balance the class sizes. Predicted probabilities are still conservative at both extremes, due in part to the use of a stronger elastic net regularization penalty (lambda.1se). Panels **C** and **D** are analogous to panels A and B but using a milder regularization penalty (lambda.min). Note that in this case very few samples had predicted probability between 0.1 and 0.9, so observed proportions in this range are not precisely determined. Calibration plots based on Q1 rather than e2.m1 are similar (not shown).

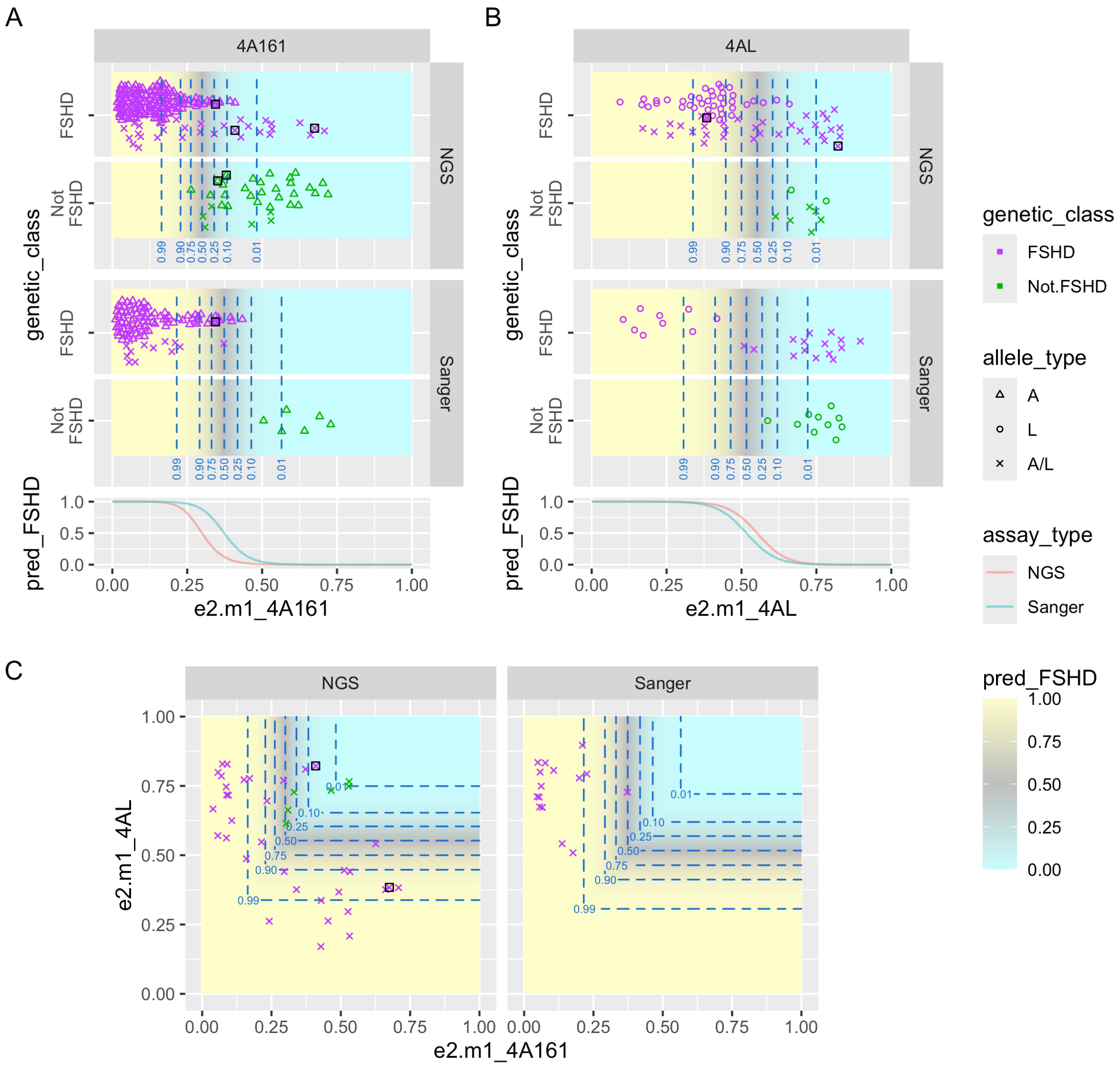


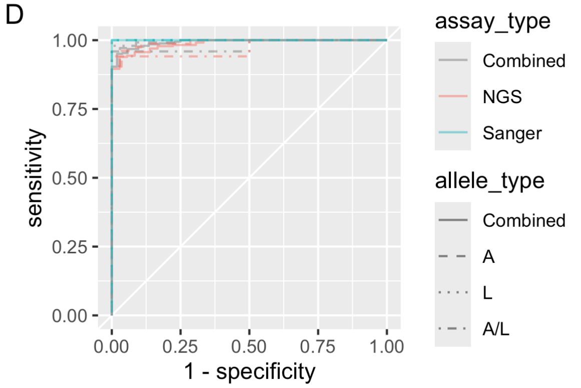


E

| assay_type | allele_type | n_pos | n_neg | AUC | AUC_CI | max_Gmean |
|---|---|---|---|---|---|---|
| Combined | Combined | 343 | 52 | 0.993 | 0.987-0.999 | 0.965 |
| NGS | Combined | 230 | 36 | 0.988 | 0.978-0.998 | 0.956 |
| Sanger | Combined | 113 | 16 | 1.000 | 1.000-1.000 | 1.000 |
| Combined | A | 245 | 34 | 0.995 | 0.988-1.000 | 0.971 |
| NGS | A | 156 | 28 | 0.992 | 0.983-1.000 | 0.963 |
| Sanger | A | 89 | 6 | 1.000 | 1.000-1.000 | 1.000 |
| Combined | L | 49 | 12 | 0.998 | 0.994-1.000 | 0.990 |
| NGS | L | 40 | 2 | 1.000 | 1.000-1.000 | 1.000 |
| Sanger | L | 9 | 10 | 1.000 | 1.000-1.000 | 1.000 |
| Combined | A/L | 49 | 6 | 0.980 | 0.946-1.000 | 0.979 |
| NGS | A/L | 34 | 6 | 0.971 | 0.923-1.000 | 0.970 |
| Sanger | A/L | 15 | 0 | NA | NA | NA |

**Figure S10:** e2.m1 methylation scores for 4A161 and 4AL assays in UNR IRB samples using milder regularization. Caption is as for Figure 10**.** The only difference is that here lambda.min rather than lambda.1se was used as the elastic net regularization penalty.

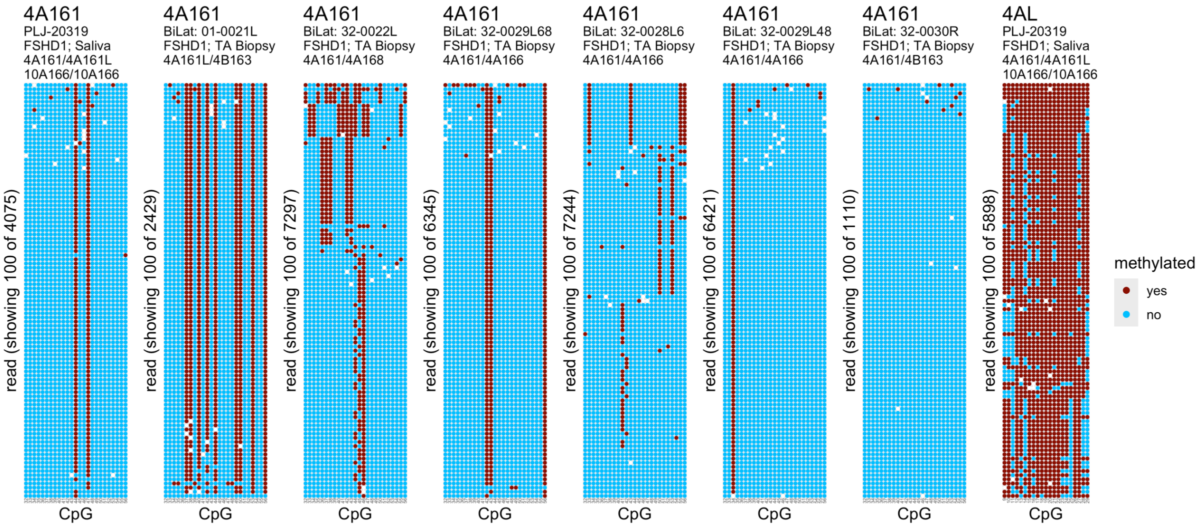


**Figure S11:** Methylation plot for NGS samples flagged as having potentially strong clonal artifacts. All are for 4A161 assays other than one 4AL assay at the far right. Assays with Q2 > 5% were flagged if they had Q2 – Q1 < 1% or Q3 – Q2 < 1%, and all assays (regardless of Q2) were flagged if they had pct90 – pct10 < 1%. Predicted haplotypes based on PCR (shown beneath sample names) can be helpful for distinguishing very low methylation of a present allele from artifactual signals for an absent allele.

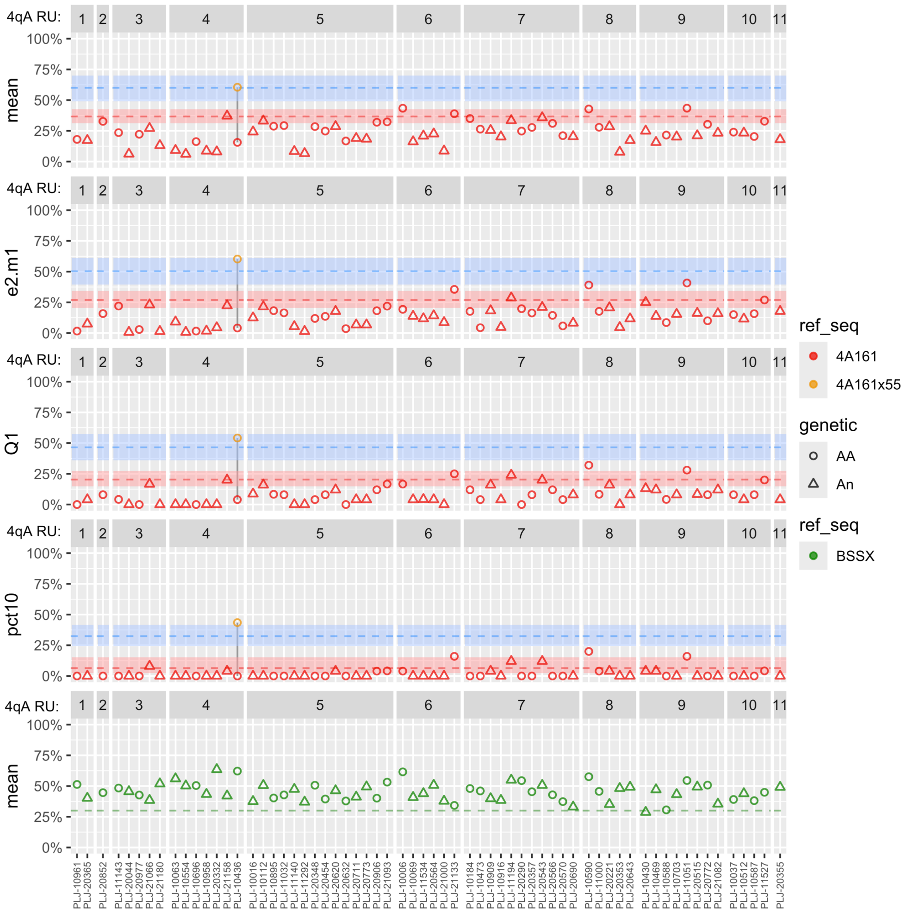


**Figure S12:** Number of D4Z4 repeat units (RUs) on contracted FSHD1-associated 4A alleles vs methylation of distal repeat. The samples and scores shown are the same as in Figure 11A but grouped by approximate number of RUs of contracted 4A allele, dropping samples for which this information was not provided. Symbols indicate the predicted number of 4A161-type alleles based on PCR of the distal repeat, either one (group An; triangle) or two (group AA; circle). Samples with a 4AL allele in additional to the 4A allele are excluded as it may not be known *a priori* which is the contracted pathogenic allele, but samples in group An may also have a 4B allele (***n***on-permissive) or 4A166-type distal repeat (not targeted by BSSA assay and typically ***n***on-pathogenic even if on contracted array). RUs were either directly reported in Table S1 or estimated from EcoRI or EcoRI/BlnI fragment lengths in kb using the conversion #RU = (kb – 5)/3.3 for the former, #RU = (kb – 2)/3.3 for the latter, and #RU = (kb – 3.5)/3.3 if it was not specified which fragment was measured. Fractional values are rounded down in the figure. All samples were reported as having genetic FSHD1, which is typically defined by 1-10 RU, but one subject here was reported as having approximate EcoRI/BlnI fragment length of ~40kb so may have ~11RU.
