## Supplemental Table 2 for "The D4Z4caster DNA methylation signature identifies individuals at epigenetic risk for developing facioscapulohumeral muscular dystrophy (FSHD)"

**Supplemental Table 2**: NGS BSS Primers

1438F-P1

5’-CCACTACGCCTCCGCTTTCCTCTCTATGGGCAGTCGGTGAT**GTTTTGTTGGAGGAGTTTTAGGA**

4ALF-P1

5’-CCACTACGCCTCCGCTTTCCTCTCTATGGGCAGTCGGTGAT**TTATTTATGAAGGGGTGGAGTTTGTT**

475F-P1

5’-CCACTACGCCTCCGCTTTCCTCTCTATGGGCAGTCGGTGAT**TTAGGAGGGAGGGAGGGAGGTAG**

3626R-BC-A

5-CCATCTCATCCCTGCGTGTCTCCGACTCAG-[BC]-**AACAAAAATATACTTTTAACCRCCAAAAA**

1036R-BC-A

5-CCATCTCATCCCTGCGTGTCTCCGACTCAG-[BC]-**AACACCRTACCRAACTTACACCCTT**

The P1 or A sequence is underlined, [the Ion Express barcode is inserted here], and **the gene-specific sequences for bisulfite-converted DUX4** **are in bold.** Ion Express barcode sequences are available through Ion Torrent Server.
