## Supplemental Table 3 for "The D4Z4caster DNA methylation signature identifies individuals at epigenetic risk for developing facioscapulohumeral muscular dystrophy (FSHD)"

**>4A161 (GenBank HM190177.1 region 2628..3298)**

GCTCTGCTGGAGGAGCTTTAGGACGCGGGGTTGGGACGGGGTCGGGTGGTTCGGGGCAGGGCGGTGGCCTCTCTTTCGCG

GGGAACACCTGGCTGGCTACGGAGGGGCGTGTCTCCGCCCCGCCCCCTCCACCGGGCTGACCGGCCTGGGATTCCTGCCT

TCTAGGTCTAGGCCCGGTGAGAGACTCCACACCGCGGAGAACTGCCATTCTTTCCTGGGCATCCCGGGGATCCCAGAGCC

GGCCCAGGTACCAGCAGGTGGGCCGCCTACTGCGCACGCGCGGGTTTGCGGGCAGCCGCCTGGGCTGTGGGAGCAGCCCG

GGCAGAGCTCTCCTGCCTCTCCACCAGCCCACCCCGCCGCCTGACCGCCCCCTCCCCACCCCCACCCCCCACCCCCGGAA

AA**CGCGTCGTCCCCTGGGCTGGGTGGAGACCCCCGTCCCGCGAAACACCGGGCCCCGCGCAGCGTCCGGGCCTGACACCG**

**CTCCGGCGGCTCGCCTCCTCTGCGCCCCCGCGCCACCGTCGCCCGCCCGCCCGGGCCCCTGCAGCCTCCCAGCTGCCAGC**

**GCG**GAGCTCCTGGCGGTCAAAAGCATACCTCTGTCTGTCTTTGCCCGCTTCCTGACTAGACCTGCGCGCAGTGCGCACCC

CGGCTGACGTGCAAGGGAGCTCGCTGGCCTC

**>4A161x55 (GenBank HM190177.1 region 2628..3298 with G_A at pos 561 of excerpt)**

GCTCTGCTGGAGGAGCTTTAGGACGCGGGGTTGGGACGGGGTCGGGTGGTTCGGGGCAGGGCGGTGGCCTCTCTTTCGCG

GGGAACACCTGGCTGGCTACGGAGGGGCGTGTCTCCGCCCCGCCCCCTCCACCGGGCTGACCGGCCTGGGATTCCTGCCT

TCTAGGTCTAGGCCCGGTGAGAGACTCCACACCGCGGAGAACTGCCATTCTTTCCTGGGCATCCCGGGGATCCCAGAGCC

GGCCCAGGTACCAGCAGGTGGGCCGCCTACTGCGCACGCGCGGGTTTGCGGGCAGCCGCCTGGGCTGTGGGAGCAGCCCG

GGCAGAGCTCTCCTGCCTCTCCACCAGCCCACCCCGCCGCCTGACCGCCCCCTCCCCACCCCCACCCCCCACCCCCGGAA

AA**CGCGTCGTCCCCTGGGCTGGGTGGAGACCCCCGTCCCGCGAAACACCGGGCCCCGCGCAGCGTCCGGGCCTGACACCG**

**CTCCGGCGGCTCGCCTCCTCTGCGCCCCCGCGCCACCGTCGCCCGCCCGCCCGGGCCCCTGCAGCCTCCCAGCTGCCAGC**

**ACG**GAGCTCCTGGCGGTCAAAAGCATACCTCTGTCTGTCTTTGCCCGCTTCCTGACTAGACCTGCGCGCAGTGCGCACCC

CGGCTGACGTGCAAGGGAGCTCGCTGGCCTC

**>4A166 (Genbank HM190195.1 region 2631..3301)**

GCTCTGCTGGAGGAGCTTTAGGACGCGGGGTTGGGACGGGGTCGGGTGGTTCGGGGCAGGGCGGTGGCCTCTCTTTCGCG

GGGAACACCTGGCTGGCTACGGAGGGGCGTGTCTCCGCCCCGCCCCCTCCACCGGGCTGACCGGCCTGGGATTCCTGCCT

TCTAGGTCTAGGCCCGGTGAGAGACTCCACACAGCGGAGAACTGCCATTCTTTCCTGGGCATCCCGGGGATCCCAGAGCC

GGCCCAGGTACCAGCAGGTGGGCCGCCTACTGCGCACGCGCGGGTTTGCGGGCAGCCGCCTGGGCTGTGGGAGCAGCCCG

GGCAGAGCTCTCCTGCCTCTCCACCAGCCCACCCCGCCGCCTGACCGCCCCCTCCCCACCCCCACCCCCCACCCCCGGAA

AA**CGCGTCGTCCCCTGGGCTGGGTGGAGACCCCCGTCCCGCGAAACACCGGGCCCCGCGCAGCGTCCGGGCCTGACACCG**

**CTCCGGCGGCTCGCCTCCTCTGCGCCCCCGCGCCACCGTCGCCCGCCCGCCCGGGCCCCTGCAGCCGCCCAGGTGCCAGC**

**ACG**GAGCGCCTGGCGGTCAAAAGCATACCTCTGTCTGTCTTTGCCCGCTTCCTGGCTAGACCTGCGTGCAGTGCGCACCC

CGGCTGACGTGCAAGGGAGCTCGCTGGCCTC

**>10A176T (GenBank HM190190.1 region 2638..3308)**

GCTCTGCTGGAGGAGCTTTAGGACGCGGGGTTGGGACGGGGTCGGGTGGTTCGGGGCAGGGCGGTGGCCTCTCTTTCGCG

GGGAACACCTGGCTGGCTACGGAGGGGCGTGTCTCCGCCCCGCCCCCTCCACCGGGCTGACCGGCCTGGGATTCCTGCCT

TCTAGGTCTAGGCCCGGTGAGAGACTCCACACCGCGGAGAACTGCCATTCTTTCCTGGGCATCCCGGGGATCCCAGAGCC

GGCCCAGGTACCAGCAGGTGGGCCGCCTACTGCGCACGCGCGGGTTTGCGGGCAGCCGCCTGGGCTGTGGGAGCAGCCCG

GGCAGAGCTCTCCTGCCTCTCCACCAGCCCACCCCGCCGCCTGACCGCCCCCTCCCCACCCCCACCCCCCGCCCCCGGAA

AA**CGCGTCGTCCCCTGGGCTGGGTGGAGACCCCCGTCCCGCGAAACACCGGGCCCCGCGCAGCGTCCGGGCCTGACACCG**

**CTCCGGCGGCTCGCCTCCTCTGCGCCCCCGCGCCACCGTCGCCCGCCCGCCCGGGCCCCTGCAGCCTCACAGCTGCCAGC**

**ACG**GAGCGCCTGGCGGTCAAAAGCATACCTCTGTCTGTCTTTGCCCGCTTCCTGGCTAGACCTGCGCGCAGTGCGCACCC

CGGCTGACGTGCAAGGGAGCTCGCTGGCCTC

**>4AL (GenBank MF422078.1 region 2469..2799)**

CCATTCATGAAGGGGTGGAGCCTGCCTGCCTGTGGGCCTTTACAAGGGCGGCTGGCTGGCTGGCTGGCTGTCCGGGCAGG

CCCCCTGGCTGCACCTGCCGCAGTGCACAGTCCGGCTGAGGTGCACGGGAGCCCGCCGGCCTCTCTCTGCCCG**CGTCCGT**

**CCGTGAAATTCCGGCCGGGGCTCACCGCGATGGCCCTCCCGACACCCTCGGACAGCACCCTCCCCGCGGAAGCCCGGGGA**

**CGAGGACGGCGACGGAGACTCGTTTGGACCCCGAGCCAAAGCGAGGCCCTGCGAGCCTGCAGCCTCCCAGCTGCCAGCG**C

GGAGCTCCTGG

**>4B168 (GenBank HM190161.1 region 3009..3190)**

GGGCTGAGGGCTGGGCCCACAGCCGC**CGCGCCGGCCGGCGGGGCACCACCCATTCGCCCCGGTTCCGGGGCCCAGGGAGT**

**GGGCGGTTTCCTCCGGGACAAAAGACCGGGACTCG**AGACTCCGTTCAATAAATGGTGCTGGGATAACTGGCTAGCCACAT

GCTGAAGATTGAACTGGGCCCC

**>BSSX (GenBank HM190174.1 region 32..597 with G_T at pos 153 of excerpt)**

TTAGGAGGGAGGGAGGGAGGCAGGGAGGCAGGGAGGAACGGAGGGAAAGACAGAGCGACGCAGGGACTGGGGGCGGGCGG

GAGGGAGCCGGGGACGGGGGGAGGAAGGCAGGGAGGAAAAGCGGTCCTCGGCCTCCGGGAGTAGCGGGACCCCCGCCCTC

CGGGAAAACGGTCAGCGTCCGGCGCGGGCTGAGGGCTGGGCCCACAGCCGCCGCGCCGGCCGGCGGGGCACCACCCATTC

GCCCCGGTTCCGTGGCCCAGGGAGTGGGCGGTTTCCTCCGGGACAAAAGACCGGGACTCGGGTTGC**CGTCGGGTCTTCAC**

**CCGCGCGGTTCACAGACCGCACATCCCCAGGC**TGAGCCCTGCAACGCGGCGCGAGGCCGACAGCCCCGGCCACGGAGGAG

CCACACGCAGGACGACGGAGGCGTGATTTTGGTTTCCGCGTGGCTTTGCCCTCCGCAAGGCGGCCTGTTGCTCACGTCTC

TCCGGCCCCCGAAAGGCTGGCCATGCCGACTGTTTGCTCCCGGAGCTCTGCGGGCACCCGGAAACATGCAGGGAAGGGTG

CAAGCC

**Supplemental Table 3:** Sequences of reference amplicons. Sequence ranges containing the CpGs used in scores for the NGS-based assays are in **bold**. DNA residues that vary among the first four sequences are highlighted (A=green, T=red, C=cyan, G=yellow). Note that 4A161x55 differs from 4A161 only in a G>A substitution in position 561, which disrupts the 55^th^ CpG in 4A161. Reads from predicted 4A159 haplotypes sometimes have this variant, and in such cases including 4A161x55 as a reference sequence allows reads from those haplotypes to be separated from reads from 4A161 haplotypes during the read mapping step, which is useful for subjects having both haplotypes. Some subjects with predicted 4A166 haplotypes have reads that map to the 4A161x55 reference sequence rather than the 4A166 reference sequence; both have A in position 561, and while there are several additional variants that differ between their reference sequences only one is in the high-coverage region shown in bold (T>G for 4A166 in position 547). The 4A166 reference sequence also has a C>A substitution in position 193 that disrupts the 16^th^ CpG in the 4A161, 4A166x55 and 10A176T references sequence; this is upstream of the high-coverage region in bold but still picked up in a small fraction of NGS reads and can be useful for interpreting results for reads that map to 4A161x55. *(Note: the 4A166 reference amplicon used for reported results also had 615G>A and 627T>C, with the former also in the 10A176T reference amplicon, but these positions are downstream of the regions being analyzed.)*
